# Common and distinct neurofunctional representations of core and social disgust in the brain: Coordinate-based and network meta-analyses

**DOI:** 10.1101/2021.09.07.459241

**Authors:** Xianyang Gan, Xinqi Zhou, Jialin Li, Guojuan Jiao, Xi Jiang, Bharat Biswal, Shuxia Yao, Benjamin Klugah-Brown, Benjamin Becker

**Affiliations:** The Clinical Hospital of Chengdu Brain Science Institute, School of Life Science and Technology, University of Electronic Science and Technology of China, Chengdu, Sichuan 610054, China; Max Planck School of Cognition, Leipzig 04103, Germany; Department of Biomedical Engineering, New Jersey Institute of Technology, New Jersey 7102, United States of America

**Author notes:** **Correspondence to** Benjamin Klugah-Brown or Benjamin Becker, University of Electronic Science and Technology, School of Life Science and Technology, No.2006, Xiyuan Ave, Chengdu, Sichuan, 611731, China.

**Keywords:** disgust, fMRI, meta-analysis, activation likelihood estimation (ALE), meta-analytic connectivity modeling (MACM), defensive-avoidance response, social cognition, face, insula, amygdala

## Abstract

Disgust represents a multifaceted defensive-avoidance response. On the behavioral level, the response includes withdrawal and a disgust-specific facial expression. While both serve the avoidance of pathogens, the latter additionally transmits social-communicative information. Given that common and distinct brain representation of the primary defensive-avoidance response (core disgust) and encoding of the social-communicative signal (social disgust) remain debated, we employed neuroimaging meta-analyses to (1) determine brain systems generally engaged in disgust processing, and (2) segregate common and distinct brain systems for core and social disgust. Disgust processing, in general, engaged a bilateral network encompassing the insula, amygdala, occipital and prefrontal regions. Core disgust evoked stronger reactivity in left-lateralized threat detection and defensive response network including amygdala, occipital and frontal regions while social disgust engaged a right-lateralized superior temporal-frontal network engaged in social cognition. Anterior insula, inferior frontal and fusiform regions were commonly engaged during core and social disgust, suggesting a common neural basis. We demonstrate a common and separable neural basis of primary disgust responses and encoding of associated social-communicative signals.

## 1. Introduction

Disgust represents a basic emotion and has been conceptualized as a multifaceted and evolutionary adaptive defensive-avoidance response. The disgust response may have initially evolved in the context of rejection of potential poisonous food (Darwin, 1965; Rozin and Fallon, 1987; Rozin et al., 2009) but the same underlying mechanism may have subsequently been adapted to facilitate avoidance of contaminations by pathogens and may even serve the avoidance of harmful social interactions (Curtis et al., 2011; Davey, 2011; Haidt et al., 1994; Kupfer and Fessler, 2018; Rozin and Fallon, 1987; Rozin et al., 2016; Vicario et al., 2017). In line with its evolutionary function, disgust is primarily elicited by exposure to potentially poisonous or infectious stimuli such as rotten food, cockroaches or individuals with poor personal hygiene as well as injury- and mutilation-related stimuli (Haidt et al., 1994; Olatunji and Sawchuk, 2005; Rozin and Fallon, 1987; Vicario et al., 2017). Exposure to these stimuli induces a distinct disgust reaction on different levels encompassing subjective experience of revulsion, as well as physiological and behavioral responses. The behavioral response includes rapid withdrawal from the eliciting stimulus as well as a distinct facial expression characterized by a wrinkled nose (Rozin et al., 1994) and raised upper lip (Rozin et al., 1994), reflecting the primary purpose of the disgust reaction in terms of avoiding or rejecting pathogens (Chapman et al., 2009; Oaten et al., 2009; Olatunji and Sawchuk, 2005; Rozin and Fallon, 1987; Rozin et al., 2009; Rozin et al., 1994; Schienle and Wabnegger, 2021) (in line with Vicario et al., (2017) henceforward referred to as *core disgust* processing). The highly distinct facial expression for disgust additionally serves social-communicative purposes (Chapman and Anderson, 2012; Chapman et al., 2009; Vicario et al., 2017; von dem Hagen et al., 2009) such that a disgusted facial expression signals to conspecifics a danger of being exposed to potential pathogens (Burklund et al., 2007; Darwin, 1965; Rozin et al., 1994; Surguladze et al., 2003; von dem Hagen et al., 2009) or an attitude of social rejection (Burklund et al., 2007; Surguladze et al., 2010; von dem Hagen et al., 2009). Accurate decoding of these signals is essential for survival and adaptive social interactions (Darwin, 1965; Ekman, 1993; Fusar-Poli et al., 2009b; Malhi et al., 2007; Phillips et al., 1999). In line with Vicario et al. (2017) who refer to the perception or appraisal of a disgusted facial expression in others as a social-communicative signal we henceforward refer to perception/appraisal of disgusted faces as *social disgust* processing. Of note, we thus specifically consider the perception of disgusted faces in others as *social disgust*, while a broader definition may also encompass other social signals of disgust, such as words, or the production of a disgusted facial expression. Although both, core and social disgust, share the essential function of a defensive-avoidance reaction towards potential pathogen exposure, the facial expression of disgust as social signal represents a more indirect and context depended signal which requires further interpretation by the observer (for similar reasoning with respect to other facial expression domains see e.g., Hadjistavropoulos et al., 2011; Zhou et al., 2020).

Initial evidence for common and separable neurobiological systems underlying core and social disgust responses additionally comes from human lesion models reporting that a patient with focal anterior insula and putamen lesions exhibited intact core disgust reactivity but exhibited selective impairments in recognizing disgust from facial signals (Calder et al., 2000) while a patient with selective lesions of the anterior insula exhibited both, dysregulated core disgust reactivity as well as altered production of disgust expressions (Cantone et al., 2019). A recent study in a larger sample of lesion patients confirmed core as well as social disgust processing impairments in patients with insula-basal ganglia lesions and additionally reported that patients with left hemispheric lesions exhibited pronounced impairments, suggesting a lateralization of disgust processes (Holtmann et al., 2020).

Indirect evidence for common and separable neurobiological systems comes from numerous functional imaging, particularly functional MRI, studies that examined the neural substrates of disgust processing in healthy subjects (the first functional MRI study on disgust was published in 1997) (Phillips et al., 1997). These studies demonstrated that visual, gustatory (Corradi-Dell’Acqua et al., 2016; Jabbi et al., 2008), olfactory (Berlin et al., 2017; Heining et al., 2004; Meier et al., 2015; Wicker et al., 2003), and auditory (Phillips et al., 1998; Zimmer et al., 2016) stimuli can induce the subjective experience of disgust and neural reactivity in the domains of core and social disgust reactivity.

Visual stimuli, such as pictures and film clips depicting core disgust elicitors (e.g., rotten food) or social disgust stimuli (i.e., disgusted faces) in combination with corresponding non-disgust related stimuli have been widely used in separate studies on core or social disgust. However, despite the large number of original studies the key regions involved in disgust in general, as well as core and social disgust in particular, remain strongly debated. For instance, while the insula has been considered and often reported as a key region for general disgust processing (Calder et al., 2001; Kirby and Robinson, 2017; Murphy et al., 2003; Phan et al., 2002; Vytal and Hamann, 2010) and insula activity has been reported during core (Schäfer et al., 2009; Schäfer et al., 2005; Schienle et al., 2020; Schienle et al., 2002; Shapira et al., 2003; Stark et al., 2007) as well as social disgust processing paradigms (Ashworth et al., 2011; Phillips et al., 1997; Wicker et al., 2003), several studies examining disgust processes did not observe insula activity (Deeley et al., 2008; Gorno-Tempini et al., 2001; Schienle et al., 2006; Schienle et al., 2005b; Stark et al., 2005; Stark et al., 2004; Stark et al., 2003; Trautmann et al., 2009), which may reflect a general involvement of the insula in interoceptive processes rather than a disgust specific engagement (Li et al., 2019; Stark et al., 2005; Yao et al., 2018a).

Moreover, several studies suggest a role of the amygdala in disgust processing, probably related to its role in early detection of threat and initiation of defensive responses (e.g., Becker et al., 2012; Zhou et al., 2021). Amygdala activity in response to core disgust stimuli (Pujol et al., 2018; Schienle et al., 2009; Schienle et al., 2006; Schienle et al., 2002; Schienle et al., 2014; Stark et al., 2005; Stark et al., 2003; Stark et al., 2007) as well as social disgust stimuli (Burklund et al., 2007; Fitzgerald et al., 2006; Phillips et al., 1998; Winston et al., 2003) has been reported in several studies, yet a considerable number of studies on the neural basis of disgust failed to find an engagement of the amygdala (Caseras et al., 2007; Deeley et al., 2007; Phillips et al., 2000; Phillips et al., 2004; Phillips et al., 1999; Phillips et al., 1997; Schienle and Scharmüller, 2013; Schroeder et al., 2005; Sprengelmeyer et al., 1998; Surguladze et al., 2010; Wicker et al., 2003). Finally, two studies included different visual disgust stimuli (i.e., core disgust and social disgust stimuli), yet surprisingly did not find robust social disgust related activity or common neural activity during social and core disgust processing, respectively (Schäfer et al., 2005; Schienle et al., 2020).

The ongoing debates about the brain systems generally engaged in disgust processing as well as during core or social disgust processing separately might be partly rooted in the limitations of single fMRI studies and particularly factors such as low and underpowered sample sizes, variations in processing depths (i.e., implicit vs explicit processing), stimuli sets, experimental designs (e.g., block design vs event-related design), study-specific samples with variations in cultural background or disgust propensity, field strength (e.g., 1T vs 3T) and methods of data analysis may have considerably contributed to the lack of robustly identified neural systems for disgust. The development and application of neuroimaging based meta-analyses have allowed to overcome these limitations by capitalizing on the results from previously published studies (Eickhoff et al., 2012; Laird et al., 2005; Wager et al., 2007), thus allowing to determine the convergent and robust neurofunctional basis of emotional and cognitive processes as well as to determine common and distinct neurofunctional underpinnings between processes (e.g., Chen et al., 2018; Feng et al., 2018). Previous studies have capitalized on neuroimaging meta-analyses to successfully determine common and separable brain representations of core emotions or emotional face expressions (Fusar-Poli et al., 2009b; Kirby and Robinson, 2017; Murphy et al., 2003; Phan et al., 2002; Vytal and Hamann, 2010).

By capitalizing on neuroimaging meta-analyses, we aimed at determining brain systems that are robustly engaged in general disgust processing, as well as core and social disgust processing in healthy subjects. Moreover, we aimed for the first time to systematically determine common and separable systems involved in core and social disgust processing which – in a conceptual context – may allow to (1) determine embodied simulation processes in the domain of disgust, with the embodied simulation theories proposing that overlapping cognitive and brain mechanisms are key to embodied simulation (Gallese and Sinigaglia, 2011, 2012) which in turn critically supports understanding of emotions, cognitions and intentions of other persons, and (2) may help to further segregate regions which facilitate survival relevant rapid evaluation of disgust-related features from regions that are involved in other domains such as e.g., face processing. Moreover, disgust has received increasing attention as a behavioral domain involved in the pathogenesis and maintenance of a number of psychiatric and neurological disorders with increasing evidence for common as well as core and social disgust-specific dysregulations in specific diagnostic entities (Vicario et al., 2017). A clearer understanding of the common and separable neural systems of disgust processing in healthy populations could thus provide a basis for further delineating transdiagnostic and disorder-specific dysregulations in these domains.

Based on overarching disgust frameworks, we hypothesized that disgust-related processing will generally recruit regions engaged in motivational aspects, defensive responses and interoception such as the insula and amygdala, and - considering the role of prefrontal cortex as an integration hub for affective and cognitive functions (Kirby and Robinson, 2017) and its activation during disgust processing (Baumann and Mattingley, 2012; Phillips et al., 1997; Tettamanti et al., 2012; Wicker et al., 2003) - we expected this region to be involved during general disgust processing. Given that previous studies reported that occipital regions showed a strong engagement during the processing of negatively valenced emotional visual stimuli (Paradiso et al., 1997; Radua et al., 2014; Stark et al., 2007; Tao et al., 2021) we expected that these regions would be activated during general disgust processing. Based on the previous literature, we further expected that common and separable (domain-specific) networks would underlie core and social disgust processes, such that the insula, together with orbitofrontal/inferior frontal regions - proposed to serve as a common endpoint of distributed networks engaged in emotion processing (Sprengelmeyer et al., 1998) and repeatedly observed during core (Pujol et al., 2018; Schienle and Scharmüller, 2013; Stark et al., 2007; Tettamanti et al., 2012) as well as social disgust processing (Chakrabarti et al., 2006; Reidy et al., 2016; Sprengelmeyer et al., 1998; von dem Hagen et al., 2009) - would be recruited across domains; while regions involved in social processing, such as the face processing network and social cognition network including e.g., superior temporal and dorsomedial prefrontal regions (Fusar-Poli et al., 2009b; Liu et al., 2021; Sabatinelli et al., 2011), would be differentially engaged.

## 2. Methods

### 2.1. Literature search

The present meta-analytic review was conducted in line with the PRISMA guidelines for systematic reviews (Page et al., 2021) and the meta-analytic protocols were preregistered on the Open Science Framework (https://osf.io/pu59t).

Articles published prior to May 2021 were identified using the following keywords: (“functional magnetic resonance imaging” OR “fMRI”) AND (“disgust facial expression” OR “disgusted facial expression” OR “disgust face” OR “disgusted face” OR “facial motion, disgust” OR “disgust elicitors” OR “disgusting elicitors” OR “disgust stimuli” OR “disgusting stimuli” OR “disgust visual scenes” OR “disgusting visual scenes” OR “disgust”) on the PubMed and Web of Science databases. The search was complemented by a corresponding search on the BrainMap functional database (May 2021; containing 3406 papers and 16901 experiments) using the Sleuth software (version 3.0.4, https://www.brainmap.org/sleuth/) with the following search terms: “Experiments: Context, Normal Mapping”, “Experiments: Activations, Activations only”, “Experiments: Imaging Modality, fMRI” and “Experiments: Behavioral Domain, Emotion, Negative-Disgust”. Finally, reference lists of included articles were searched for relevant articles not identified during the initial database search. Noteworthy, in line with our working definition of social disgust, studies for the social disgust analyses were limited to studies examining the perception/appraisal of disgusted facial expressions in others. Following removal of duplicates, 419 neuroimaging papers on disgust processing were retained.

### 2.2. Inclusion and exclusion criteria

To identify suitable publications according to our criteria, titles and abstracts of identified papers were initially examined by two independent reviewers (X. G. and X. Z.). Full texts of eligible papers were downloaded for detailed examination. Any disagreement between these two reviewers was resolved by discussion and a final decision by a third independent reviewer (J. L.). Papers that passed the screening process underwent a thorough and detailed evaluation of eligibility (B. B. and X. G.), studies with ambiguous experimental contrasts were removed at this stage and disagreements were further resolved by another reviewer (B. K-B.). A flowchart illustrating the detailed article screening and exclusion process is shown in **Fig. 1**. The following inclusion criteria were applied: (1) studies reporting neuroimaging experiments on disgust processing which presented static pictures or film clips displaying clearly disgust-related contents (e.g., disgusted facial expressions and disgusting visual scenes; we limited the stimuli only to the visual modality for two reasons: one is to control for modality when comparing core and social disgust, the other reason is that the number of studies using olfactory, gustatory and auditory stimuli or script-induced imagination was too low to warrant a robust modality-specific meta-analyses); (2) results were published as original papers in peer-reviewed English-language journals; (3) contrasts for healthy subjects were reported (i.e., healthy volunteers only). Studies reporting only spatial coordinates across or between healthy controls and clinical groups were excluded, but included if data was separately reported for the healthy control group; (4) task-related fMRI was employed; (5) activation contrasts generated by the subtraction approach represent main effects of disgust relative to a baseline or control condition (e.g., disgust faces > neutral faces, disgust elicitors > fixation); (6) activation in terms of higher activity for the disgust-related stimuli reported (deactivations were not included because the underlying functional process remains difficult to interpret in fMRI studies and because the current version of the ALE software can’t differentiate activations from deactivations, similar approach see e.g., Vytal and Hamann, 2010); (7) results reported in standard stereotactic coordinates (either Talairach or Montreal Neurological Institute [MNI] space); (8) results describing associations between disgust-related neural activity and behavioral ratings of disgust, psychophysiological responses or personality and symptom dimensions were not included; (9) other types of emotional faces including schematic faces were excluded; (10) whole-brain results were reported (we excluded studies reporting only region-of-interest, ROI, analyses); (11) for studies reporting multiple but not independent contrasts, only the contrast that substantially reflected the process of interest was included. For instance, if one study reported two contrasts employed disgust stimuli with different intensity levels, only the results corresponding to higher disgust intensity were included (similar approach see Tao et al., 2021); (12) for studies that included multiple relevant and independent contrasts, all such contrasts were included (similar approach see Tao et al., 2021).

**Fig. 1.**
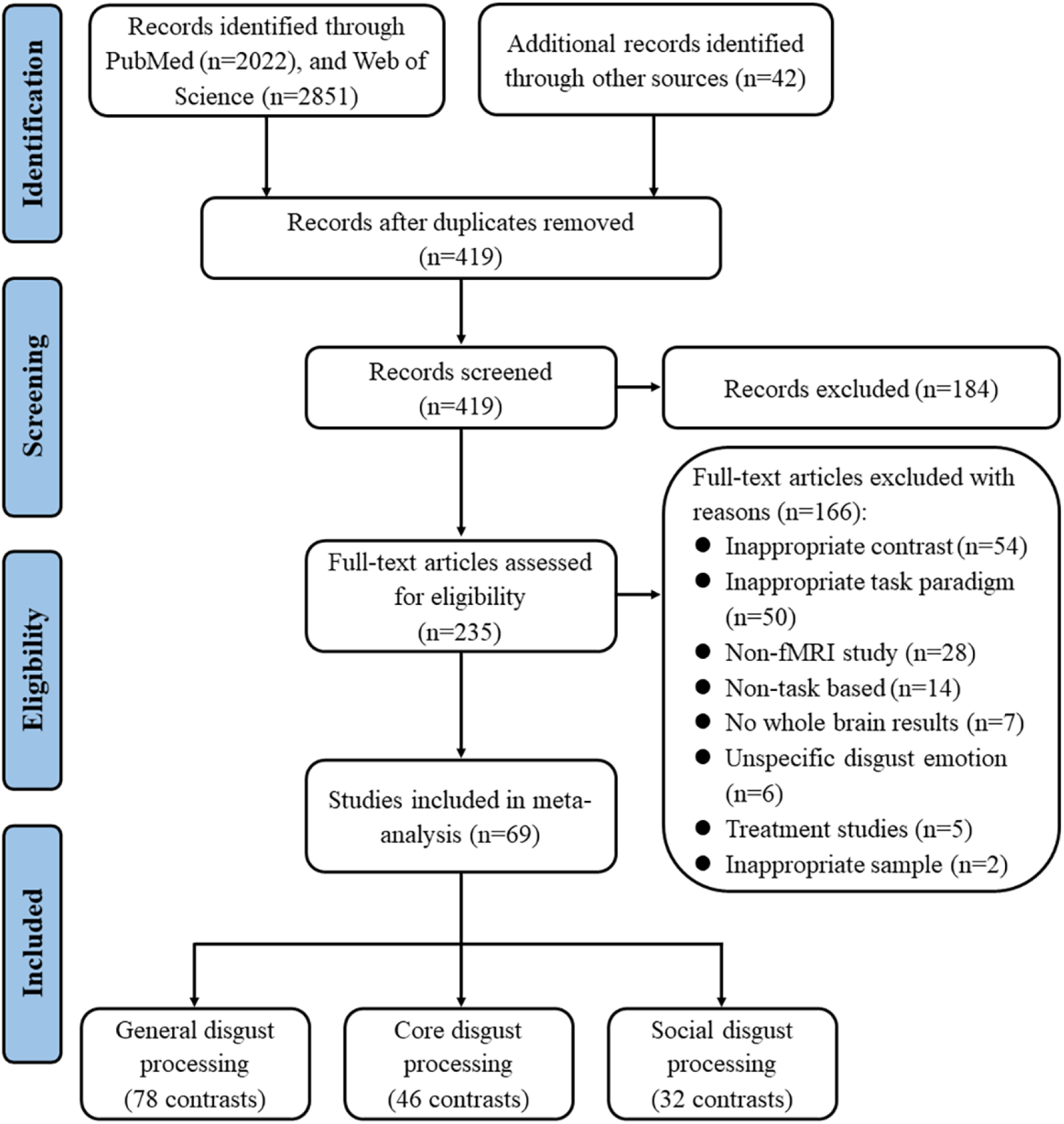
Flowchart illustrating the article screening, exclusion and inclusion process according to PRISMA procedure.

To reduce the possibility of biased study sampling, we contacted the corresponding authors of 11 studies for which Talairach or MNI coordinates for the appropriate contrast were not reported (e.g., for healthy controls in a case-control neuroimaging study).

After the in-depth screening procedure, a total of 69 papers (**Supplementary Table 1**) were included in the meta-analyses, and according to the aims of the analyses organized as follows: (1) general disgust processing: 69 papers, 71 experiments, 717 foci, 78 contrasts, 1409 subjects; (2) core disgust processing: 39 papers, 41experiments, 402 foci, 46 contrasts, 897 subjects; (3) social disgust processing: 30 papers, 30 experiments, 315 foci, 32 contrasts, 512 subjects. Based on recent recommendations for activation likelihood estimation (ALE) meta-analyses suggesting that at least 20 (and 17 in worst case) experiments are sufficient to determine moderate effects with sufficient statistical power (80%) (Eickhoff et al., 2016), this database allowed us to draw robust inferences.

### 2.3. Activation likelihood estimation

We performed a series of coordinate-based meta-analyses using the current version of GingerALE 3.0.2 (Eickhoff et al., 2012; Eickhoff et al., 2011; Eickhoff et al., 2009) available via http://www.brainmap.org/ale/. All coordinates (either in Talairach or MNI space) and sample size were manually extracted from the original articles. The coordinates in Talairach space were converted to MNI space using the Lancaster transform algorithm, while Talairach space coordinates transformed from MNI space using the Brett transformation were converted back to MNI space using the Brett transformation (Laird et al., 2010; Lancaster et al., 2007). Briefly, the ALE method estimates the likelihood of each voxel being activated under multiple independent experiments with consideration of participants in each experiment. In other words, ALE aims to find the most robust convergent foci between a number of included experiments. To address spatial uncertainty, ALE treats activation foci from single studies as 3D Gaussian probability distributions. Modeled activation (MA) maps for each study were generated by combining the probabilities of all activation foci for each voxel (Eickhoff et al., 2009; Turkeltaub et al., 2012). The union of these generated MA maps was computed while taking into account the disparities between true activation and noise to determine a voxel-wise ALE score (Klugah-Brown et al., 2020). These ALE scores use the convergence of whole-brain coordinates from experiments under a non-random spatial distribution to create clusters formed by equal probabilities of voxels (Eickhoff et al., 2012; Müller et al., 2018). The ALE map reflects clusters corresponding to the location of convergence. And the resulting non-parametric p values account for the proportion of the random spatial relation between the various experiments under the null distribution (Klugah-Brown et al., 2021; Kohn et al., 2014).

For single dataset analyses (i.e., general disgust processing, core disgust processing, and social disgust, respectively), the ALE statistics were calculated at each voxel, and cluster-level family-wise error correction (cluster-FWE) (Eickhoff et al., 2016) was applied to the ALE results to control for multiple comparisons. The voxel-level threshold was set at p < 0.001 (uncorrected cluster forming), and the cluster-level threshold was set at p < 0.05 (Eickhoff et al., 2017; Eickhoff et al., 2016; Müller et al., 2018), p values in our analyses were generated by 5,000 permutations (Laird et al., 2010; Tao et al., 2021).

### 2.4. Contrast and conjunction analyses

The present meta-analysis aimed at determining regions robustly involved in disgust processing across domains as well as to differentiate regions that are convergent and divergent between core and social disgust processing in healthy subjects. To this end we employed a series of contrast and conjunction analyses (comparing the meta-analytic maps for core and social disgust processing) according to the following steps: (1) single dataset analyses were performed separately for core disgust processing and social disgust processing; (2) the two single datasets to be compared were pooled together using the “Merge & Save Foci” function; (3) a single dataset analysis was conducted on the pooled foci file employing identical statistical criteria as for the preceding single dataset analyses; (4) via the separate single dataset analysis three corrected ALE images were obtained for general disgust processing, core disgust processing, and social disgust processing, respectively. Then, we further determined common and distinct brain regions by conducting contrast and conjunction analyses by subjecting these images into GingerALE toolbox to obtain the similarities as well as differences in the neurobiological basis of core and social disgust processing. Conjunction analyses were performed to determine common regions across core and social disgust processing which were considered as brain systems involved in disgust across processing domains (domain-general). While contrast analyses were implemented to determine brain regions differentially activated for core and social disgust processing (domain-specific) (See **Supplementary Materials** for detailed methodology of conjunction and contrast analyses).

In accordance with previous studies employing contrast and conjunction analyses (Alain et al., 2018; Papitto et al., 2020), the ALE images were thresholded at p < 0.05 (uncorrected), with minimum cluster size of 100 mm^3^ and 10,000 permutations, thereby resulting in a larger number of permutations as compared to single dataset analyses (Alain et al., 2018; Papitto et al., 2020; Wertheim and Ragni, 2018; Yu et al., 2018). In general, the rationale was to combine a strict threshold for the determination of the general networks examined in the single dataset analyses with a more sensitive threshold for examining common and separable neural bases in the conjunction and contrast analyses. Moreover, we applied a higher number of permutations to the conjunction and contrast analyses to determine a more accurate number of voxels reflecting the overlapping neural basis of the different disgust processes (i.e., social and core disgust). Results based on an identical number of permutations revealed similar results and are additionally presented in the supplements.

### 2.5. Meta-analytic connectivity modeling (MACM) analyses

To further functionally characterize the domain-general disgust regions on the network level MACM was employed. Briefly, MACM examines whole-brain functional connectivity patterns by identifying the above-chance covariance between two or more brain regions (Laird et al., 2009; Robinson et al., 2012; Robinson et al., 2010). This approach is similar to the functional connectivity patterns observed in resting-state fMRI (Kotkowski et al., 2018; Reid et al., 2017; Smith et al., 2009), yet further complements the resting-state functional connectivity approach as it provides a measure of functional connectivity during a range of task-associated mental states (Langner et al., 2014). Coordinates of the peak activation clusters (i.e., right inferior frontal gyrus and left fusiform gyrus) from the conjunction analyses were selected, with the right inferior frontal gyrus located at (44, 26, -8) and the left fusiform gyrus located at (−42, -50, -14). For the fusiform gyrus clusters, we focused on the peak with the higher ALE value. For the MACM analyses, the regions were defined in MNI space using a 10 mm sphere centered at the peak coordinates using the SPM toolbox MarsBaR (http://marsbar.sourceforge.net). A standard MNI brain template (Colin27_T1_seg_MNI.nii; https://www.brainmap.org/ale/Colin27_T1_seg_MNI.nii.gz) was used to visualize both ROI masks. Next region-specific functional connectivity was employed using the functional database of BrainMap’s Sleuth 3.0.4 software. To increase the robustness, we employed two convergent approaches (labeled as MACM-A and MACM-B, respectively) which enable us to determine the functional network characterization by determining co-activated regions in previous experiments. Specifically, we used two different search terms when conducting these two separate MACM analyses in Sleuth database. The search terms for “MACM-A” incorporating (1) “Locations: MNI Image” used to upload the corresponding 10 mm spherical ROI in MNI space, (2) “Experiments: Behavioral Domain, Emotion, Negative-Disgust”. In this way, we can get the co-activation patterns engaged in disgust processing, and this approach has been adopted by one recent published paper (Tao et al., 2021). The search terms for “MACM-B” are: (1) “Locations: MNI Image” used to upload the corresponding 10 mm spherical ROI in MNI space, (2) “Experiments: Activations, Activations only”, (3) “Experiments: Context, Normal Mapping”, (4) “Subjects: Diagnosis, Normals”. This approach has been recommended by many researchers (Meier et al., 2021; Papitto et al., 2020), including the user manual for GingerALE (https://www.brainmap.org/ale/manual.pdf). This approach enables the investigation of brain region covariance above chance within a given seed ROI across a large and different pool of neuroimaging experiments from the functional database. After each of these separate searching approaches, coordinates from downloaded experiments meeting the criteria were automatically exported as a text file and converted into MNI space.

The composition of the two datasets for MACM-A is as follows: (1) the ROI in right inferior frontal gyrus (73 experiments, 891 subjects, 719 foci), (2) the ROI in left fusiform gyrus (28 experiments, 414 subjects, 163 foci). And the composition of the two datasets for MACM-B is the following: (1) the ROI in right inferior frontal gyrus (473 experiments, 7,768 subjects, 7,195 foci), (2) the ROI in left fusiform gyrus (366 experiments, 5,439 subjects, 5,801 foci). Finally, these new datasets were analyzed separately using GingerALE 3.0.2 with the same statistical criteria previously applied in the single dataset analyses (voxel-level of p < 0.001 (uncorrected), cluster-FWE of p < 0.05, 5,000 permutations).

### 2.6. Functional characterization of ALE activations on the behavioral level

To facilitate a robust characterization of the identified regions on the behavioral level a meta-analytic decoding approach was employed. To this end, we capitalized on the Brain Annotation Toolbox (BAT, available at https://istbi.fudan.edu.cn/lnen/info/1173/1788.htm) (Liu et al., 2019), which is based on the activation maps in MNI space for the 217 functional terms from the Neurosynth database (http://neurosynth.org/) (Yarkoni et al., 2011). The corresponding functional annotation analysis hypothesizes that voxels within a given cluster/region are more likely to be co-activated by the same terms that are functionally connected with the cluster/region, compared with voxels selected randomly, this way BAT enables the functional characterization of a given region based on the meta-analytically associated terms (Liu et al., 2019). In the present study, we employed this approach to characterize the domain-general disgust (core ∩ social; where ∩ represents intersection) as well as core disgust and social disgust regions. P < 0.05 was considered statistically significant based on 10,000 permutations. For further details, please turn to the BAT website (https://istbi.fudan.edu.cn/lnen/softwares/BAT_User_Manual_V1.1.docx).

### 2.7. Report and visualization of results

For the results presented in tabular overview, the following indices were reported: volume of clusters, lateralization of brain regions, labels of brain regions, peak coordinates from voxels, ALE/Z value (Tao et al., 2021). For the conjunction and contrast results, information of percentage of overlap was further reported. With respect to the statistical indices ALE and Z values are provided for single dataset analyses; while ALE values are provided for conjunction and Z values are provided for contrast analyses (see Papitto et al., (2020) for details regarding the underlying inferential statistic approach). For visualization, the ALE results were displayed using the current version of Mango 4.1 (https://rii.uthscsa.edu/mango/), overlaid onto a standard MNI brain template (Colin27_T1_seg_MNI.nii).

## 3. Results

### 3.1. General disgust processing

The ALE meta-analysis encompassing all disgust processing studies revealed twelve significant clusters (**Table 1, Fig. 2a**) that were robustly engaged in general disgust processing. The clusters were located in the bilateral insula, amygdala spreading into the right globus pallidus, fusiform gyrus spreading into the declive, middle occipital gyrus, right inferior occipital gyrus, right lingual gyrus, left inferior temporal gyrus, left inferior parietal lobule, right inferior frontal gyrus, and left superior frontal gyrus. See **Supplementary Materials** for a detailed presentation of percentage of overlap with atlas-based brain regions.

**Table 1.**
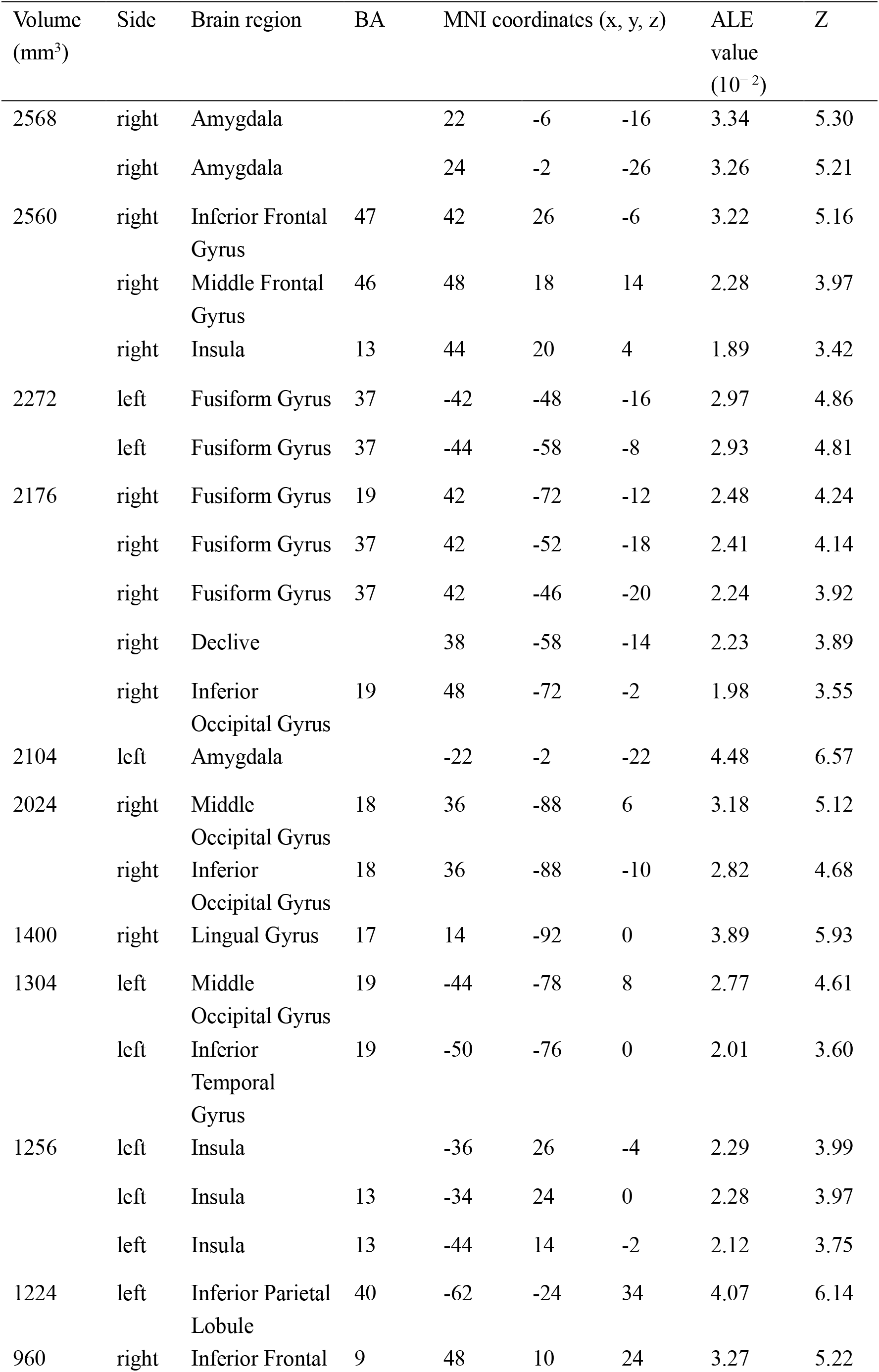
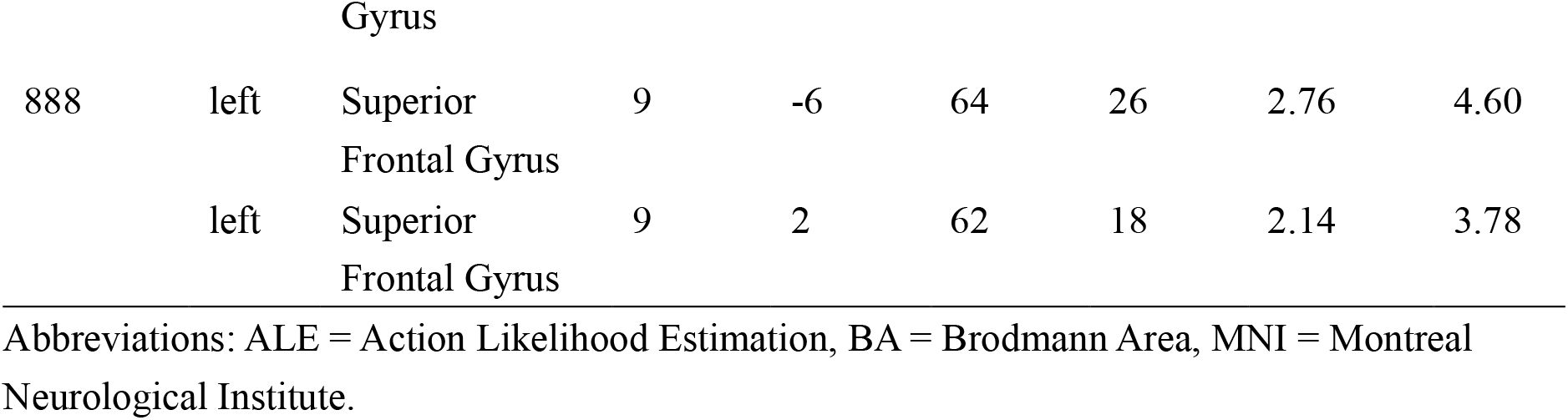
Brain regions generally activated by all studies of disgust processing in healthy subjects.

**Fig. 2.**
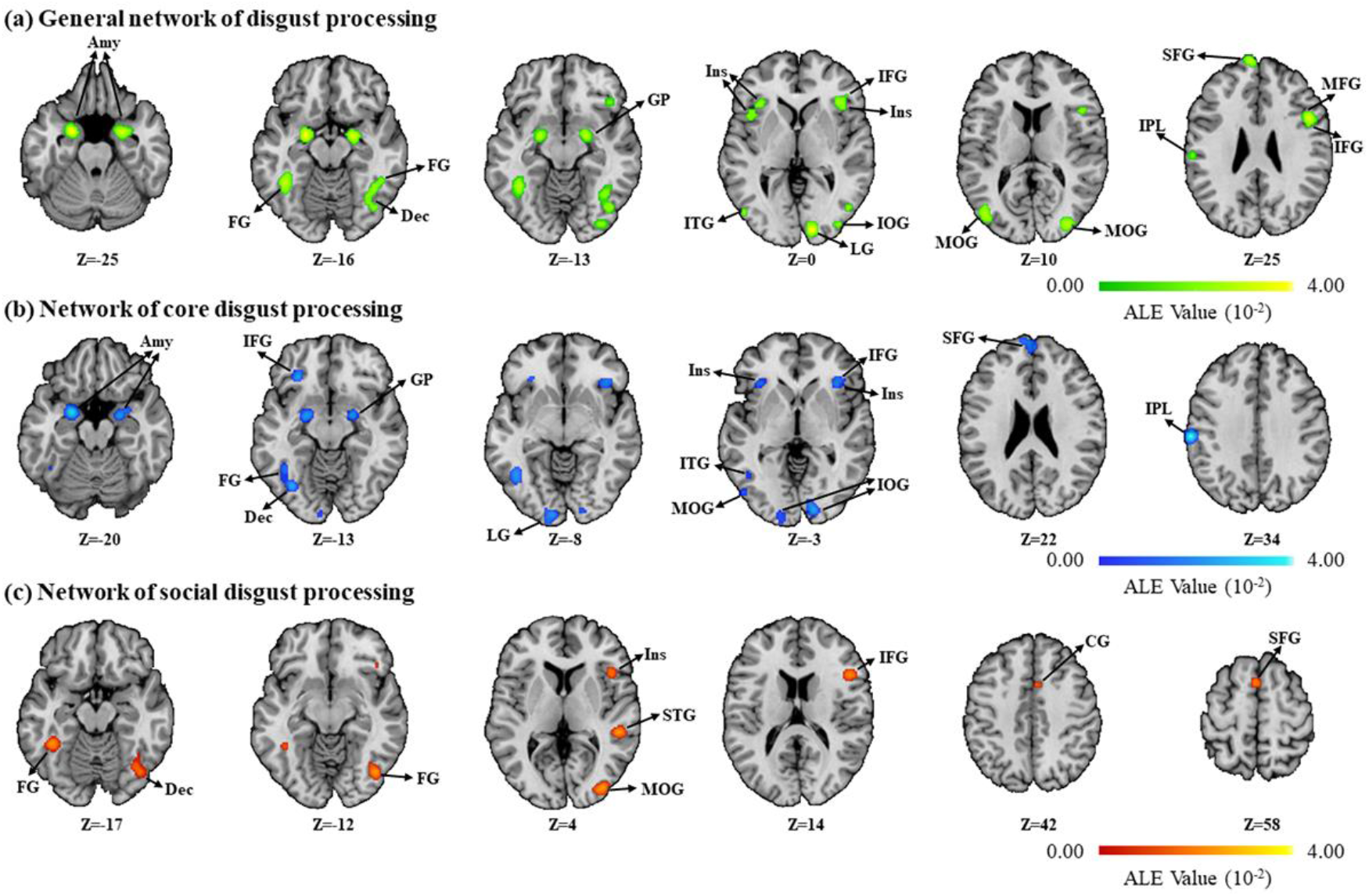
Overview of significant clusters from the single dataset analyses. **(a)** General network of disgust processing in healthy subjects. **(b)** Network of core disgust processing in healthy subjects. **(c)** Network of social disgust processing in healthy subjects. Coordinates are in the MNI space. Abbreviations: Amy = Amygdala, CG = Cingulate Gyrus, Dec = Declive, FG = Fusiform Gyrus, GP = Globus Pallidus, IFG = Inferior Frontal Gyrus, Ins = Insula, IOG = Inferior Occipital Gyrus, IPL = Inferior Parietal Lobule, ITG = Inferior Temporal Gyrus, LG = Lingual Gyrus, MFG = Middle Frontal Gyrus, MOG = Middle Occipital Gyrus, SFG = Superior Frontal Gyrus, STG = Superior Temporal Gyrus.

### 3.2. Core disgust processing

Core disgust robustly engaged ten clusters of significant activation, primarily encompassing the bilateral amygdala (spreading into the right globus pallidus) and insula, left inferior frontal gyrus, left declive, left fusiform gyrus, left inferior parietal lobule, left middle occipital gyrus, left inferior temporal gyrus, left superior frontal gyrus, bilateral inferior occipital gyrus spreading into the adjacent lingual gyrus (**Table 2, Fig. 2b**). See **Supplementary Materials** for a detailed presentation of percentage of overlap with atlas-based brain regions.

**Table 2.** Brain regions activated during core disgust processing in healthy subjects.

| Volume<br>(mm<sup>3</sup>) | Side | Brain region | BA | MNI coordinates (x, y, z) |  |  | ALE<br>value<br>(10<sup>-2</sup>) | Z |
| --- | --- | --- | --- | --- | --- | --- | --- | --- |
| 2216 | left | Amygdala |  | -22 | -2 | -22 | 3.80 | 6.49 |
| 1736 | left | Declive |  | -34 | -70 | -14 | 2.53 | 4.89 |
|  | left | Fusiform Gyrus | 37 | -44 | -60 | -8 | 2.09 | 4.27 |
|  | left | Fusiform Gyrus | 37 | -42 | -54 | -18 | 1.69 | 3.67 |
| 1696 | right | Amygdala |  | 22 | -6 | -16 | 2.85 | 5.31 |
|  | right | Amygdala |  | 26 | -2 | -24 | 2.33 | 4.62 |
| 1512 | left | Inferior Parietal<br>Lobule | 40 | -62 | -24 | 34 | 3.99 | 6.71 |
| 1472 | left | Middle Occipital<br>Gyrus | 19 | -44 | -76 | 8 | 2.47 | 4.81 |
|  | left | Inferior<br>Temporal Gyrus | 19 | -50 | -76 | 0 | 2.01 | 4.16 |
| 1240 | left | Superior Frontal<br>Gyrus | 9 | -6 | 64 | 26 | 2.21 | 4.44 |
|  | left | Superior Frontal<br>Gyrus | 9 | 2 | 62 | 18 | 2.14 | 4.34 |
| 1120 | right | Inferior Occipital<br>Gyrus | 17 | 14 | -92 | -2 | 2.64 | 5.04 |
| 976 | left | Inferior Frontal<br>Gyrus | 47 | -30 | 32 | -14 | 2.68 | 5.09 |
|  | left | Insula | 13 | -34 | 24 | -2 | 1.73 | 3.72 |
| 896 | right | Insula |  | 40 | 26 | -6 | 2.16 | 4.37 |
| 808 | left | Inferior Occipital<br>Gyrus | 17 | -10 | -98 | -8 | 2.14 | 4.34 |

Abbreviations: ALE = Action Likelihood Estimation, BA = Brodmann Area, MNI = Montreal Neurological Institute.

### 3.3. Social disgust processing

Examining social disgust processing studies revealed six significant clusters of robust activation located in the right insula and adjacent inferior frontal gyrus, bilateral fusiform gyrus spreading into the right declive, left superior frontal gyrus, right cingulate gyrus, right middle occipital gyrus, and right superior temporal gyrus (**Table 3, Fig. 2c**). See **Supplementary Materials** for a detailed presentation of percentage of overlap with atlas-based brain regions.

**Table 3.**
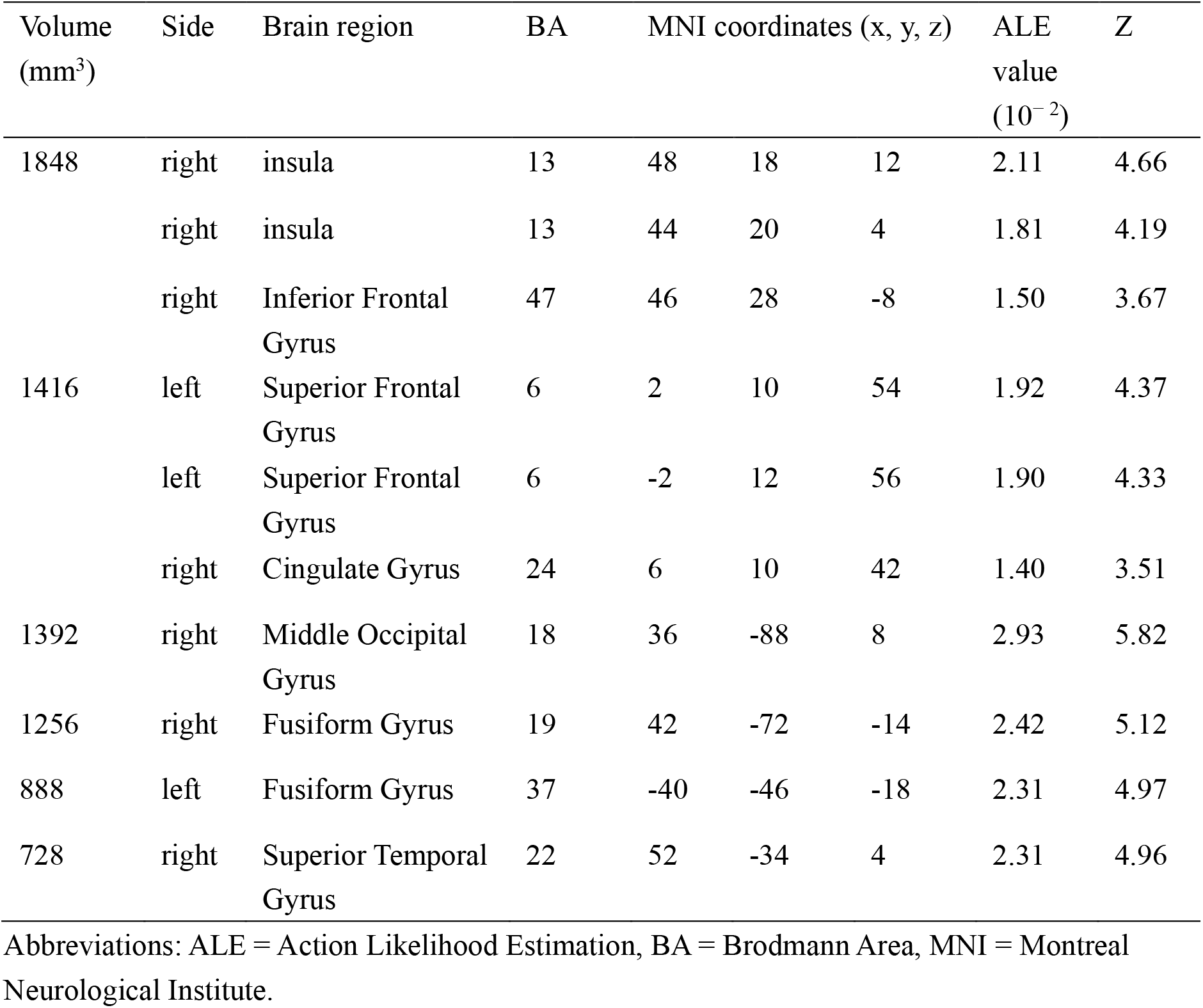
Brain regions activated during social disgust processing in healthy subjects.

| Volume<br>(mm<sup>3</sup>) | Side | Brain region | BA | MNI coordinates (x, y, z) |  |  | ALE<br>value<br>(10<sup>-2</sup>) | Z |
| --- | --- | --- | --- | --- | --- | --- | --- | --- |
| 1848 | right | insula | 13 | 48 | 18 | 12 | 2.11 | 4.66 |
|  | right | insula | 13 | 44 | 20 | 4 | 1.81 | 4.19 |
|  | right | Inferior Frontal<br>Gyrus | 47 | 46 | 28 | -8 | 1.50 | 3.67 |
| 1416 | left | Superior Frontal<br>Gyrus | 6 | 2 | 10 | 54 | 1.92 | 4.37 |
|  | left | Superior Frontal<br>Gyrus | 6 | -2 | 12 | 56 | 1.90 | 4.33 |
|  | right | Cingulate Gyrus | 24 | 6 | 10 | 42 | 1.40 | 3.51 |
| 1392 | right | Middle Occipital<br>Gyrus | 18 | 36 | -88 | 8 | 2.93 | 5.82 |
| 1256 | right | Fusiform Gyrus | 19 | 42 | -72 | -14 | 2.42 | 5.12 |
| 888 | left | Fusiform Gyrus | 37 | -40 | -46 | -18 | 2.31 | 4.97 |
| 728 | right | Superior Temporal<br>Gyrus | 22 | 52 | -34 | 4 | 2.31 | 4.96 |

Abbreviations: ALE = Action Likelihood Estimation, BA = Brodmann Area, MNI = Montreal Neurological Institute.

### 3.4. Common and differential brain regions for core and social disgust processing

Examining the common brain systems by means of conjunction analyses revealed that the right inferior frontal gyrus (extending into the anterior insula) and left fusiform gyrus were robustly activated during core and social disgust processing (domain-general disgust processing) (**Table 4, Fig. 3a**).

**Table 4.**
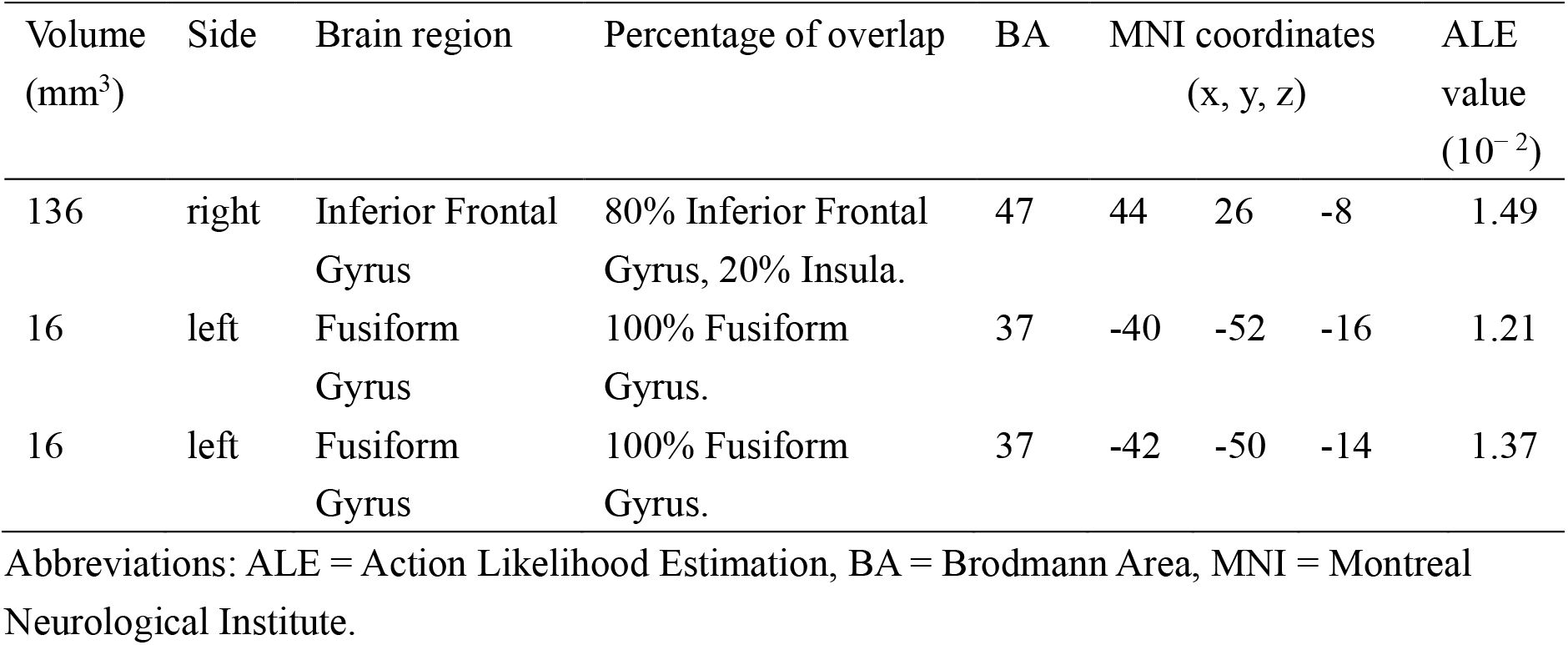
Brain regions activated by conjunction analysis between core and social disgust processing in healthy subjects.

**Fig. 3.**
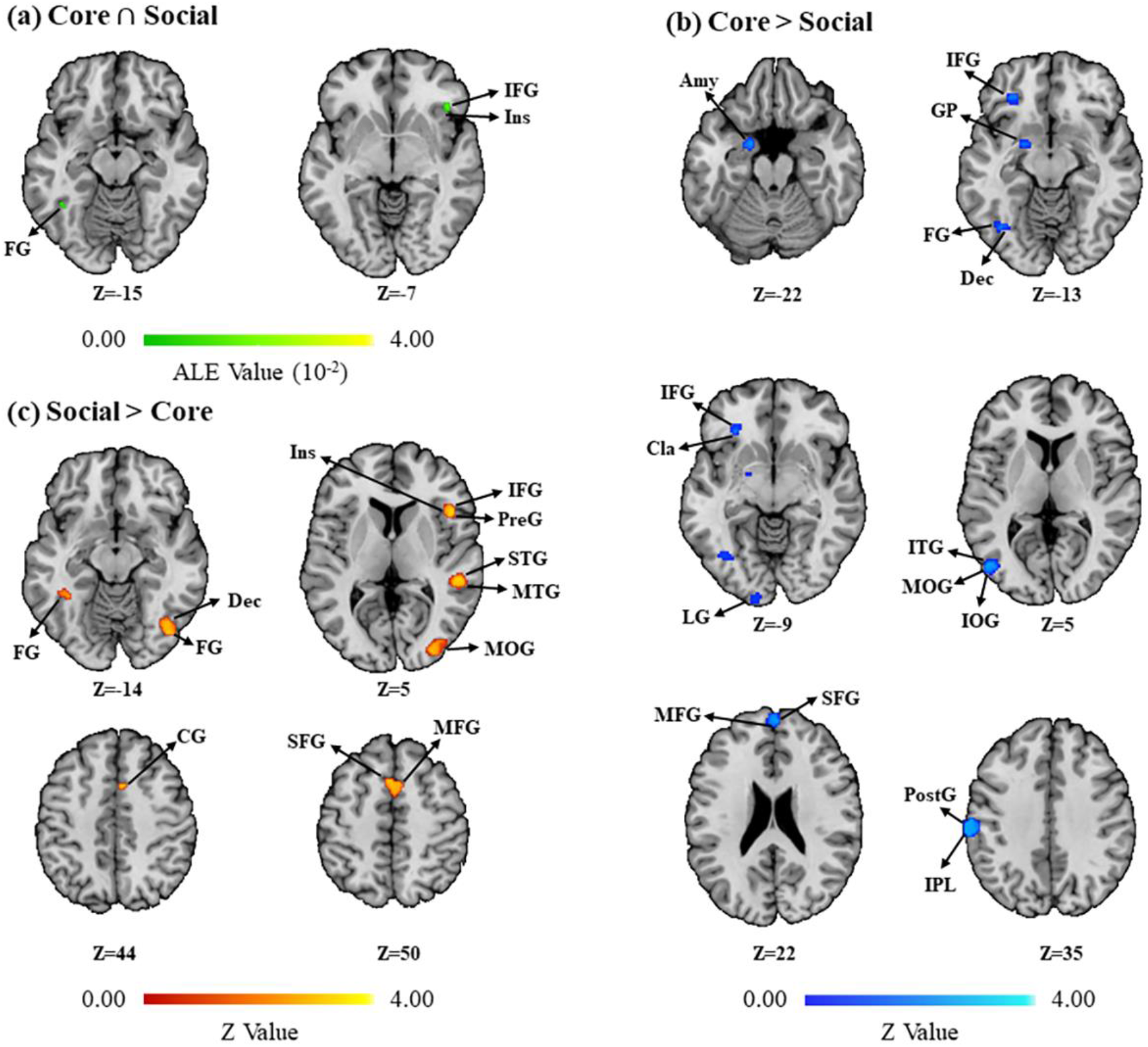
Overview of significant clusters from the conjunction and contrast analyses. **(a)** Common activations for core and social disgust processing in healthy subjects (domain-general disgust processing). **(b)** Greater activations for Core disgust processing (Core > Social) in healthy subjects (domain-specific regions for core disgust). **(c)** Greater activations for social disgust processing (Social > Core) in healthy subjects (domain-specific regions for social disgust). Coordinates are in the MNI space. Abbreviations: Amy = Amygdala, CG = Cingulate Gyrus, Cla = Claustrum, Dec = Declive, FG = Fusiform Gyrus, GP = Globus Pallidus, IFG = Inferior Frontal Gyrus, Ins = Insula, IOG = Inferior Occipital Gyrus, IPL = Inferior Parietal Lobule, ITG = Inferior Temporal Gyrus, LG = Lingual Gyrus, MFG = Medial Frontal Gyrus, MOG = Middle Occipital Gyrus, MTG = Middle Temporal Gyrus, PostG = Postcentral Gyrus, PreG = Precentral Cyrus, SFG = Superior Frontal Gyrus, STG = Superior Temporal Gyrus.

The contrast analyses revealed stronger recruitment of the left inferior parietal lobule, left postcentral gyrus, left middle occipital gyrus, left inferior temporal gyrus, left limbic-striatal cluster (encompassing the amygdala, hippocampus and globus pallidus), right superior frontal gyrus, left medial frontal gyrus, left inferior frontal gyrus, left declive, left fusiform gyrus and left lingual gyrus for core disgust compared with social disgust (domain-specific core disgust processing), suggesting a left hemisphere predominance (**Table 5, Fig. 3b**). In contrast social disgust processing was characterized by a stronger engagement of the right inferior frontal gyrus (extending to the insula), left medial frontal gyrus, right cingulate gyrus, right middle occipital gyrus, right declive, right superior temporal gyrus, right middle temporal gyrus, and bilateral fusiform gyrus as compared to core disgust processing (domain-specific social disgust processing), reflecting a right hemispheric predominance (**Table 5, Fig. 3c**). We additionally repeated the contrast and conjunction analyses with the same number of permutations as employed in the single dataset analyses (5,000 permutations, p < 0.05, uncorrected, minimum cluster size of 100 mm^3^) which generated a highly similar pattern of results as the initial analyses (details are reported in **Supplementary Table 2** and **Supplementary Table 3**).

**Table 5.**
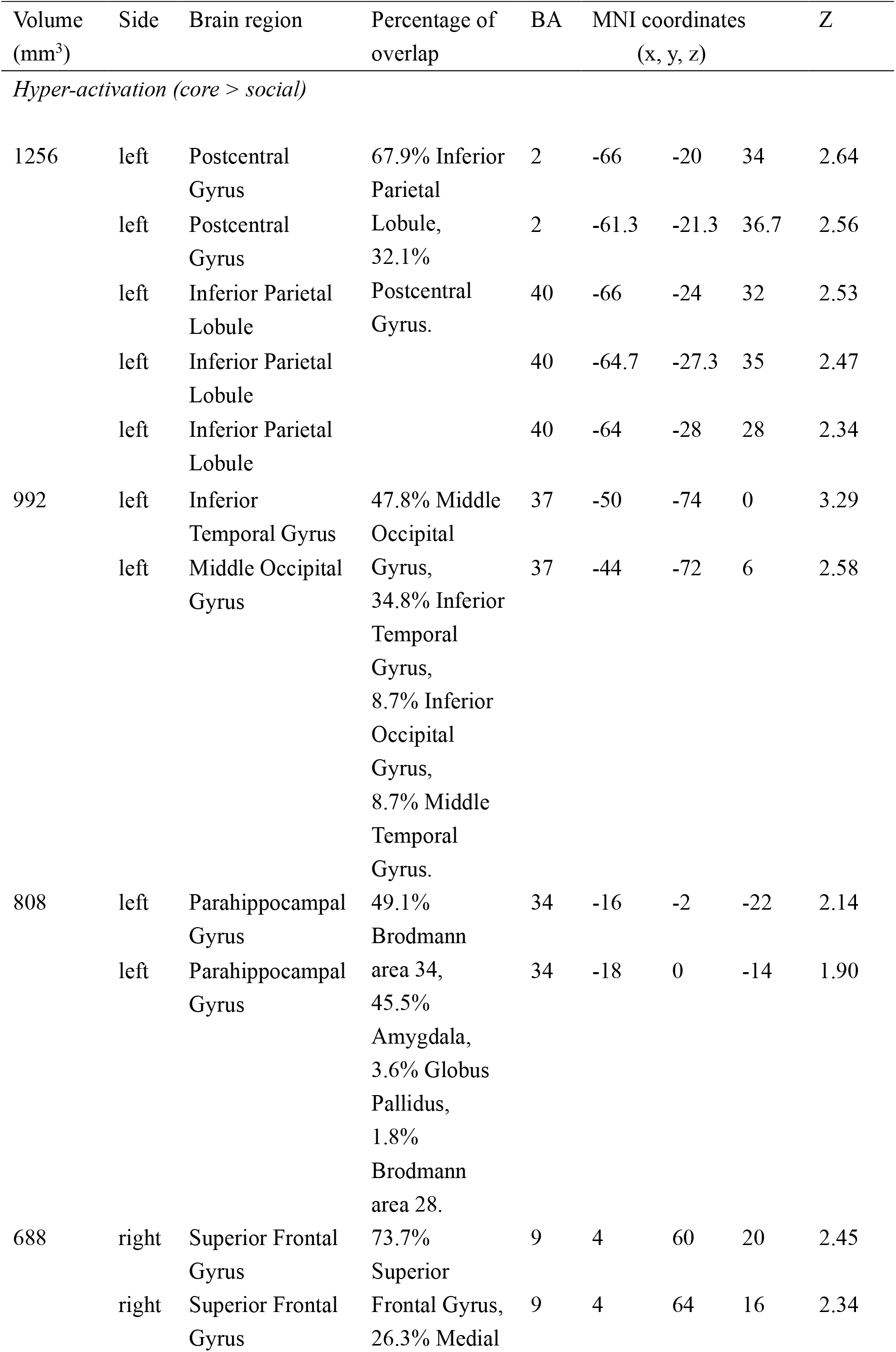
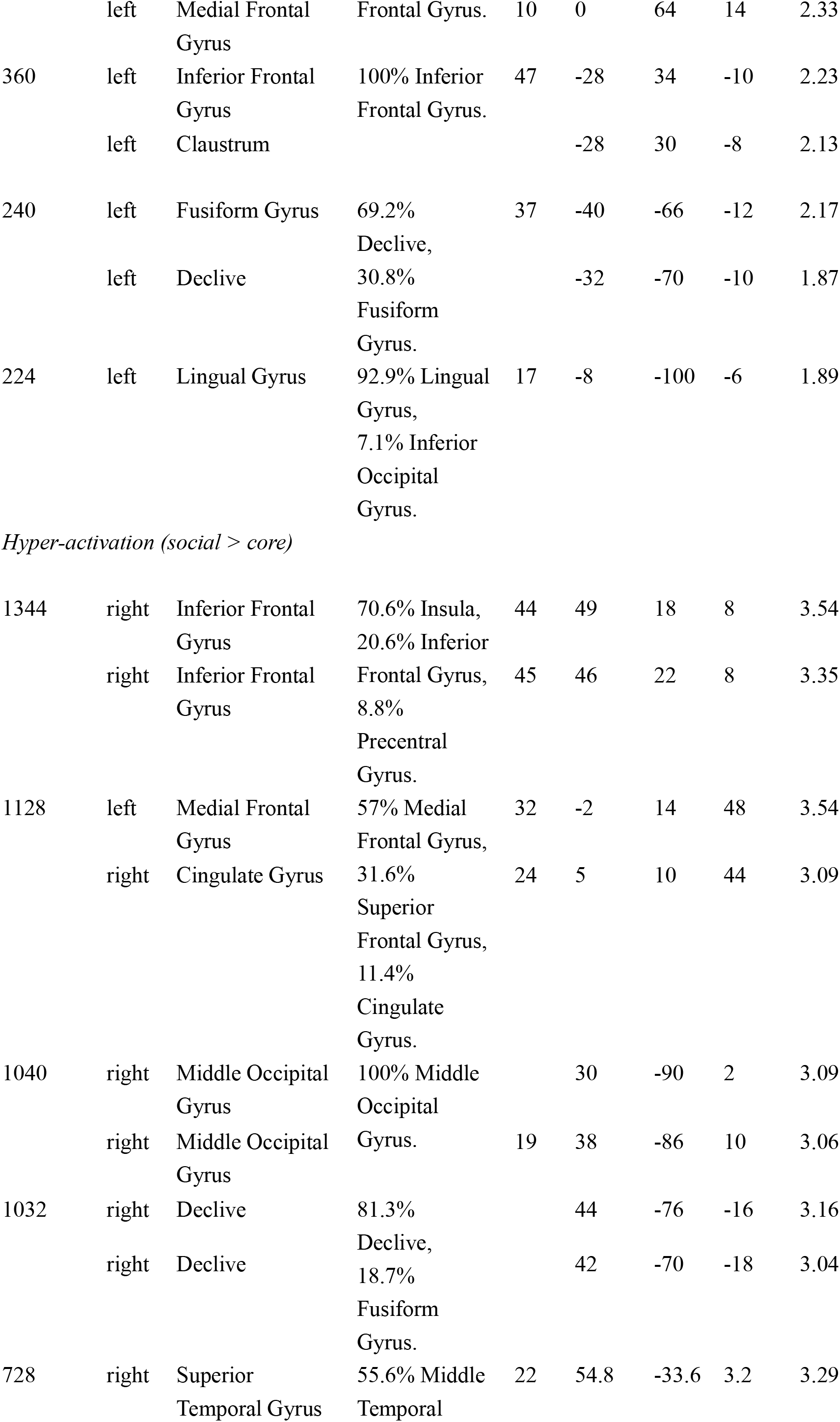
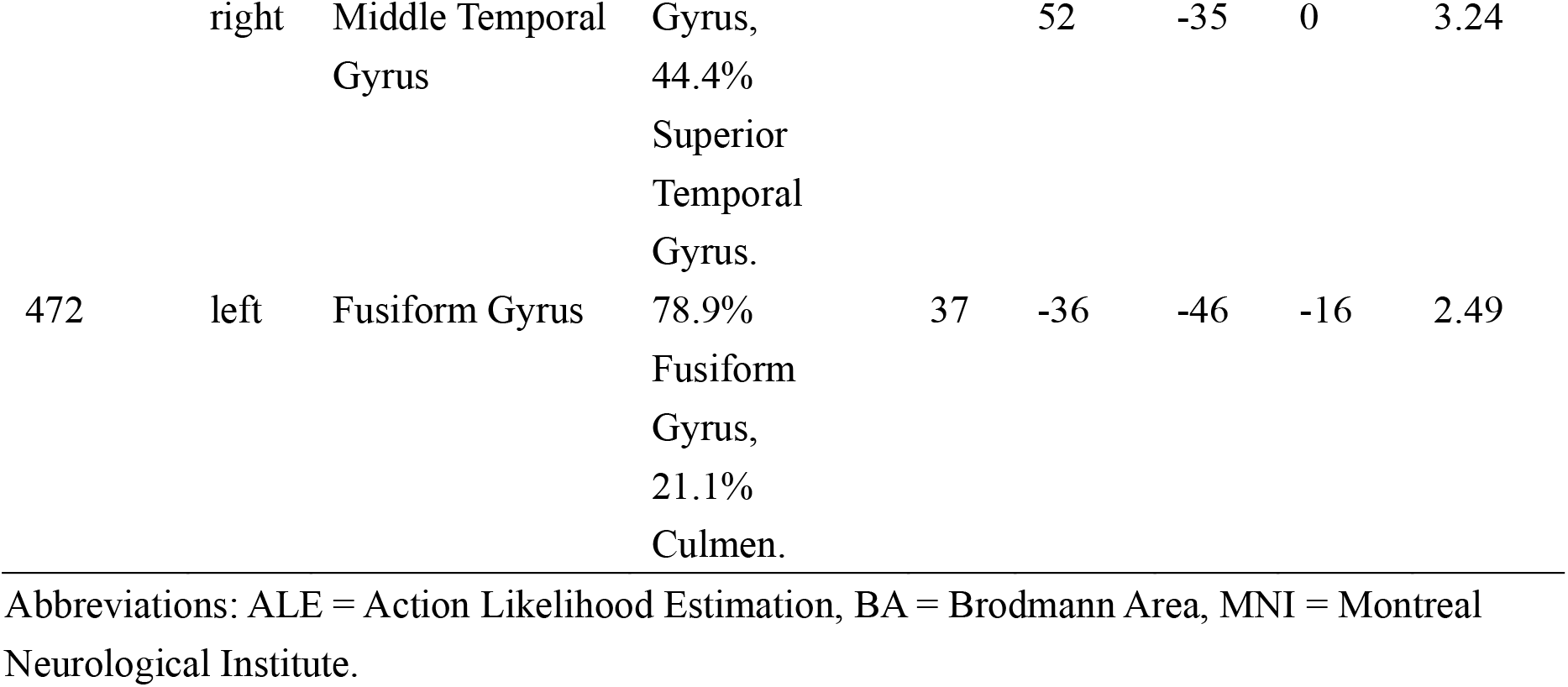
Brain regions activated by contrast analysis between core and social disgust processing in healthy subjects.

### 3.5. Network level functional characterization via MACM analyses

We adopted two different approaches (labeled as MACM A as well as MACM B) aiming to unveil the functional network of right inferior frontal gyrus and left fusiform gyrus from conjunction analyses across core and social disgust processing.

From MACM-A, co-activation maps for the right inferior frontal gyrus included bilateral insula, bilateral inferior frontal gyrus, right anterior cingulate, bilateral medial frontal gyrus and left cingulate gyrus (**Table 6, Fig. 4a**). Co-activation maps for the left fusiform gyrus were significant for bilateral fusiform gyrus, bilateral parahippocampal gyrus, bilateral amygdala, right putamen and right inferior occipital gyrus (**Table 7, Fig. 4a**).

**Table 6.** Network of right inferior frontal gyrus revealed by MACM-A analysis.

| Volume<br>(mm<sup>3</sup>) | Side | Brain region | BA | MNI coordinates (x, y, z) |  |  | ALE value<br>(10<sup>-2</sup>) | Z |
| --- | --- | --- | --- | --- | --- | --- | --- | --- |
| 5352 | right | Inferior Frontal Gyrus | 47 | 40 | 26 | -8 | 3.54 | 5.85 |
|  | right | Insula | 13 | 46 | 10 | -8 | 2.47 | 4.46 |
|  | right | Inferior Frontal Gyrus | 46 | 38 | 36 | 6 | 2.42 | 4.39 |
|  | right | Inferior Frontal Gyrus | 13 | 38 | 28 | 2 | 2.14 | 3.99 |
| 4808 | left | Insula | 13 | -34 | 24 | -2 | 3.88 | 6.25 |
|  | left | Inferior Frontal Gyrus | 46 | -48 | 38 | 4 | 1.98 | 3.75 |
| 1536 | right | Anterior Cingulate | 32 | 6 | 50 | -4 | 2.43 | 4.41 |
|  | right | Medial Frontal Gyrus | 11 | 2 | 38 | -22 | 2.13 | 3.97 |
|  | left | Medial Frontal Gyrus | 11 | 0 | 42 | -26 | 2.08 | 3.90 |
|  | right | Anterior Cingulate | 32 | 4 | 46 | -16 | 1.94 | 3.69 |
| 1192 | left | Cingulate Gyrus | 32 | -8 | 26 | 42 | 3.24 | 5.48 |
| 1104 | left | Inferior Frontal Gyrus | 9 | -46 | 12 | 26 | 2.27 | 4.18 |

Abbreviations: ALE = Action Likelihood Estimation, BA = Brodmann Area, MNI = Montreal Neurological Institute.

**Table 7.** Network of left fusiform gyrus revealed by MACM-A analysis.

| Volume<br>(mm<sup>3</sup>) | Side | Brain region | BA | MNI coordinates (x, y, z) |  |  | ALE<br>value<br>(10<sup>-2</sup>) | Z |
| --- | --- | --- | --- | --- | --- | --- | --- | --- |
| 2776 | left | Fusiform Gyrus | 37 | -44 | -50 | -14 | 2.04 | 5.18 |
|  | left | Parahippocampal<br>Gyrus | 19 | -40 | -48 | -6 | 1.58 | 4.39 |
| 2240 | right | Amygdala |  | 30 | -4 | -20 | 1.88 | 4.90 |
|  | right | Parahippocampal<br>Gyrus | 28 | 20 | -16 | -24 | 1.18 | 3.59 |
|  | right | Putamen |  | 34 | 4 | -10 | 1.16 | 3.55 |
| 1872 | left | Amygdala |  | -26 | -2 | -22 | 2.27 | 5.56 |
| 1184 | right | Inferior Occipital<br>Gyrus | 19 | 48 | -74 | -2 | 2.21 | 5.46 |
|  | right | Fusiform Gyrus | 19 | 40 | -72 | -14 | 1.43 | 4.11 |

Abbreviations: ALE = Action Likelihood Estimation, BA = Brodmann Area, MNI = Montreal Neurological Institute.

**Fig. 4.**
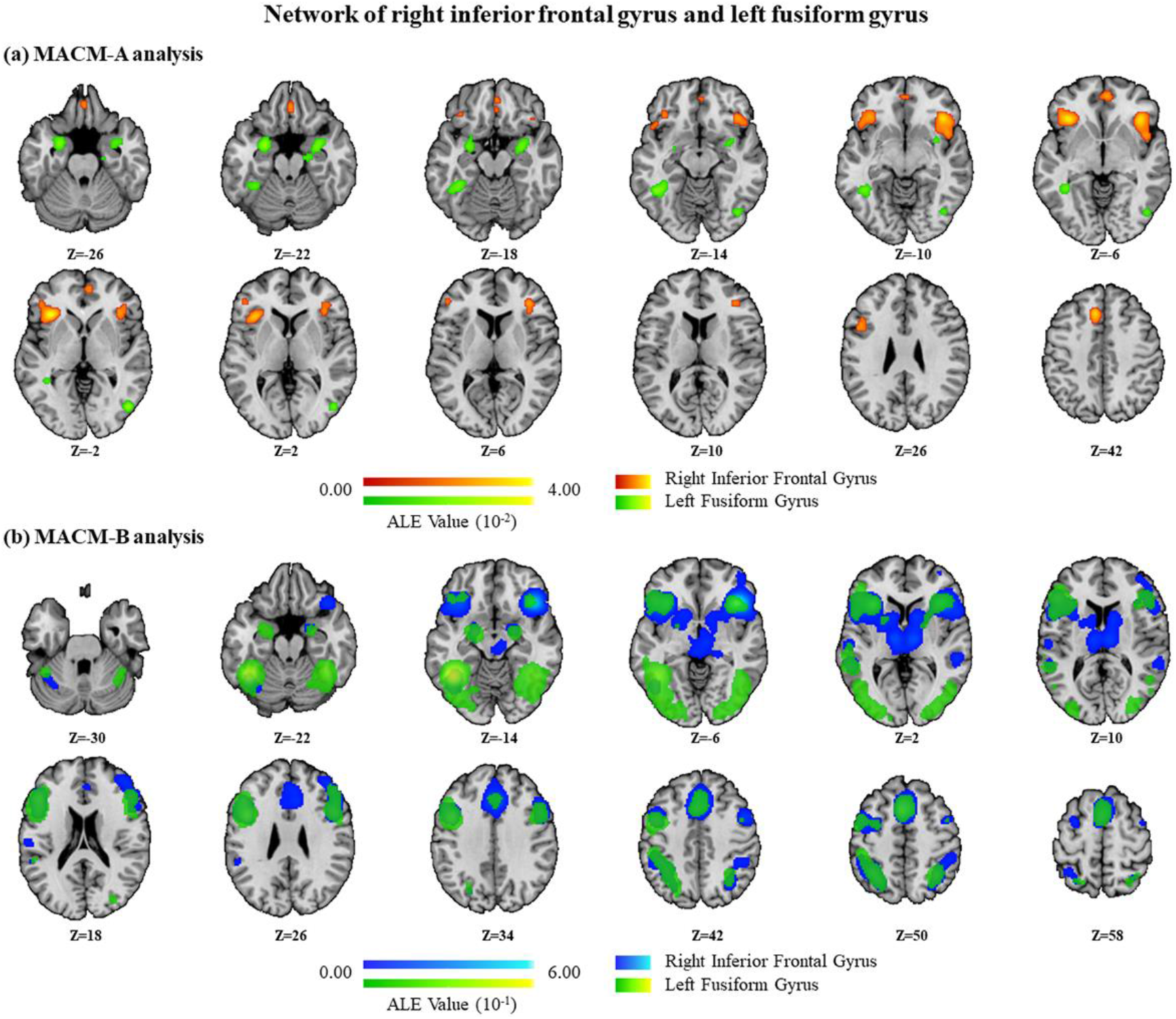
Overview of the functional networks of right inferior frontal gyrus and left fusiform gyrus by MACM analyses. **(a)** Network of right inferior frontal gyrus and left fusiform gyrus by MACM-A analyses. **(b)** Network of right inferior frontal gyrus and left fusiform gyrus by MACM-B analyses. Coordinates are in the MNI space.

The additional functional network analyses (MACM-B) of right inferior frontal gyrus showed converging results as with MACM-A analyses, including bilateral inferior frontal gyrus, bilateral insula, bilateral medial frontal gyrus, left cingulate gyrus, and right anterior cingulate. MACM-B analyses of left fusiform gyrus also showed highly overlapping regions with MACM-A analyses, including bilateral fusiform gyrus, bilateral amygdala, right putamen and right inferior occipital gyrus (**Fig. 4b**, see **Supplementary Table 4** and **Supplementary Table 5** for detailed activated regions). In other words, both functional network characterization analyses confirmed each other.

### 3.6. Functional characterization of the ALE maps on the behavioral level

We conducted three separate quantitative functional annotation analyses by testing the associations of functional terms provided by the Neurosynth database with the brain regions robustly engaged during domain-general disgust processing generated from conjunction analyses as well as core disgust processing and social disgust processing obtained from single dataset analyses. The top 20 terms were visualized with a larger font size reflecting a smaller p-value (see **Fig. 5**). Whereas the domain-general network was mainly characterized by general attention and emotional processes (**Fig. 5a**), core disgust processing regions primarily associated with threat- and avoidance-related processes (**Fig. 5b**), while social disgust processing revealed social- and adaptation-related terms (**Fig. 5c**).

**Fig. 5.**
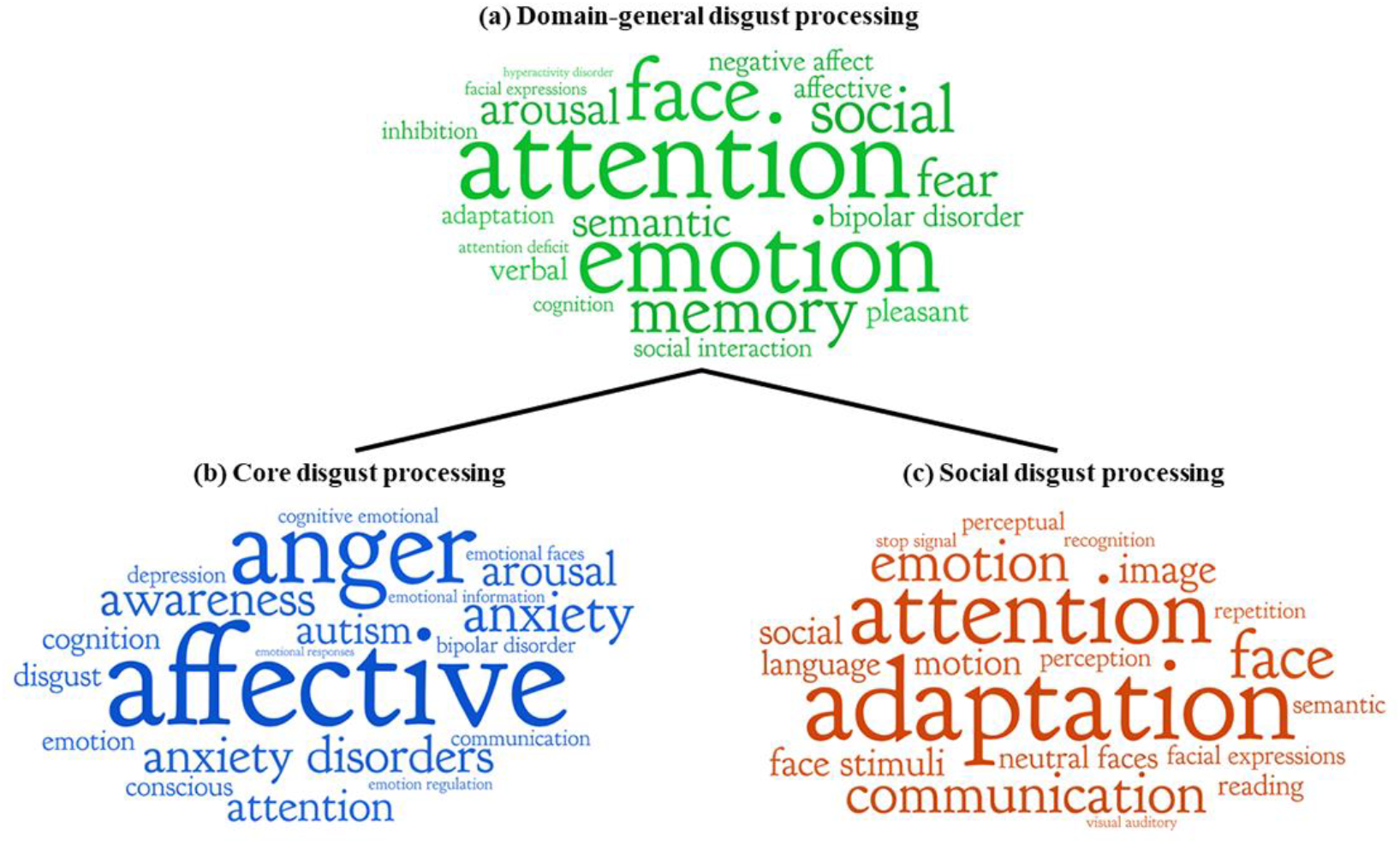
Functional characterization of the neural correlates from three categories. **(a)** Neural substrates for domain-general disgust processing (Core ∩ Social). **(b)** Neural substrates for core disgust processing. **(c)** Neural substrates for social disgust processing. Only the top 20 terms are displayed. The font size of words is proportional to the permutation p value for association between user-provided cluster of regions and each functional term provided by Neurosynth, with smaller p value indicating bigger term size.

## 4. Discussion

Previous studies on the brain systems that mediate disgust-related processes and initial studies that examined common and separable neural representations of direct exposure to disgust elicitors (core disgust) and social-communicative signals of disgust (social disgust) revealed inconsistent results. Against this background, we employed neuroimaging meta-analyses that capitalized on a large number of previous fMRI studies to determine the neural substrates robustly engaged in general, core as well as social disgust processing, respectively. Next, by employing meta-analytic conjunction and contrast analyses, we further segregated common (domain-general) and distinct (domain-specific) neural correlates of core and social disgust processing and utilized meta-analytic behavioral and network level decoding to functionally characterize the identified brain regions.

### 4.1. Brain systems mediating general, core and social disgust processing

The overarching meta-analysis pooling the data from the 69 qualified studies, comprising 78 eligible contrasts (general disgust processing) yielded robust engagement of a bilateral network encompassing primarily the bilateral anterior insular cortex, bilateral amygdala and adjacent globus pallidus, inferior and superior frontal, visual occipital (fusiform gyrus, inferior and middle occipital gyri, lingual gyrus) and cerebellar regions. Separate analyses of the subdomains of core and social disgust processing revealed that the core disgust network largely overlapped with the general disgust network while the social disgust processing network additionally engaged superior temporal regions involved in social processing and face processing regions such as fusiform-extrastriate and frontal cortices, mainly in the right hemisphere.

The identified network for general disgust processing largely resembles regions that have been consistently engaged in disgust-related processes (Fusar-Poli et al., 2009b; Kirby and Robinson, 2017; Vytal and Hamann, 2010; Wager et al., 2015) and may reflect the multifaceted nature of the disgust response. The identified regions may support the engagement of different simultaneous processes which allow a concerted disgust reaction, ranging from initial detection of biologically salient stimuli in occipital visual regions (Tao et al., 2021), early threat detection and implementation of the defensive motor response by the amygdala, globus pallidus, fusiform and cerebellar regions (e.g., Phelps and LeDoux, 2005; Surguladze et al., 2003; Yao et al., 2018b), as well as salience allocation and interoceptive awareness of the emotional state in the insula cortex (Craig, 2009; Li et al., 2019; Uddin, 2015), and the subjective experience and interpretation of the emotional state in prefrontal regions (LeDoux and Pine, 2016).

Examining the core disgust network, i.e., regions robustly activated by disgust eliciting visual stimuli, revealed a similar network primarily encompassing the bilateral insula and adjacent inferior frontal regions, the bilateral amygdala and adjacent globus pallidus as well as temporo-occipital regions. The overlap may reflect that the direct exposure to potential infectious and harmful stimuli triggers the evolutionary defensive-avoidance circuits engaged in early salience and threat detection and the initiation of the physiological and behavioral defensive response, thus facilitating rapid withdrawal. Previous studies have for instance extensively documented reactivity of the amygdala towards aversive stimuli (Zald, 2003) with meta-analytic data demonstrating a higher probability of amygdala activation for stimuli signaling a potential danger, including fear and disgust inducing stimuli, as compared to non-threatening stimuli (Costafreda et al., 2008), indicating a critical role in threat detection and initiation of protective responses (Mihov et al., 2013; Schienle et al., 2016). The amygdala receives dense projections from the visual cortex and via projections with cerebellar regions and the globus pallidus initiates motor and physiological defensive responses towards aversive stimuli (Giovanniello et al., 2020; Lindquist, 2020; Salih et al., 2009; Sambataro et al., 2006), while bilateral projections to the orbitofrontal cortex further mediate withdrawal or avoidance behavior (Rolls, 1999, 2000b). In support of this interpretation, the meta-analytic functional characterization of the identified core disgust network revealed primarily terms associated with negative affective responses, including e.g., anxiety, arousal as well as disgust.

Separately examining the social disgust studies revealed a right lateralized network encompassing the right anterior insula and inferior frontal gyrus, middle occipital cortex, fusiform gyrus, superior temporal gyrus and cingulate regions. The corresponding meta-analytic functional decoding of the regions revealed primarily terms associated with salience and social communication, including attention as well as face and communication. In line with the functional decoding, the identified regions have been previously associated with emotional face processing (Fusar-Poli et al., 2009b), with the middle occipital cortex involved in early face processing stages (Tu et al., 2013) and the fusiform gyrus being involved in visual attention and higher level face perception processing (Weiner and Zilles, 2016), while superior temporal regions have been strongly involved in social cognitive processes including theory of mind and mentalizing (Frith, 2007; Zilbovicius et al., 2006).

In contrast to several previous original studies, the present meta-analysis did not reveal a robust engagement of the amygdala or main basal ganglia regions (i.e., caudate and putamen) during social or general disgust processing, respectively. However, despite a number of previous studies reporting amygdala engagement during the perception of disgusted faces, a previous meta-analysis failed to corroborate amygdala engagement during disgust faces processing (Fusar-Poli et al., 2009b) and previous studies found either strongest amygdala responses to fear (Mattavelli et al., 2014) or comparable responses to emotional and neutral faces (Derntl et al., 2009; Fitzgerald et al., 2006; Winston et al., 2003) while amygdala lesion patients exhibited intact disgusted face decoding (Becker et al., 2012). With respect to the role of the basal ganglia in disgust, previous imaging (Ahn et al., 2014; Benuzzi et al., 2008; Lassalle et al., 2019; Phillips et al., 2000; Phillips et al., 2004; Phillips et al., 1998; Phillips et al., 1997; Pujol et al., 2018; Shapira et al., 2003; Sprengelmeyer et al., 1998; Tettamanti et al., 2012; Ziegler et al., 2018) and lesion studies (Calder et al., 2000; Sprengelmeyer et al., 1996; Straube et al., 2010) revealed inconsistent results and the present meta-analysis did not confirm a robust engagement.

### 4.2. Common and distinct neural correlates of core and social disgust processing

Notably the right anterior insula was consistently and specifically identified across both, the separate analyses for core disgust and social disgust. Although the consistent involvement of the insula in disgust processing has long been debated in the original disgust literature (Deeley et al., 2008; Gorno-Tempini et al., 2001; Schienle et al., 2006; Schienle et al., 2005b; Stark et al., 2005; Stark et al., 2004; Stark et al., 2003; Trautmann et al., 2009), several previous studies reported engagement of this region during core disgust as well as social disgust processing (Chapman and Anderson, 2012; Fusar-Poli et al., 2009b; Vytal and Hamann, 2010). Further support for a role of the insula in disgust processing comes from lesion (Adolphs et al., 2003; Calder et al., 2000; Kipps et al., 2007) and intracerebral recording studies (Krolak-Salmon et al., 2003), and studies in healthy subjects reporting a convergent engagement of the insula across disgust paradigms employing not only visual but also olfactory (Heining et al., 2004; Wicker et al., 2003), gustatory (Corradi-Dell’Acqua et al., 2016; Jabbi et al., 2008) and imagination (Jabbi et al., 2008) procedures. The insula is a functionally heterogenous region and has been involved not only in general salience and interoceptive processes but also in the awareness and experience of emotional states (e.g., Craig, 2009; Uddin, 2015; Zhou et al., 2020). Previous studies have demonstrated that negative affective experience induced by visual stimuli (Viinikainen et al., 2010), subjective experienced unpleasantness (Benuzzi et al., 2008), as well as aversive electrical stimulations and unfair offers in social interactions (Corradi-Dell’Acqua et al., 2016) consistently engage the anterior insula, confirming its potential role in the representation of negative emotional experiences.

Using a conjunction and contrast meta-analytic approach, we aimed to further segregate common and domain-specific neural correlates for core and social disgust processing. In line with the observations from the separate analyses, the right anterior insula extending into the adjacent inferior frontal gyrus and the left fusiform gyrus specifically exhibited convergent engagement during core and social disgust processing. Together with the anterior insula, the inferior frontal cortex plays an important role in general emotional processes such as evaluation of biological significant stimuli (Rolls, 2000a, 2004), interoception (Critchley et al., 2004), multimodal representation emotion salient stimuli (Yamasaki et al., 2002), and emotion recognition (Sprengelmeyer et al., 1998) while the fusiform gyrus plays a role in higher order visual processing (Fusar-Poli et al., 2009b) and emotional reactivity towards negative and threatening stimuli (Kirby and Robinson, 2017; Tao et al., 2021). Initial original studies have begun to examine the shared neural substrates for core and social disgust. For example, Wicker et al. (2003) found that observing disgusted faces activated the anterior insula which was also implicated in first-person disgust experience evoked by unpleasant odorants (Wicker et al., 2003). Similarly, Jabbi et al. (2008) observed that the anterior insula was activated both during the observation of another person’s disgusted facial expressions and during the taste of disgusting liquids (Jabbi et al., 2008). A subsequent model from Vicario et al. that was based on data from clinical populations proposed that the insula mediates disgust processing in both core and social disgust domains (Vicario et al., 2017) and explained the common activity between core and social disgust in the context of embodied simulation theories of emotion perception (Gallese and Goldman, 1998; Gallese and Sinigaglia, 2011; Vicario et al., 2017; Wicker et al., 2003). These theories propose that observing a facial emotional expression (i.e., a disgusted face, in the present work referred to as *social disgust*) triggers activation in some of the neural systems engaged in the actual (first-person) experience of the same emotion (core disgust). The present findings extend the previous literature and suggest that the right anterior insula and left fusiform gyrus may support this emotion perception mechanism for disgust. A number of previous studies reported an involvement of the insula in the first-hand experience of pain and vicarious pain induced by observing another person in a painful situation or expressing a painful facial expression (Bzdok et al., 2012; Corradi-Dell’Acqua et al., 2016; Fan et al., 2011; Singer et al., 2004). In the context of the present findings this may indicate a rather general role of the insula in mirroring during different aversive emotional states such as pain and disgust. In contrast the fusiform gyrus has not been consistently reported in emotional empathy or mirroring studies and thus may represent a rather process-specific common neural substrate in the domain of observed and self-experienced disgust. Given the role of the fusiform gyrus in visual processing and its dense projections with the amygdala (Ardila et al., 2015; Fusar-Poli et al., 2009b; Weiner and Zilles, 2016), this region may support an early discrimination of salient visual stimuli across disgust-related processes (for a potential role of the fusiform gyrus in not face-related fear and threat detection see e.g., Larson et al., 2009; Mueller and Pizzagalli, 2016).

Further meta-analytic network decoding via MACM demonstrated that the identified anterior insula cortex region coupled with a bilateral network encompassing the insula and ventrolateral prefrontal regions as well as anterior cingulate and medial frontal regions. The lateral prefrontal cortex, insula and anterior cingulate exhibit strong anatomical and functional connections (Craig, 2009; Mesulam and Mufson, 1982b) and represent key hubs within the salience network, the fronto-parietal executive control network and the sensorimotor networks (Belyk et al., 2017; Cloutman et al., 2012; Cox et al., 2010). Functional interactions between these regions have been involved in attentional processes (Wang et al., 2019), emotion regulation (Morawetz et al., 2017) and goal-directed behavior (Dosenbach et al., 2006), which may support disgust processing across core and social domains. The identified left fusiform gyrus network comprised a limbic-occipital network encompassing the amygdala, parahippocampal gyrus, putamen and inferior occipital regions. The functional interplay between these regions has been demonstrated in several previous studies (Amting et al., 2010; Fairhall and Ishai, 2007; Herrington et al., 2011; Molapour et al., 2015; Turk-Browne et al., 2010) and plays an important role in early threat detection (Tao et al., 2021), suggesting that this network may facilitate detection of potential threats across core and social disgust processes.

Examining regions specifically engaged during the two disgust processing domains revealed stronger activity of several regions in the left hemispheric disgust network during core disgust processing, including e.g., occipito-temporal visual, limbic, inferior frontal and parietal regions. These regions mainly mirror the results of the core disgust processing network and may suggest a neurobiological model for domain-specific core disgust processing in which visual disgust stimuli undergo an early screening in occipito-temporal, which – due to the salience of the stimuli – encompass both, primary and secondary visual cortex processing (Bradley et al., 2003). Next the information is transmitted via inferior temporal pathways to the amygdala and orbitofrontal cortex regions strongly engaged in emotional evaluation (Rolls, 2000b), while the parahippocampal gyrus may further qualify the contextual relevance of the stimuli (Tao et al., 2021). The survival relevant reactivity moreover requires rapid withdrawal behavior (Rolls, 1999, 2000b), which are potentially mediated by prefrontal, inferior parietal and pallidal regions. In particular prefrontal regions (e.g., superior and medial frontal gyrus) exhibit dense connections with parietal, temporal, and occipital regions (Cechetto and Topolovec, 2002) and play a key role in both, emotional awareness and executive control, including attention and initiation of appropriate responses (LeDoux, 2014; LeDoux and Pine, 2016; Rozzi and Fogassi, 2017) while the inferior parietal lobule is an important locus for motor planning (Buxbaum and Randerath, 2018) and receives input from both, visual as well as somatosensory regions thus being optimal positioned to integrate these modalities (Andersen, 2011). The globus pallidus finally has been implicated in implementing rapid motor reactions in response to emotional stimuli (Sambataro et al., 2006) and via connections with the amygdala regulates responses to (unconditioned) threat stimuli (Giovanniello et al., 2020). Together these findings indicate that – compared to social disgust stimuli - exposure to disgusting and thus potentially dangerous stimuli engages the core defensive-avoidance circuits.

In contrast, for domain-specific social disgust processing, a right lateralized network encompassing anterior insula (extending into the inferior frontal gyrus), middle occipital gyrus (occipital face area), superior temporal gyrus and anterior cingulate regions as well as the left fusiform face area exhibited stronger activation. In the context of previous face processing models (Haxby et al., 2000; Liu et al., 2021), the present findings may suggest a model of social disgust processing that aligns with the ventral and dorsal face processing streams. The dorsal pathway includes superior temporal and middle temporal regions, while the ventral pathway comprises of occipital face area, fusiform face area, and inferior frontal regions. The dorsal pathway has been implicated in emotional face processing (Britton et al., 2006; Fusar-Poli et al., 2009b) and may facilitate subsequent social cognitive processes including theory of mind and mentalizing (Frith, 2007; Zilbovicius et al., 2006), while the ventral stream is strongly engaged in determining dynamic processes of facial emotion processing, such that the right middle occipital gyrus may relay the information of disgusted faces to the left fusiform face area, and the right inferior frontal gyrus may play a key role in maintaining attention to the social information in the perception of disgusted facial expressions (Liu et al., 2021). In addition to core regions of these pathways the social disgust systems encompassed the right insula, right anterior cingulate gyrus, left prefrontal regions and right declive, which all have been involved in face and particular emotional face processing (Britton et al., 2006; Fusar-Poli et al., 2009a; Fusar-Poli et al., 2009b; Liu et al., 2021; Sabatinelli et al., 2011) and accumulating evidence from human and non-human primate models suggests a role of the insula in decoding the emotional content of faces (Desimone et al., 1984; Haxby et al., 2000; Mesulam and Mufson, 1982a; Perrett et al., 1982) as well as emotional awareness and interoception (Craig, 2009) and disgust processing it is conceivable that activation in this region may support the emotional and cognitive decoding of the facial disgust signals.

Notably, a strong lateralization effect was observed such that core disgust processing primarily engaged a left-lateralized, while social disgust processing primarily engaged a right-lateralized network. Although we did not have a priori lateralization hypothesis, the findings resonate with some previous studies suggesting a left lateralization of withdrawal-related activity (Tucker et al., 1995; Wager et al., 2003) while conscious emotional perception is represented in a right lateralized network (Adolphs, 2002; Adolphs et al., 1996; Davidson and Irwin, 1999; Oliveri et al., 2003). However, the explanation of a left-lateralized withdrawal-network stands in contrast with overarching approach/avoidance models (Chen et al., 2021; Güntürkün et al., 2020; Palomero-Gallagher and Amunts, 2021), which assumes that the left hemisphere is relatively specialized in processing approach behavior, while the right hemisphere is relatively specialized in processing avoidance or withdrawal behavior. However, evidence for either of the proposed models is not unequivocal such that for instance several previous meta-analyses reported bilateral networks in response to withdrawal-related stimuli including threat- and disgust-related stimuli (Fusar-Poli et al., 2009a; Fusar-Poli et al., 2009b; Tao et al., 2021; Vytal and Hamann, 2010).

### 4.3. Specificity of the identified brain systems for disgust-related processes

While several avoidance-related processes such as the processing of threat- and disgust-related stimuli has been associated with increased activity in the defensive networks described above evidence from lesion studies suggests a relative specialization of the underlying systems. For instance, evidence from previous lesion models showed that insula lesion led to deficits in disgust processing (Adolphs et al., 2003; Calder et al., 2000; Cantone et al., 2019; Holtmann et al., 2020; Kipps et al., 2007) while amygdala lesion led to deficits in fear processing (Adolphs et al., 2005; Becker et al., 2012; Calder et al., 1996; Feinstein et al., 2011; Mihov et al., 2013). However, the conventional fMRI approach is limited with respect to disentangling process-specific neurofunctional correlates (Woo et al., 2017; Zhou et al., 2021). Recent studies thus adopted multivariate pattern analyses which allows a more precise determination of common and separable neurofunctional representations revealed for instance that the anterior insula represents vicarious and self-experienced pain, however, the common local patterns in this regions also represented disgust (Corradi-Dell’Acqua et al., 2016), or that both, empathic pain evolved by the observation of noxious stimulation of body limbs and painful facial expressions share a common representation in the mid-insula (Zhou et al., 2020), while other defensive responses such as fear and conditioned threat exhibit separable neurofunctional representations (Zhou et al., 2021). Within this context the specificity of the determined defensive network for disgust versus other defensive responses needs to be further determined in future studies.

### 4.4. Limitations

Results of the present study need to be considered in the context of several limitations. First, core disgust in the present study included contamination and envelope violations (Vicario et al., 2017) while some previous disgust conceptualizations differentiate these stimuli (Haidt et al., 1994; Olatunji and Sawchuk, 2005; Olatunji et al., 2007; Rozin and Fallon, 1987; van Overveld et al., 2009). Initial neuroimaging studies revealed inconsistent support of this notion (Borg et al., 2012; Harrison et al., 2010; Schienle et al., 2006; Wright et al., 2004) and an increasing number of neuroimaging studies on disgust may allow a reasonable powered examination of further disgust subprocesses in the future. Second, in line with the conceptualization in Vicario et al. (2017) *social disgust* in the present study specifically referred to experiments that employed disgusted facial expression as social communicative signal. However, a broader definition of *social* forms of disgust could also include social disgusting words and social disgusting scenes. While the common and separable neural bases of these social disgust aspects are of high interest for future research the low number of studies employing these stimuli does currently not allow a robust meta-analytic examination. Third, the included studies on social disgust investigated perception rather than production of disgusted facial expression. Although the number of facial expression production fMRI studies for disgust is currently too small to warrant a meta-analytic examination, determining common and separable neural correlates of production and perception would be of interest for future studies. Fourth, the present study focused on core and social processing in the domain of disgust. Previous original studies have employed experimental designs to compare core and social disgust processing but have been limited due to limitations inherent to original studies (Schäfer et al., 2005; Schienle et al., 2020). Previous meta-analyses have employed a similar strategy in the domain of empathy processing by comparing meta-analytic activity maps for facial pain expressions and stimuli presenting acute pain infliction (Timmers et al., 2018). While future studies may extend this approach to other emotional domains such as fear and encompass emotional domain as additional dimension of common and shared neural representations in neuroimaging meta-analysis, the conventional fMRI and meta-analytic approach is limited with respect to determining fine-grained separable neurofunctional representations and multivariate representational models may help to further determine separable process-specific neurofunctional representations (see e.g., Zhou et al., 2020; Zhou et al., 2021). Fifth, the present database of original studies did not allow to examine sex-differences in the neural basis of disgust processing. Initial studies reported e.g., stronger disgust-related inferior frontal reactivity in women (Caseras et al., 2007), while other studies did not find sex differences (Baumann and Mattingley, 2012; Schienle et al., 2005a; Stark et al., 2003). Lastly, the present study included task-based fMRI studies focusing on regional analyses, while recent advanced approaches and findings suggest a distributed neural representation (Zhou et al., 2021) or a functional division between gyri and sulci (Jiang et al., 2015; Jiang et al., 2018; Jiang et al., 2021) of cognitive and emotional processes. Future studies may capitalize on these developments to further dissect common and differential neural representation of social and non-social processes or different defensive responses such as disgust and fear.

## 5. Conclusions

The present study employed a coordinate-based neuroimaging meta-analytic approach to determine the neurobiological basis of disgust processing in humans. Exposure to disgust eliciting stimuli induced robust activity in a network encompassing regions engaged in early visual threat detection, emotional experience and defensive motor responses, while social stimuli of disgust primarily engaged regions engaged in facial and social cognitive processes. Across both disgust domains, the anterior insula/inferior frontal gyrus and the fusiform gyrus were robustly engaged suggesting an involvement of these regions across disgust processing modalities.

## Acknowledgements

This work was supported by the National Key Research and Development Program of China (2018YFA0701400); and the National Natural Science Foundation of China (NSFC, 61871420).

We would also like to thank Zonglin He and Qian Tao from Jinan University (China) for their kind help with resolving some data analyzing issues.

## References

Adolphs, R., 2002. Recognizing emotion from facial expressions: psychological and neurological mechanisms. Behav. Cogn. Neurosci. Rev. 1, 21–62. https://doi.org/10.1177/1534582302001001003.

Adolphs, R., Damasio, H., Tranel, D., Damasio, A.R., 1996. Cortical systems for the recognition of emotion in facial expressions. J. Neurosci. 16, 7678–7687. https://doi.org/10.1523/JNEUROSCI.16-23-07678.1996.

Adolphs, R., Gosselin, F., Buchanan, T.W., Tranel, D., Schyns, P., Damasio, A.R., 2005. A mechanism for impaired fear recognition after amygdala damage. Nature 433, 68–72. https://doi.org/10.1038/nature03086.

Adolphs, R., Tranel, D., Damasio, A.R., 2003. Dissociable neural systems for recognizing emotions. Brain Cogn. 52, 61–69. https://doi.org/10.1016/S0278-2626(03)00009-5.

Ahn, W.-Y., Kishida, K.T., Gu, X., Lohrenz, T., Harvey, A., Alford, J.R., Smith, K.B., Yaffe, G., Hibbing, J.R., Dayan, P., Montague, P.R., 2014. Nonpolitical images evoke neural predictors of political ideology. Curr. Biol. 24, 2693–2699. https://doi.org/10.1016/j.cub.2014.09.050.

Alain, C., Du, Y., Bernstein, L.J., Barten, T., Banai, K., 2018. Listening under difficult conditions: an activation likelihood estimation meta-analysis. Hum. Brain Mapp. 39, 2695–2709. https://doi.org/10.1002/hbm.24031.

Amting, J.M., Greening, S.G., Mitchell, D.G.V., 2010. Multiple mechanisms of consciousness: the neural correlates of emotional awareness. J. Neurosci. 30, 10039–10047. https://doi.org/10.1523/JNEUROSCI.6434-09.2010.

Andersen, R.A., 2011. Inferior parietal lobule function in spatial perception and visuomotor integration, Comprehensive Physiology. Wiley Online Library, pp. 483–518.

Ardila, A., Bernal, B., Rosselli, M., 2015. Language and visual perception associations: meta-analytic connectivity modeling of Brodmann area 37. Behav. Neurol. 2015, 565871. https://doi.org/10.1155/2015/565871.

Ashworth, F., Pringle, A., Norbury, R., Harmer, C.J., Cowen, P.J., Cooper, M.J., 2011. Neural response to angry and disgusted facial expressions in bulimia nervosa. Psychol. Med. 41, 2375–2384. https://doi.org/10.1017/S0033291711000626.

Baumann, O., Mattingley, J.B., 2012. Functional topography of primary emotion processing in the human cerebellum. Neuroimage 61, 805–811. https://doi.org/10.1016/j.neuroimage.2012.03.044.

Becker, B., Mihov, Y., Scheele, D., Kendrick, K.M., Feinstein, J.S., Matusch, A., Aydin, M., Reich, H., Urbach, H., Oros-Peusquens, A.-M., Shah, N.J., Kunz, W.S., Schlaepfer, T.E., Zilles, K., Maier, W., Hurlemann, R., 2012. Fear processing and social networking in the absence of a functional amygdala. Biol. Psychiatry 72, 70–77. https://doi.org/10.1016/j.biopsych.2011.11.024.

Belyk, M., Brown, S., Lim, J., Kotz, S.A., 2017. Convergence of semantics and emotional expression within the IFG pars orbitalis. Neuroimage 156, 240–248. https://doi.org/10.1016/j.neuroimage.2017.04.020.

Benuzzi, F., Lui, F., Duzzi, D., Nichelli, P., Porro, C., 2008. Does it look painful or disgusting? Ask your parietal and cingulate cortex. J. Neurosci. 28, 923–931. https://doi.org/10.1523/JNEUROSCI.4012-07.2008.

Berlin, H.A., Stern, E.R., Ng, J., Zhang, S., Rosenthal, D., Turetzky, R., Tang, C., Goodman, W., 2017. Altered olfactory processing and increased insula activity in patients with obsessive-compulsive disorder: an fMRI study. Psychiatry Res Neuroimaging 262, 15–24. https://doi.org/10.1016/j.pscychresns.2017.01.012.

Borg, C., de Jong, P.J., Renken, R.J., Georgiadis, J.R., 2012. Disgust trait modulates frontal-posterior coupling as a function of disgust domain. Soc. Cogn. Affect. Neurosci. 8, 351–358. https://doi.org/10.1093/scan/nss006

Bradley, M.M., Sabatinelli, D., Lang, P.J., Fitzsimmons, J.R., King, W., Desai, P., 2003. Activation of the visual cortex in motivated attention. Behav. Neurosci. 117, 369–380. https://doi.org/10.1037/0735-7044.117.2.369.

Britton, J.C., Taylor, S.F., Sudheimer, K.D., Liberzon, I., 2006. Facial expressions and complex IAPS pictures: common and differential networks. Neuroimage 31, 906–919. https://doi.org/10.1016/j.neuroimage.2005.12.050.

Burklund, L., Eisenberger, N., Lieberman, M., 2007. The face of rejection: rejection sensitivity moderates dorsal anterior cingulate activity to disapproving facial expressions. Soc. Neurosci. 2, 238–253. https://doi.org/10.1080/17470910701391711.

Buxbaum, L.J., Randerath, J., 2018. Chapter 17 - Limb apraxia and the left parietal lobe, in: Vallar, G., Coslett, H.B. (Eds.), Handbook of Clinical Neurology. Elsevier, pp. 349–363.

Bzdok, D., Schilbach, L., Vogeley, K., Schneider, K., Laird, A.R., Langner, R., Eickhoff, S.B., 2012. Parsing the neural correlates of moral cognition: ALE meta-analysis on morality, theory of mind, and empathy. Brain Struct. Funct. 217, 783–796. https://doi.org/10.1007/s00429-012-0380-y.

Calder, A.J., Keane, J., Manes, F., Antoun, N., Young, A.W., 2000. Impaired recognition and experience of disgust following brain injury. Nat. Neurosci. 3, 1077–1078. https://doi.org/10.1038/80586.

Calder, A.J., Lawrence, A.D., Young, A.W., 2001. Neuropsychology of fear and loathing. Nat. Rev. Neurosci. 2, 352–363. https://doi.org/10.1038/35072584.

Calder, A.J., Young, A.W., Rowland, D., Perrett, D.I., Hodges, J.R., Etcoff, N.L., 1996. Facial emotion recognition after bilateral amygdala damage: differentially severe impairment of fear. Cogn. Neuropsychol. 13, 699–745. https://doi.org/10.1080/026432996381890.

Cantone, M., Lanza, G., Bella, R., Pennisi, G., Santalucia, P., Bramanti, P., Pennisi, M., 2019. Fear and disgust: case report of two uncommon emotional disturbances evoked by visual disperceptions after a right temporal-insular stroke. BMC Neurol. 19, 193. https://doi.org/10.1186/s12883-019-1417-0.

Caseras, X., Mataix-Cols, D., An, S.K., Lawrence, N.S., Speckens, A., Giampietro, V., Brammer, M.J., Phillips, M.L., 2007. Sex differences in neural responses to disgusting visual stimuli: implications for disgust-related psychiatric disorders. Biol. Psychiatry 62, 464–471. https://doi.org/10.1016/j.biopsych.2006.10.030.

Cechetto, D.F., Topolovec, J.C., 2002. Cerebral cortex, in: Ramachandran, V.S. (Ed.), Encyclopedia of the Human Brain. Academic Press, New York, pp. 663–679.

Chakrabarti, B., Bullmore, E., Baron-Cohen, S., 2006. Empathizing with basic emotions: common and discrete neural substrates. Soc. Neurosci. 1, 364–384. https://doi.org/10.1080/17470910601041317.

Chapman, H.A., Anderson, A.K., 2012. Understanding disgust. Ann. N. Y. Acad. Sci. 1251, 62–76. https://doi.org/10.1111/j.1749-6632.2011.06369.x.

Chapman, H.A., Kim, D.A., Susskind, J.M., Anderson, A.K., 2009. In bad taste: evidence for the oral origins of moral disgust. Science 323, 1222–1226. https://doi.org/10.1126/science.1165565.

Chen, K.-H., Hua, A.Y., Lwi, S.J., Haase, C.M., Rosen, H.J., Miller, B.L., Levenson, R.W., 2021. Smaller volume in left-lateralized brain structures correlates with greater experience of negative non-target emotions in neurodegenerative diseases. Cereb. Cortex 31, 15–31. https://doi.org/10.1093/cercor/bhaa193.

Chen, T., Becker, B., Camilleri, J., Wang, L., Yu, S., Eickhoff, S.B., Feng, C., 2018. A domain-general brain network underlying emotional and cognitive interference processing: evidence from coordinate-based and functional connectivity meta-analyses. Brain Struct. Funct. 223, 3813–3840. https://doi.org/10.1007/s00429-018-1727-9.

Cloutman, L.L., Binney, R.J., Drakesmith, M., Parker, G.J.M., Lambon Ralph, M.A., 2012. The variation of function across the human insula mirrors its patterns of structural connectivity: evidence from in vivo probabilistic tractography. Neuroimage 59, 3514–3521. https://doi.org/10.1016/j.neuroimage.2011.11.016.

Corradi-Dell’Acqua, C., Tusche, A., Vuilleumier, P., Singer, T., 2016. Cross-modal representations of first-hand and vicarious pain, disgust and fairness in insular and cingulate cortex. Nat. Commun. 7, 10904. https://doi.org/10.1038/ncomms10904.

Costafreda, S.G., Brammer, M.J., David, A.S., Fu, C.H.Y., 2008. Predictors of amygdala activation during the processing of emotional stimuli: a meta-analysis of 385 PET and fMRI studies. Brain Res. Rev. 58, 57–70. https://doi.org/10.1016/j.brainresrev.2007.10.012.

Cox, C.L., Gotimer, K., Roy, A.K., Castellanos, F.X., Milham, M.P., Kelly, C., 2010. Your resting brain cares about your risky behavior. PLoS ONE 5, e12296. https://doi.org/10.1371/journal.pone.0012296.

Craig, A.D., 2009. How do you feel — now? The anterior insula and human awareness. Nat. Rev. Neurosci. 10, 59–70. https://doi.org/10.1038/nrn2555.

Critchley, H.D., Wiens, S., Rotshtein, P., Öhman, A., Dolan, R.J., 2004. Neural systems supporting interoceptive awareness. Nat. Neurosci. 7, 189–195. https://doi.org/10.1038/nn1176.

Curtis, V., de Barra, M., Aunger, R., 2011. Disgust as an adaptive system for disease avoidance behaviour. Phil. Trans. R. Soc. B 366, 389–401. https://doi.org/10.1098/rstb.2010.0117.

Darwin, C., 1965. The expression of emotion in man and animals. University of Chicago Press: Chicago, IL.

Davey, G.C.L., 2011. Disgust: the disease-avoidance emotion and its dysfunctions. Phil. Trans. R. Soc. B 366, 3453–3465. https://doi.org/10.1098/rstb.2011.0039.

Davidson, R.J., Irwin, W., 1999. The functional neuroanatomy of emotion and affective style. Trends Cogn. Sci. 3, 11–21. https://doi.org/10.1016/s1364-6613(98)01265-0.

Deeley, Q., Daly, E.M., Azuma, R., Surguladze, S., Giampietro, V., Brammer, M.J., Hallahan, B., Dunbar, R.I.M., Phillips, M.L., Murphy, D.G.M., 2008. Changes in male brain responses to emotional faces from adolescence to middle age. Neuroimage 40, 389–397. https://doi.org/10.1016/j.neuroimage.2007.11.023.

Deeley, Q., Daly, E.M., Surguladze, S., Page, L., Toal, F., Robertson, D., Curran, S., Giampietro, V., Seal, M., Brammer, M.J., Andrew, C., Murphy, K., Phillips, M.L., Murphy, D.G.M., 2007. An event related functional magnetic resonance imaging study of facial emotion processing in asperger syndrome. Biol. Psychiatry 62, 207–217. https://doi.org/10.1016/j.biopsych.2006.09.037.

Derntl, B., Habel, U., Windischberger, C., Robinson, S., Kryspin-Exner, I., Gur, R.C., Moser, E., 2009. General and specific responsiveness of the amygdala during explicit emotion recognition in females and males. BMC Neurosci. 10, 91. https://doi.org/10.1186/1471-2202-10-91.

Desimone, R., Albright, T.D., Gross, C.G., Bruce, C., 1984. Stimulus-selective properties of inferior temporal neurons in the macaque. The Journal of Neuroscience 4, 2051–2062. https://doi.org/10.1523/JNEUROSCI.04-08-02051.1984.

Dosenbach, N.U.F., Visscher, K.M., Palmer, E.D., Miezin, F.M., Wenger, K.K., Kang, H.C., Burgund, E.D., Grimes, A.L., Schlaggar, B.L., Petersen, S.E., 2006. A core system for the implementation of task sets. Neuron 50, 799–812. https://doi.org/10.1016/j.neuron.2006.04.031.

Eickhoff, S.B., Bzdok, D., Laird, A.R., Kurth, F., Fox, P.T., 2012. Activation likelihood estimation meta-analysis revisited. Neuroimage 59, 2349–2361. https://doi.org/10.1016/j.neuroimage.2011.09.017.

Eickhoff, S.B., Bzdok, D., Laird, A.R., Roski, C., Caspers, S., Zilles, K., Fox, P.T., 2011. Co-activation patterns distinguish cortical modules, their connectivity and functional differentiation. Neuroimage 57, 938–949. https://doi.org/10.1016/j.neuroimage.2011.05.021.

Eickhoff, S.B., Laird, A.R., Fox, P.M., Lancaster, J.L., Fox, P.T., 2017. Implementation errors in the GingerALE software: description and recommendations. Hum. Brain Mapp. 38, 7–11. https://doi.org/10.1002/hbm.23342.

Eickhoff, S.B., Laird, A.R., Grefkes, C., Wang, L.E., Zilles, K., Fox, P.T., 2009. Coordinate-based activation likelihood estimation meta-analysis of neuroimaging data: a random-effects approach based on empirical estimates of spatial uncertainty. Hum. Brain Mapp. 30, 2907–2926. https://doi.org/10.1002/hbm.20718.

Eickhoff, S.B., Nichols, T.E., Laird, A.R., Hoffstaedter, F., Amunts, K., Fox, P.T., Bzdok, D., Eickhoff, C.R., 2016. Behavior, sensitivity, and power of activation likelihood estimation characterized by massive empirical simulation. Neuroimage 137, 70–85. https://doi.org/10.1016/j.neuroimage.2016.04.072.

Ekman, P., 1993. Facial expression and emotion. Am. Psychol. 48, 384–392. https://doi.org/10.1037//0003-066x.48.4.384.

Fairhall, S.L., Ishai, A., 2007. Effective connectivity within the distributed cortical network for face perception. Cereb. Cortex 17, 2400–2406. https://doi.org/10.1093/cercor/bhl148.

Fan, Y., Duncan, N.W., de Greck, M., Northoff, G., 2011. Is there a core neural network in empathy? An fMRI based quantitative meta-analysis. Neurosci. Biobehav. Rev. 35, 903–911. https://doi.org/10.1016/j.neubiorev.2010.10.009.

Feinstein, J.S., Adolphs, R., Damasio, A., Tranel, D., 2011. The human amygdala and the induction and experience of fear. Curr. Biol. 21, 34–38. https://doi.org/10.1016/j.cub.2010.11.042.

Feng, C., Becker, B., Huang, W., Wu, X., Eickhoff, S.B., Chen, T., 2018. Neural substrates of the emotion-word and emotional counting Stroop tasks in healthy and clinical populations: a meta-analysis of functional brain imaging studies. Neuroimage 173, 258–274. https://doi.org/10.1016/j.neuroimage.2018.02.023.

Fitzgerald, D.A., Angstadt, M., Jelsone, L.M., Nathan, P.J., Phan, K.L., 2006. Beyond threat: amygdala reactivity across multiple expressions of facial affect. Neuroimage 30, 1441–1448. https://doi.org/10.1016/j.neuroimage.2005.11.003.

Frith, C.D., 2007. The social brain? Phil. Trans. R. Soc. B 362, 671–678. https://doi.org/10.1098/rstb.2006.2003.

Fusar-Poli, P., Placentino, A., Carletti, F., Allen, P., Landi, P., Abbamonte, M., Barale, F., Perez, J., McGuire, P., Politi, P.L., 2009a. Laterality effect on emotional faces processing: ALE meta-analysis of evidence. Neurosci. Lett. 452, 262–267. https://doi.org/10.1016/j.neulet.2009.01.065.

Fusar-Poli, P., Placentino, A., Carletti, F., Landi, P., Allen, P., Surguladze, S., Benedetti, F., Abbamonte, M., Gasparotti, R., Barale, F., Pérez, J., McGuire, P., Politi, P., 2009b. Functional atlas of emotional faces processing: a voxel-based meta-analysis of 105 functional magnetic resonance imaging studies. J. Psychiatry Neurosci. 34, 418–432. https://doi.org/10.1111/j.1365-2850.2009.01434.x.

Gallese, V., Goldman, A., 1998. Mirror neurons and the simulation theory of mind-reading. Trends Cogn. Sci. 2, 493–501. https://doi.org/10.1016/S1364-6613(98)01262-5.

Gallese, V., Sinigaglia, C., 2011. What is so special about embodied simulation? Trends Cogn. Sci. 15, 512–519. https://doi.org/10.1016/j.tics.2011.09.003.

Gallese, V., Sinigaglia, C., 2012. Response to de Bruin and Gallagher: embodied simulation as reuse is a productive explanation of a basic form of mind-reading. Trends Cogn. Sci. 16, 99–100. https://doi.org/10.1016/j.tics.2011.12.002.

Giovanniello, J., Yu, K., Furlan, A., Nachtrab, G.T., Sharma, R., Chen, X., Li, B., 2020. A central amygdala-globus pallidus circuit conveys unconditioned stimulus-related information and controls fear learning. J. Neurosci. 40, 9043–9054. https://doi.org/10.1523/JNEUROSCI.2090-20.2020.

Gorno-Tempini, M.L., Pradelli, S., Serafini, M., Pagnoni, G., Baraldi, P., Porro, C., Nicoletti, R., Umità, C., Nichelli, P., 2001. Explicit and incidental facial expression processing: an fMRI study. Neuroimage 14, 465–473. https://doi.org/10.1006/nimg.2001.0811.

Güntürkün, O., Ströckens, F., Ocklenburg, S., 2020. Brain lateralization: a comparative perspective. Physiol. Rev. 100, 1019–1063. https://doi.org/10.1152/physrev.00006.2019.

Hadjistavropoulos, T., Craig, K.D., Duck, S., Cano, A., Goubert, L., Jackson, P.L., Mogil, J.S., Rainville, P., Sullivan, M.J.L., Williams, A.C.C., Vervoort, T., Fitzgerald, T.D., 2011. A biopsychosocial formulation of pain communication. Psychol. Bull. 137, 910–939. https://doi.org/10.1037/a0023876.

Haidt, J., McCauley, C., Rozin, P., 1994. Individual differences in sensitivity to disgust: a scale sampling seven domains of disgust elicitors. Pers. Individ. Differ. 16, 701–713. https://doi.org/10.1016/0191-8869(94)90212-7.

Harrison, N.A., Gray, M.A., Gianaros, P.J., Critchley, H.D., 2010. The embodiment of emotional feelings in the brain. J. Neurosci. 30, 12878–12884. https://doi.org/10.1523/JNEUROSCI.1725-10.2010.

Haxby, J.V., Hoffman, E.A., Gobbini, M.I., 2000. The distributed human neural system for face perception. Trends Cogn. Sci. 4, 223–233. https://doi.org/10.1016/S1364-6613(00)01482-0.

Heining, M., Young, A., Ioannou, G., Andrew, C., Brammer, M., Gray, J., Phillips, M., 2004. Disgusting smells activate human anterior insula and ventral striatum. Ann. N. Y. Acad. Sci. 1000, 380–384. https://doi.org/10.1196/annals.1280.035.

Herrington, J.D., Taylor, J.M., Grupe, D.W., Curby, K.M., Schultz, R.T., 2011. Bidirectional communication between amygdala and fusiform gyrus during facial recognition. Neuroimage 56, 2348–2355. https://doi.org/10.1016/j.neuroimage.2011.03.072.

Holtmann, O., Bruchmann, M., Mönig, C., Schwindt, W., Melzer, N., Miltner, W.H.R., Straube, T., 2020. Lateralized deficits of disgust processing after insula-basal ganglia damage. Front. Psychol. 11, 1429. https://doi.org/10.3389/fpsyg.2020.01429.

Jabbi, M., Bastiaansen, J., Keysers, C., 2008. A common anterior insula representation of disgust observation, experience and imagination shows divergent functional connectivity pathways. PLoS ONE 3, e2939. https://doi.org/10.1371/journal.pone.0002939.

Jiang, X., Li, X., Lv, J., Zhang, T., Zhang, S., Guo, L., Liu, T., 2015. Sparse representation of HCP grayordinate data reveals novel functional architecture of cerebral cortex. Hum. Brain Mapp. 36, 5301–5319. https://doi.org/10.1002/hbm.23013.

Jiang, X., Li, X., Lv, J., Zhao, S., Zhang, S., Zhang, W., Zhang, T., Han, J., Guo, L., Liu, T., 2018. Temporal dynamics assessment of spatial overlap pattern of functional brain networks reveals novel functional architecture of cerebral cortex. IEEE Trans. Biomed. Eng. 65, 1183–1192. https://doi.org/10.1109/TBME.2016.2598728.

Jiang, X., Zhang, T., Zhang, S., Kendrick, K.M., Liu, T., 2021. Fundamental functional differences between gyri and sulci: implications for brain function, cognition, and behavior. Psychoradiology 1, 23–41. https://doi.org/10.1093/psyrad/kkab005.

Kipps, C.M., Duggins, A.J., McCusker, E.A., Calder, A.J., 2007. Disgust and happiness recognition correlate with anteroventral insula and amygdala volume respectively in preclinical Huntington’s disease. J. Cogn. Neurosci. 19, 1206–1217. https://doi.org/10.1162/jocn.2007.19.7.1206.

Kirby, L.A.J., Robinson, J.L., 2017. Affective mapping: an activation likelihood estimation (ALE) meta-analysis. Brain Cogn. 118, 137–148. https://doi.org/10.1016/j.bandc.2015.04.006.

Klugah-Brown, B., Di, X., Zweerings, J., Mathiak, K., Becker, B., Biswal, B., 2020. Common and separable neural alterations in substance use disorders: a coordinate-based meta-analyses of functional neuroimaging studies in humans. Hum. Brain Mapp. 41, 4459–4477. https://doi.org/10.1002/hbm.25085.

Klugah-Brown, B., Zhou, X., Pradhan, B.K., Zweerings, J., Mathiak, K., Biswal, B., Becker, B., 2021. Common neurofunctional dysregulations characterize obsessive–compulsive, substance use, and gaming disorders -an activation likelihood meta-analysis of functional imaging studies. Addict. Biol. 26, e12997. https://doi.org/10.1111/adb.12997.

Kohn, N., Eickhoff, S.B., Scheller, M., Laird, A.R., Fox, P.T., Habel, U., 2014. Neural network of cognitive emotion regulation - an ALE meta-analysis and MACM analysis. Neuroimage 87, 345–355. https://doi.org/10.1016/j.neuroimage.2013.11.001.

Kotkowski, E., Price, L.R., Mickle Fox, P., Vanasse, T.J., Fox, P.T., 2018. The hippocampal network model: a transdiagnostic metaconnectomic approach. Neuroimage Clin. 18, 115–129. https://doi.org/10.1016/j.nicl.2018.01.002.

Krolak-Salmon, P., Hénaff, M.-A., Isnard, J., Tallon-Baudry, C., Guénot, M., Vighetto, A., Bertrand, O., Mauguière, F., 2003. An attention modulated response to disgust in human ventral anterior insula. Ann. Neurol. 53, 446–453. https://doi.org/10.1002/ana.10502.

Kupfer, T.R., Fessler, D.M.T., 2018. Ectoparasite defence in humans: relationships to pathogen avoidance and clinical implications. Phil. Trans. R. Soc. B 373, 20170207. https://doi.org/doi:10.1098/rstb.2017.0207.

Laird, A.R., Eickhoff, S.B., Li, K., Robin, D.A., Glahn, D.C., Fox, P.T., 2009. Investigating the functional heterogeneity of the default mode network using coordinate-based meta-analytic modeling. J. Neurosci. 29, 14496–14505. https://doi.org/10.1523/JNEUROSCI.4004-09.2009.

Laird, A.R., Fox, P.M., Price, C.J., Glahn, D.C., Uecker, A.M., Lancaster, J.L., Turkeltaub, P.E., Kochunov, P., Fox, P.T., 2005. ALE meta-analysis: controlling the false discovery rate and performing statistical contrasts. Hum. Brain Mapp. 25, 155–164. https://doi.org/10.1002/hbm.20136.

Laird, A.R., Robinson, J.L., McMillan, K.M., Tordesillas-Gutiérrez, D., Moran, S.T., Gonzales, S.M., Ray, K.L., Franklin, C., Glahn, D.C., Fox, P.T., Lancaster, J.L., 2010. Comparison of the disparity between Talairach and MNI coordinates in functional neuroimaging data: validation of the Lancaster transform. Neuroimage 51, 677–683. https://doi.org/10.1016/j.neuroimage.2010.02.048.

Lancaster, J.L., Tordesillas-Gutiérrez, D., Martinez, M., Salinas, F., Evans, A., Zilles, K., Mazziotta, J.C., Fox, P.T., 2007. Bias between MNI and Talairach coordinates analyzed using the ICBM-152 brain template. Hum. Brain Mapp. 28, 1194–1205. https://doi.org/10.1002/hbm.20345.

Langner, R., Rottschy, C., Laird, A.R., Fox, P.T., Eickhoff, S.B., 2014. Meta-analytic connectivity modeling revisited: controlling for activation base rates. Neuroimage 99, 559–570. https://doi.org/10.1016/j.neuroimage.2014.06.007.

Larson, C.L., Aronoff, J., Sarinopoulos, I.C., Zhu, D.C., 2009. Recognizing threat: a simple geometric shape activates neural circuitry for threat detection. J. Cogn. Neurosci. 21, 1523–1535. https://doi.org/10.1162/jocn.2009.21111.

Lassalle, A., Zürcher, N.R., Porro, C.A., Benuzzi, F., Hippolyte, L., Lemonnier, E., Åsberg Johnels, J., Hadjikhani, N., 2019. Influence of anxiety and alexithymia on brain activations associated with the perception of others’ pain in autism. Soc. Neurosci. 14, 359–377. https://doi.org/10.1080/17470919.2018.1468358.

LeDoux, J.E., 2014. Coming to terms with fear. Proc. Natl. Acad. Sci. U. S. A. 111, 2871. https://doi.org/10.1073/pnas.1400335111.

LeDoux, J.E., Pine, D.S., 2016. Using neuroscience to help understand fear and anxiety: a two-system framework. Am. J. Psychiatry 173, 1083–1093. https://doi.org/10.1176/appi.ajp.2016.16030353.

Li, J., Xu, L., Zheng, X., Fu, M., Zhou, F., Xu, X., Ma, X., Li, K., Kendrick, K.M., Becker, B., 2019. Common and dissociable contributions of alexithymia and autism to domain-specific interoceptive dysregulations: a dimensional neuroimaging approach. Psychother. Psychosom. 88, 187–189. https://doi.org/10.1159/000495122.

Lindquist, D.H., 2020. Emotion in motion: a three-stage model of aversive classical conditioning. Neurosci. Biobehav. Rev. 115, 363–377. https://doi.org/10.1016/j.neubiorev.2020.04.025.

Liu, M., Liu, C.H., Zheng, S., Zhao, K., Fu, X., 2021. Reexamining the neural network involved in perception of facial expression: a meta-analysis. Neurosci. Biobehav. Rev. 131, 179–191. https://doi.org/10.1016/j.neubiorev.2021.09.024.

Liu, Z., Rolls, E.T., Liu, Z., Zhang, K., Yang, M., Du, J., Gong, W., Cheng, W., Dai, F., Wang, H., Ugurbil, K., Zhang, J., Feng, J., 2019. Brain annotation toolbox: exploring the functional and genetic associations of neuroimaging results. Bioinformatics 35, 3771–3778. https://doi.org/10.1093/bioinformatics/btz128.

Malhi, G., Lagopoulos, J., Sachdev, P., Ivanovski, B., Shnier, R., Ketter, T., 2007. Is a lack of disgust something to fear? An fMRI facial recognition study in euthymic bipolar disorder patients. Bipolar Disord. 9, 345–357. https://doi.org/10.1111/j.1399-5618.2007.00485.x.

Mattavelli, G., Sormaz, M., Flack, T., Asghar, A.U.R., Fan, S., Frey, J., Manssuer, L., Usten, D., Young, A.W., Andrews, T.J., 2014. Neural responses to facial expressions support the role of the amygdala in processing threat. Soc. Cogn. Affect. Neurosci. 9, 1684–1689. https://doi.org/10.1093/scan/nst162.

Meier, L., Friedrich, H., Federspiel, A., Jann, K., Morishima, Y., Landis, B.N., Wiest, R., Strik, W., Dierks, T., 2015. Rivalry of homeostatic and sensory-evoked emotions: dehydration attenuates olfactory disgust and its neural correlates. Neuroimage 114, 120–127. https://doi.org/10.1016/j.neuroimage.2015.03.048.

Meier, S.K., Ray, K.L., Mastan, J.C., Salvage, S.R., Robin, D.A., 2021. Meta-analytic connectivity modelling of deception-related brain regions. PLoS ONE 16, e0248909. https://doi.org/10.1371/journal.pone.0248909.

Mesulam, M.M., Mufson, E.J., 1982a. Insula of the old world monkey. I: architectonics in the insulo-orbito-temporal component of the paralimbic brain. J. Comp. Neurol. 212, 1–22. https://doi.org/10.1002/cne.902120102.

Mesulam, M.M., Mufson, E.J., 1982b. Insula of the old world monkey. III: efferent cortical output and comments on function. J. Comp. Neurol. 212, 38–52. https://doi.org/10.1002/cne.902120104.

Mihov, Y., Kendrick, K.M., Becker, B., Zschernack, J., Reich, H., Maier, W., Keysers, C., Hurlemann, R., 2013. Mirroring fear in the absence of a functional amygdala. Biol. Psychiatry 73, e9–e11. https://doi.org/10.1016/j.biopsych.2012.10.029.

Molapour, T., Golkar, A., Navarrete, C.D., Haaker, J., Olsson, A., 2015. Neural correlates of biased social fear learning and interaction in an intergroup context. Neuroimage 121, 171–183. https://doi.org/10.1016/j.neuroimage.2015.07.015.

Morawetz, C., Bode, S., Baudewig, J., Heekeren, H.R., 2017. Effective amygdala-prefrontal connectivity predicts individual differences in successful emotion regulation. Soc. Cogn. Affect. Neurosci. 12, 569–585. https://doi.org/10.1093/scan/nsw169.

Mueller, E.M., Pizzagalli, D.A., 2016. One-year-old fear memories rapidly activate human fusiform gyrus. Soc. Cogn. Affect. Neurosci. 11, 308–316. https://doi.org/10.1093/scan/nsv122.

Müller, V.I., Cieslik, E.C., Laird, A.R., Fox, P.T., Radua, J., Mataix-Cols, D., Tench, C.R., Yarkoni, T., Nichols, T.E., Turkeltaub, P.E., Wager, T.D., Eickhoff, S.B., 2018. Ten simple rules for neuroimaging meta-analysis. Neurosci. Biobehav. Rev. 84, 151–161. https://doi.org/10.1016/j.neubiorev.2017.11.012.

Murphy, F.C., Nimmo-Smith, I., Lawrence, A.D., 2003. Functional neuroanatomy of emotions: a meta-analysis. Cogn. Affect. Behav. Neurosci. 3, 207–233. https://doi.org/10.3758/cabn.3.3.207.

Oaten, M., Stevenson, R.J., Case, T.I., 2009. Disgust as a disease-avoidance mechanism. Psychol. Bull. 135, 303–321. https://doi.org/10.1037/a0014823.

Olatunji, B.O., Sawchuk, C., 2005. Disgust: characteristic features, social manifestations, and clinical implications. J. Soc. Clin. Psychol. 24, 932–962. https://doi.org/10.1521/jscp.2005.24.7.932.

Olatunji, B.O., Williams, N.L., Tolin, D.F., Abramowitz, J.S., Sawchuk, C.N., Lohr, J.M., Elwood, L.S., 2007. The disgust scale: item analysis, factor structure, and suggestions for refinement. Psychol. Assess. 19, 281–297. https://doi.org/10.1037/1040-3590.19.3.281.

Oliveri, M., Babiloni, C., Filippi, M.M., Caltagirone, C., Babiloni, F., Cicinelli, P., Traversa, R., Palmieri, M.G., Rossini, P.M., 2003. Influence of the supplementary motor area on primary motor cortex excitability during movements triggered by neutral or emotionally unpleasant visual cues. Exp. Brain Res. 149, 214–221. https://doi.org/10.1007/s00221-002-1346-8.

Page, M.J., Moher, D., Bossuyt, P.M., Boutron, I., Hoffmann, T.C., Mulrow, C.D., Shamseer, L., Tetzlaff, J.M., Akl, E.A., Brennan, S.E., Chou, R., Glanville, J., Grimshaw, J.M., Hróbjartsson, A., Lalu, M.M., Li, T., Loder, E.W., Mayo-Wilson, E., McDonald, S., McGuinness, L.A., Stewart, L.A., Thomas, J., Tricco, A.C., Welch, V.A., Whiting, P., McKenzie, J.E., 2021. PRISMA 2020 explanation and elaboration: updated guidance and exemplars for reporting systematic reviews. BMJ 372, n160. https://doi.org/10.1136/bmj.n160.

Palomero-Gallagher, N., Amunts, K., 2021. A short review on emotion processing: a lateralized network of neuronal networks. Brain Struct. Funct. https://doi.org/10.1007/s00429-021-02331-7.

Papitto, G., Friederici, A.D., Zaccarella, E., 2020. The topographical organization of motor processing: an ALE meta-analysis on six action domains and the relevance of Broca’s region. Neuroimage 206, 116321. https://doi.org/10.1016/j.neuroimage.2019.116321.

Paradiso, S., Robinson, R.G., Andreasen, N.C., Downhill, J.E., Davidson, R.J., Kirchner, P.T., Watkins, G.L., Ponto, L.L., Hichwa, R.D., 1997. Emotional activation of limbic circuitry in elderly normal subjects in a PET study. Am. J. Psychiatry 154, 384–389. https://doi.org/10.1176/ajp.154.3.384.

Perrett, D.I., Rolls, E.T., Caan, W., 1982. Visual neurones responsive to faces in the monkey temporal cortex. Exp. Brain Res. 47, 329–342. https://doi.org/10.1007/BF00239352.

Phan, K.L., Wager, T., Taylor, S.F., Liberzon, I., 2002. Functional neuroanatomy of emotion: a meta-analysis of emotion activation studies in PET and fMRI. Neuroimage 16, 331–348. https://doi.org/10.1006/nimg.2002.1087.

Phelps, E.A., LeDoux, J.E., 2005. Contributions of the amygdala to emotion processing: from animal models to human behavior. Neuron 48, 175–187. https://doi.org/10.1016/j.neuron.2005.09.025.

Phillips, M., Marks, I., Senior, C., Lythgoe, D., O’Dwyer, A., Meehan, O., Williams, S., Brammer, M., Bullmore, E., McGuire, P., 2000. A differential neural response in obsessive-compulsive disorder patients with washing compared with checking symptoms to disgust. Psychol. Med. 30, 1037–1050. https://doi.org/10.1017/S0033291799002652.

Phillips, M., Williams, L., Heining, M., Herba, C., Russell, T., Andrew, C., Bullmore, E., Brammer, M., Williams, S., Morgan, M., Young, A., Gray, J., 2004. Differential neural responses to overt and covert presentations of facial expressions of fear and disgust. Neuroimage 21, 1484–1496. https://doi.org/10.1016/j.neuroimage.2003.12.013.

Phillips, M., Williams, L., Senior, C., Bullmore, E.T., Brammer, M.J., Andrew, C., Williams, S.C.R., David, A.S., 1999. A differential neural response to threatening and non-threatening negative facial expressions in paranoid and non-paranoid schizophrenics. Psychiatry Res Neuroimaging 92, 11–31. https://doi.org/10.1016/S0925-4927(99)00031-1.

Phillips, M., Young, A.W., Scott, S.K., Calder, A.J., Andrew, C., Giampietro, V., Williams, S.C.R., Bullmore, E.T., Brammer, M., Gray, J.A., 1998. Neural responses to facial and vocal expressions of fear and disgust. Proc. R. Soc. Lond. B 265, 1809–1817. https://doi.org/doi:10.1098/rspb.1998.0506.

Phillips, M., Young, A.W., Senior, C., Brammer, M., Andrew, C., Calder, A.J., Bullmore, E.T., Perrett, sD.I., Rowland, D., Williams, S.C.R., Gray, J.A., David, A.S., 1997. A specific neural substrate for perceiving facial expressions of disgust. Nature 389, 495–498. https://doi.org/10.1038/39051.

Pujol, J., Blanco-Hinojo, L., Coronas, R., Esteba-Castillo, S., Rigla, M., Martínez-Vilavella, G., Deus, J., Novell, R., Caixàs, A., 2018. Mapping the sequence of brain events in response to disgusting food. Hum. Brain Mapp. 39, 369–380. https://doi.org/10.1002/hbm.23848.

Radua, J., Sarró, S., Vigo, T., Alonso-Lana, S., Bonnín, C.M., Ortiz-Gil, J., Canales-Rodríguez, E.J., Maristany, T., Vieta, E., McKenna, P.J., Salvador, R., Pomarol-Clotet, E., 2014. Common and specific brain responses to scenic emotional stimuli. Brain Struct. Funct. 219, 1463–1472. https://doi.org/10.1007/s00429-013-0580-0.

Reid, A.T., Hoffstaedter, F., Gong, G., Laird, A.R., Fox, P., Evans, A.C., Amunts, K., Eickhoff, S.B., 2017. A seed-based cross-modal comparison of brain connectivity measures. Brain Struct. Funct. 222, 1131–1151. https://doi.org/10.1007/s00429-016-1264-3.

Reidy, B.L., Hamann, S., Inman, C., Johnson, K.C., Brennan, P.A., 2016. Decreased sleep duration is associated with increased fMRI responses to emotional faces in children. Neuropsychologia 84, 54–62. https://doi.org/10.1016/j.neuropsychologia.2016.01.028.

Robinson, J.L., Laird, A.R., Glahn, D.C., Blangero, J., Sanghera, M.K., Pessoa, L., Fox, P.M., Uecker, A., Friehs, G., Young, K.A., Griffin, J.L., Lovallo, W.R., Fox, P.T., 2012. The functional connectivity of the human caudate: an application of meta-analytic connectivity modeling with behavioral filtering. Neuroimage 60, 117–129. https://doi.org/10.1016/j.neuroimage.2011.12.010.

Robinson, J.L., Laird, A.R., Glahn, D.C., Lovallo, W.R., Fox, P.T., 2010. Metaanalytic connectivity modeling: delineating the functional connectivity of the human amygdala. Hum. Brain Mapp. 31, 173–184. https://doi.org/10.1002/hbm.20854.

Rolls, E.T., 1999. The brain and emotion. Oxford University Press, New York.

Rolls, E.T., 2000a. The orbitofrontal cortex and reward. Cereb. Cortex 10, 284–294. https://doi.org/10.1093/cercor/10.3.284.

Rolls, E.T., 2000b. Précis of the brain and emotion. Behav. Brain Sci. 23, 177–191. https://doi.org/10.1017/S0140525X00002429.

Rolls, E.T., 2004. The functions of the orbitofrontal cortex. Brain Cogn. 55, 11–29. https://doi.org/10.1016/S0278-2626(03)00277-X.

Rozin, P., Fallon, A.E., 1987. A perspective on disgust. Psychol. Rev. 94, 23–41. https://doi.org/10.1037/0033-295X.94.1.23.

Rozin, P., Haidt, J., Fincher, K., 2009. From oral to moral. Science 323, 1179–1180. https://doi.org/10.1126/science.1170492.

Rozin, P., Haidt, J., McCauley, C., 2016. Disgust, in: Barrett, L.F., Lewis, M., Haviland-Jone, J.M. (Eds.), Handbook of Emotions, Fourth Edition. Guilford Publications, New York, pp. 815–834.

Rozin, P., Lowery, L., Ebert, R., 1994. Varieties of disgust faces and the structure of disgust. J. Pers. Soc. Psychol. 66, 870–881. https://doi.org/10.1037//0022-3514.66.5.870.

Rozzi, S., Fogassi, L., 2017. Neural coding for action execution and action observation in the prefrontal cortex and its role in the organization of socially driven behavior. Front. Neurosci. 11. https://doi.org/10.3389/fnins.2017.00492.

Sabatinelli, D., Fortune, E.E., Li, Q., Siddiqui, A., Krafft, C., Oliver, W.T., Beck, S., Jeffries, J., 2011. Emotional perception: meta-analyses of face and natural scene processing. Neuroimage 54, 2524–2533. https://doi.org/10.1016/j.neuroimage.2010.10.011.

Salih, F., Sharott, A., Khatami, R., Trottenberg, T., Schneider, G., Kupsch, A., Brown, P., Grosse, P., 2009. Functional connectivity between motor cortex and globus pallidus in human non-REM sleep. J. Physiol. 587, 1071–1086. https://doi.org/10.1113/jphysiol.2008.164327.

Sambataro, F., Dimalta, S., Di Giorgio, A., Taurisano, P., Blasi, G., Scarabino, T., Giannatempo, G., Nardini, M., Bertolino, A., 2006. Preferential responses in amygdala and insula during presentation of facial contempt and disgust. Eur. J. Neurosci. 24, 2355–2362. https://doi.org/10.1111/j.1460-9568.2006.05120.x.

Schäfer, A., Leutgeb, V., Reishofer, G., Ebner, F., Schienle, A., 2009. Propensity and sensitivity measures of fear and disgust are differentially related to emotion-specific brain activation. Neurosci. Lett. 465, 262–266. https://doi.org/10.1016/j.neulet.2009.09.030.

Schäfer, A., Schienle, A., Vaitl, D., 2005. Stimulus type and design influence hemodynamic responses towards visual disgust and fear elicitors. Int. J. Psychophysiol. 57, 53–59. https://doi.org/10.1016/j.ijpsycho.2005.01.011.

Schienle, A., Höfler, C., Keck, T., Wabnegger, A., 2020. Neural underpinnings of perception and experience of disgust in individuals with a reduced sense of smell: an fMRI study. Neuropsychologia 141, 107411. https://doi.org/10.1016/j.neuropsychologia.2020.107411.

Schienle, A., Schäfer, A., Hermann, A., Vaitl, D., 2009. Binge-eating disorder: reward sensitivity and brain activation to images of food. Biol. Psychiatry 65, 654–661. https://doi.org/10.1016/j.biopsych.2008.09.028.

Schienle, A., Schäfer, A., Hermann, A., Walter, B., Stark, R., Vaitl, D., 2006. fMRI responses to pictures of mutilation and contamination. Neurosci. Lett. 393, 174–178. https://doi.org/10.1016/j.neulet.2005.09.072.

Schienle, A., Schäfer, A., Stark, R., Walter, B., Vaitl, D., 2005a. Gender differences in the processing of disgust- and fear-inducing pictures: an fMRI study. Neuroreport 16, 277–280. https://doi.org/10.1097/00001756-200502280-00015.

Schienle, A., Schäfer, A., Stark, R., Walter, B., Vaitl, D., 2005b. Relationship between disgust sensitivity, trait anxiety and brain activity during disgust induction. Neuropsychobiology 51, 86–92. https://doi.org/10.1159/000084165.

Schienle, A., Scharmüller, W., 2013. Cerebellar activity and connectivity during the experience of disgust and happiness. Neuroscience 246, 375–381. https://doi.org/10.1016/j.neuroscience.2013.04.048.

Schienle, A., Stark, R., Walter, B., Blecker, C., Ott, U., Kirsch, P., Sammer, G., Vaitl, D., 2002. The insula is not specifically involved in disgust processing: an fMRI study. Neuroreport 13, 2023–2026. https://doi.org/10.1097/00001756-200211150-00006.

Schienle, A., Übel, S., Schöngaßner, F., Ille, R., Scharmüller, W., 2014. Disgust regulation via placebo: an fMRI study. Soc. Cogn. Affect. Neurosci. 9, 985–990. https://doi.org/10.1093/scan/nst072.

Schienle, A., Übel, S., Wabnegger, A., 2016. When opposites lead to the same: a direct comparison of explicit and implicit disgust regulation via fMRI. Soc. Cogn. Affect. Neurosci. 12, 445–451. https://doi.org/10.1093/scan/nsw144.

Schienle, A., Wabnegger, A., 2021. The processing of visual food cues during bitter aftertaste perception in females with high vs. low disgust propensity: an fMRI study. Brain Imaging Behav. https://doi.org/10.1007/s11682-021-00455-2.

Schroeder, U., Hennenlotter, A., Erhard, P., Haslinger, B., Stahl, R., Lange, K., Ceballos-Baumann, A., 2005. Functional neuroanatomy of perceiving surprised faces. Hum. Brain Mapp. 23, 181–187. https://doi.org/10.1002/hbm.20057.

Shapira, N.A., Liu, Y., He, A.G., Bradley, M.M., Lessig, M.C., James, G.A., Stein, D.J., Lang, P.J., Goodman, W.K., 2003. Brain activation by disgust-inducing pictures in obsessive-compulsive disorder. Biol. Psychiatry 54, 751–756. https://doi.org/10.1016/S0006-3223(03)00003-9.

Singer, T., Seymour, B., Doherty, J., Kaube, H., Dolan, R.J., Frith, C.D., 2004. Empathy for pain involves the affective but not sensory components of pain. Science 303, 1157–1162. https://doi.org/10.1126/science.1093535.

Smith, S.M., Fox, P.T., Miller, K.L., Glahn, D.C., Fox, P.M., Mackay, C.E., Filippini, N., Watkins, K.E., Toro, R., Laird, A.R., Beckmann, C.F., 2009. Correspondence of the brain’s functional architecture during activation and rest. Proc. Natl. Acad. Sci. U. S. A. 106, 13040–13045. https://doi.org/10.1073/pnas.0905267106.

Sprengelmeyer, R., Rausch, M., Eysel, U., Przuntek, H., 1998. Neural structures associated with recognition of facial expressions of basic emotions. Proc. R. Soc. Lond. B 265, 1927–1931. https://doi.org/10.1098/rspb.1998.0522.

Sprengelmeyer, R., Young, A.W., Calder, A.J., Karnat, A., Lange, H., Hömberg, V., Perrett, D.I., Rowland, D., 1996. Loss of disgust: perception of faces and emotions in Huntington’s disease. Brain 119, 1647–1665. https://doi.org/10.1093/brain/119.5.1647.

Stark, R., Schienle, A., Girod, C., Walter, B., Kirsch, P., Blecker, C., Ott, U., Schäfer, A., Sammer, G., Zimmermann, M., Vaitl, D., 2005. Erotic and disgust-inducing pictures - differences in the hemodynamic responses of the brain. Biol. Psychol. 70, 19–29. https://doi.org/10.1016/j.biopsycho.2004.11.014.

Stark, R., Schienle, A., Walter, B., Kirsch, P., Blecker, C., Ott, U., Schäfer, A., Sammer, G., Zimmermann, M., Vaitl, D., 2004. Hemodynamic effects of negative emotional pictures – a test-retest analysis. Neuropsychobiology 50, 108–118. https://doi.org/10.1159/000077948.

Stark, R., Schienle, A., Walter, B., Kirsch, P., Sammer, G., Ott, U., Blecker, C., Vaitl, D., 2003. Hemodynamic responses to fear and disgust-inducing pictures: an fMRI study. Int. J. Psychophysiol. 50, 225–234. https://doi.org/10.1016/S0167-8760(03)00169-7.

Stark, R., Zimmermann, M., Kagerer, S., Schienle, A., Walter, B., Weygandt, M., Vaitl, D., 2007. Hemodynamic brain correlates of disgust and fear ratings. Neuroimage 37, 663–673. https://doi.org/10.1016/j.neuroimage.2007.05.005.

Straube, T., Weisbrod, A., Schmidt, S., Raschdorf, C., Preul, C., Mentzel, H.-J., Miltner, W.H.R., 2010. No impairment of recognition and experience of disgust in a patient with a right-hemispheric lesion of the insula and basal ganglia. Neuropsychologia 48, 1735–1741. https://doi.org/10.1016/j.neuropsychologia.2010.02.022.

Surguladze, S.A., Brammer, M.J., Young, A.W., Andrew, C., Travis, M.J., Williams, S.C.R., Phillips, M.L., 2003. A preferential increase in the extrastriate response to signals of danger. Neuroimage 19, 1317–1328. https://doi.org/10.1016/S1053-8119(03)00085-5.

Surguladze, S.A., El-Hage, W., Dalgleish, T., Radua, J., Gohier, B., Phillips, M.L., 2010. Depression is associated with increased sensitivity to signals of disgust: a functional magnetic resonance imaging study. J. Psychiatr. Res. 44, 894–902. https://doi.org/10.1016/j.jpsychires.2010.02.010.

Tao, D., He, Z., Lin, Y., Liu, C., Tao, Q., 2021. Where does fear originate in the brain? A coordinate-based meta-analysis of explicit and implicit fear processing. Neuroimage 227, 117686. https://doi.org/10.1016/j.neuroimage.2020.117686.

Tettamanti, M., Rognoni, E., Cafiero, R., Costa, T., Galati, D., Perani, D., 2012. Distinct pathways of neural coupling for different basic emotions. Neuroimage 59, 1804–1817. https://doi.org/10.1016/j.neuroimage.2011.08.018.

Timmers, I., Park, A.L., Fischer, M.D., Kronman, C.A., Heathcote, L.C., Hernandez, J.M., Simons, L.E., 2018. Is empathy for pain unique in its neural correlates? A meta-analysis of neuroimaging studies of empathy. Front. Behav. Neurosci. 12. https://doi.org/10.3389/fnbeh.2018.00289.

Trautmann, S.A., Fehr, T., Herrmann, M., 2009. Emotions in motion: dynamic compared to static facial expressions of disgust and happiness reveal more widespread emotion-specific activations. Brain Res. 1284, 100–115. https://doi.org/10.1016/j.brainres.2009.05.075.

Tu, S., Qiu, J., Martens, U., Zhang, Q., 2013. Category-selective attention modulates unconscious processes in the middle occipital gyrus. Conscious. Cogn. 22, 479–485. https://doi.org/10.1016/j.concog.2013.02.007.

Tucker, D.M., Luu, P., Pribram, K.H., 1995. Social and emotional self-regulation. Ann. N. Y. Acad. Sci. 769, 213–240. https://doi.org/10.1111/j.1749-6632.1995.tb38141.x.

Turk-Browne, N., Norman-Haignere, S., McCarthy, G., 2010. Face-specific resting functional connectivity between the fusiform gyrus and posterior superior temporal sulcus. Front. Hum. Neurosci. 4, 176. https://doi.org/10.3389/fnhum.2010.00176.

Turkeltaub, P.E., Eickhoff, S.B., Laird, A.R., Fox, M., Wiener, M., Fox, P., 2012. Minimizing within-experiment and within-group effects in activation likelihood estimation meta-analyses. Hum. Brain Mapp. 33, 1–13. https://doi.org/10.1002/hbm.21186.

Uddin, L.Q., 2015. Salience processing and insular cortical function and dysfunction. Nat. Rev. Neurosci. 16, 55–61. https://doi.org/10.1038/nrn3857.

van Overveld, W.J.M., de Jong, P.J., Peters, M.L., 2009. Digestive and cardiovascular responses to core and animal-reminder disgust. Biol. Psychol. 80, 149–157. https://doi.org/10.1016/j.biopsycho.2008.08.002.

Vicario, C.M., Rafal, R.D., Martino, D., Avenanti, A., 2017. Core, social and moral disgust are bounded: a review on behavioral and neural bases of repugnance in clinical disorders. Neurosci. Biobehav. Rev. 80, 185–200. https://doi.org/10.1016/j.neubiorev.2017.05.008.

Viinikainen, M., Jääskeläinen, I.P., Alexandrov, Y., Balk, M.H., Autti, T., Sams, M., 2010. Nonlinear relationship between emotional valence and brain activity: evidence of separate negative and positive valence dimensions. Hum. Brain Mapp. 31, 1030–1040. https://doi.org/10.1002/hbm.20915.

von dem Hagen, E.A.H., Beaver, J.D., Ewbank, M.P., Keane, J., Passamonti, L., Lawrence, A.D., Calder, A.J., 2009. Leaving a bad taste in your mouth but not in my insula. Soc. Cogn. Affect. Neurosci. 4, 379–386. https://doi.org/10.1093/scan/nsp018.

Vytal, K., Hamann, S., 2010. Neuroimaging support for discrete neural correlates of basic emotions: a voxel-based meta-analysis. J. Cogn. Neurosci. 22, 2864–2885. https://doi.org/10.1162/jocn.2009.21366.

Wager, T.D., Kang, J., Johnson, T.D., Nichols, T.E., Satpute, A.B., Barrett, L.F., 2015. A Bayesian model of category-specific emotional brain responses. PLoS Comput. Biol. 11, e1004066. https://doi.org/10.1371/journal.pcbi.1004066.

Wager, T.D., Lindquist, M., Kaplan, L., 2007. Meta-analysis of functional neuroimaging data: current and future directions. Soc. Cogn. Affect. Neurosci. 2, 150–158. https://doi.org/10.1093/scan/nsm015.

Wager, T.D., Phan, K.L., Liberzon, I., Taylor, S.F., 2003. Valence, gender, and lateralization of functional brain anatomy in emotion: a meta-analysis of findings from neuroimaging. Neuroimage 19, 513–531. https://doi.org/10.1016/S1053-8119(03)00078-8.

Wang, Y., Braver, T.S., Yin, S., Hu, X., Wang, X., Chen, A., 2019. Reward improves response inhibition by enhancing attentional capture. Soc. Cogn. Affect. Neurosci. 14, 35–45. https://doi.org/10.1093/scan/nsy111.

Weiner, K.S., Zilles, K., 2016. The anatomical and functional specialization of the fusiform gyrus. Neuropsychologia 83, 48–62. https://doi.org/10.1016/j.neuropsychologia.2015.06.033.

Wertheim, J., Ragni, M., 2018. The neural correlates of relational reasoning: a meta-analysis of 47 functional magnetic resonance studies. J. Cogn. Neurosci. 30, 1734–1748. https://doi.org/10.1162/jocn_a_01311.

Wicker, B., Keysers, C., Plailly, J., Royet, J.-P., Gallese, V., Rizzolatti, G., 2003. Both of us disgusted in my insula: the common neural basis of seeing and feeling disgust. Neuron 40, 655–664. https://doi.org/10.1016/S0896-6273(03)00679-2.

Winston, J.S., O’Doherty, J., Dolan, R.J., 2003. Common and distinct neural responses during direct and incidental processing of multiple facial emotions. Neuroimage 20, 84–97. https://doi.org/10.1016/S1053-8119(03)00303-3.

Woo, C.-W., Chang, L.J., Lindquist, M.A., Wager, T.D., 2017. Building better biomarkers: brain models in translational neuroimaging. Nat. Neurosci. 20, 365–377. https://doi.org/10.1038/nn.4478.

Wright, P., He, G., Shapira, N., Goodman, W., Liu, Y., 2004. Disgust and the insula: fMRI responses to pictures of mutilation and contamination. Neuroreport 15, 2347–2351. https://doi.org/10.1097/00001756-200410250-00009.

Yamasaki, H., LaBar, K.S., McCarthy, G., 2002. Dissociable prefrontal brain systems for attention and emotion. Proc. Natl. Acad. Sci. U. S. A. 99, 11447–11451. https://doi.org/10.1073/pnas.182176499.

Yao, S., Becker, B., Zhao, W., Zhao, Z., Kou, J., Ma, X., Geng, Y., Ren, P., Kendrick, K.M., 2018a. Oxytocin modulates attention switching between interoceptive signals and external social cues. Neuropsychopharmacology 43, 294–301. https://doi.org/10.1038/npp.2017.189.

Yao, S., Zhao, W., Geng, Y., Chen, Y., Zhao, Z., Ma, X., Xu, L., Becker, B., Kendrick, K.M., 2018b. Oxytocin facilitates approach behavior to positive social stimuli via decreasing anterior insula activity. Int. J. Neuropsychopharmacol. 21, 918–925. https://doi.org/10.1093/ijnp/pyy068.

Yarkoni, T., Poldrack, R.A., Nichols, T.E., Van Essen, D.C., Wager, T.D., 2011. Large-scale automated synthesis of human functional neuroimaging data. Nat. Meth. 8, 665–670. https://doi.org/10.1038/nmeth.1635.

Yu, F., Barron, D.S., Tantiwongkosi, B., Fox, M., Fox, P., 2018. Characterisation of meta-analytical functional connectivity in progressive supranuclear palsy. Clin. Radiol. 73, 415.e411-415.e417. https://doi.org/10.1016/j.crad.2017.11.007.

Zald, D.H., 2003. The human amygdala and the emotional evaluation of sensory stimuli. Brain Res. Rev. 41, 88–123. https://doi.org/10.1016/S0165-0173(02)00248-5.

Zhou, F., Li, J., Zhao, W., Xu, L., Zheng, X., Fu, M., Yao, S., Kendrick, K.M., Wager, T.D., Becker, B., 2020. Empathic pain evoked by sensory and emotional-communicative cues share common and process-specific neural representations. eLife 9, e56929. https://doi.org/10.7554/eLife.56929.

Zhou, F., Zhao, W., Qi, Z., Geng, Y., Yao, S., Kendrick, K.M., Wager, T.D., Becker, B., 2021. A distributed fMRI-based signature for the subjective experience of fear. Nat. Commun. 12, 6643. https://doi.org/10.1038/s41467-021-26977-3.

Ziegler, J., Montant, M., Briesemeister, B., Brink, T., Wicker, B., Ponz, A., Bonnard, M., Jacobs, A., Braun, M., 2018. Do words stink? Neural reuse as a principle for understanding emotions in reading. J. Cogn. Neurosci. 30, 1023–1032. https://doi.org/10.1162/jocn_a_01268.

Zilbovicius, M., Meresse, I., Chabane, N., Brunelle, F., Samson, Y., Boddaert, N., 2006. Autism, the superior temporal sulcus and social perception. Trends Neurosci. 29, 359–366. https://doi.org/10.1016/j.tins.2006.06.004.

Zimmer, U., Höfler, M., Koschutnig, K., Ischebeck, A., 2016. Neuronal interactions in areas of spatial attention reflect avoidance of disgust, but orienting to danger. Neuroimage 134, 94–104. https://doi.org/10.1016/j.neuroimage.2016.03.050.

