## Supplementary Materials for "Common and distinct neurofunctional representations of core and social disgust in the brain: Coordinate-based and network meta-analyses"

1. Contrast and conjunction analyses

Contrast analyses compare and contrast two ALE datasets. When performing contrast analyses, GingerALE randomly divides the pooled foci datasets into two new datasets of the same size as the original individual computed datasets. Next, voxel-wise differences between these two new datasets are determined through a subtracting procedure: an ALE image is created for each new dataset, then subtracted from the other, and finally compared with the true data. After many permutations, this generates a voxel-wise p value image showing where the true data's values sit on the distribution of values in that voxel. Different from the statistical procedure used for contrast analyses, conjunction is conducted using voxel-wise minimum value of the input ALE images to create the output image. The expected results represent the shared neural substrates in core disgust and social disgust processing (Eickhoff et al., 2011).

2. Percentage of overlap

2.1 For general disgust processing

| Cluster | Percentage of overlap |
|---|---|
| 1 | 64.3% Amygdala, 18.1% Brodmann area 34, 13.7% Globus Pallidus, 3.8% Brodmann area 28 |
| 2 | 53.7% Insula, 37.9% Inferior Frontal Gyrus, 4.2% Claustrum, 4.2% Extra-Nuclear |
| 3 | 95.5% Fusiform Gyrus, 1.5% Inferior Temporal Gyrus, 1.5% Declive, 1.5% Parahippocampal Gyrus |
| 4 | 51.6% Fusiform Gyrus, 41.9% Declive, 6.5% Culmen |
| 5 | 64.9% Amygdala, 31.1% Brodmann area 34, 4.1% Brodmann area 28 |
| 6 | 60.4% Middle Occipital Gyrus, 18.9% Inferior Occipital Gyrus, 15.3% Fusiform Gyrus, 5.4% Declive |
| 7 | 44.4% Inferior Occipital Gyrus, 33.3% Lingual Gyrus, 22.2% Fusiform Gyrus |
| 8 | 80.8% Middle Occipital Gyrus, 7.7% Inferior Temporal Gyrus, 7.7% Middle Temporal Gyrus, 3.8% Inferior Occipital Gyrus |
| 9 | 83.5% Insula, 16.5% Inferior Frontal Gyrus |
| 10 | 75% Inferior Parietal Lobule, 25% Postcentral Gyrus |
| 11 | 73.1% Inferior Frontal Gyrus, 26.9% Precentral Gyrus |
| 12 | 95.8% Superior Frontal Gyrus, 4.2% Medial Frontal Gyrus |

2.2 For core disgust processing

| Cluster | Percentage of overlap |
|---|---|
| 1 | 62.4% Amygdala, 32.5% Brodmann area 34, 3.8% Brodmann area 28 |
| 2 | 68.1% Declive, 31.9% Fusiform Gyrus |
| 3 | 70.6% Amygdala, 18.3% Brodmann area 34, 4.6% Brodmann area 28, 6.5 % Globus Pallidus |
| 4 | 69% Inferior Parietal Lobule, 31% Postcentral Gyrus |
| 5 | 60.6% Middle Occipital Gyrus, 24.2% Inferior Temporal Gyrus, 9.1% Middle Temporal Gyrus, 6.1% Inferior Occipital Gyrus |
| 6 | 81.3% Superior Frontal Gyrus, 18.8% Medial Frontal Gyrus |
| 7 | 55.6% Lingual Gyrus, 44.4% Inferior Occipital Gyrus |

| | |
|---|---|
| 8 | 53.8% Insula, 46.2% Inferior Frontal Gyrus |
| 9 | 54% Insula, 38% Inferior Frontal Gyrus, 8% Extra-Nuclear |
| 10 | 70.2% Lingual Gyrus, 29.8% Inferior Occipital Gyrus |

2.3 For social disgust processing

| Cluster | Percentage of overlap |
|---|---|
| 1 | 54.2% Insula, 40.7% Inferior Frontal Gyrus, 5.1% Precentral Gyrus |
| 2 | 58.4% Medial Frontal Gyrus, 31.5% Superior Frontal Gyrus, 10.1% Cingulate Gyrus |
| 3 | 100% Middle Occipital Gyrus |
| 4 | 83.1% Declive, 16.9% Fusiform Gyrus |
| 5 | 88.1% Fusiform Gyrus, 9.5% Culmen, 2.4% Inferior Temporal Gyrus |
| 6 | 55.6% Middle Temporal Gyrus, 44.4% Superior Temporal Gyrus |

References

Eickhoff, S.B., Bzdok, D., Laird, A.R., Roski, C., Caspers, S., Zilles, K., Fox, P.T., 2011. Co-activation patterns distinguish cortical modules, their connectivity and functional differentiation. Neuroimage 57, 938-949. https://doi.org/10.1016/j.neuroimage.2011.05.021.
