## Supplementary Table 1 for "Common and distinct neurofunctional representations of core and social disgust in the brain: Coordinate-based and network meta-analyses"

Supplementary Table 1. Characteristics of the included studies with reference.

| Authors | Magnetic field, T | Gender Ratio (f/m) | Age | Stimuli type | Contrasts | Coordinates | Analyze software |
|---|---|---|---|---|---|---|---|
| Ahn et al., 2014 | 3 | 42/41 | M=29.0; SD=11.3; (18–62) | core disgust affective picture (IAPS) | Disgust > Neutral | MNI | SPM8 |
| Ashworth et al., 2011 | 1.5 | 16/0 | M=27.4; SD=5.4 | facial expressions (Matsumoto & Ekman) | Disgust > Baseline | MNI | FSL v5.98 |
| Azuma et al., 2015 | 1.5 | 5/9 | M=13; SD=3; (8–17) | facial expressions (FEEST) | 100% Disgust > Fixation | TAL | XBAM v4 |
| Baumann et al., 2012 | 3 | 15/15 | M=22.2; SD=2.9; (18-30) | core disgust affective picture (IAPS) | Disgust > Neutral | MNI | SPM5 |
| Benuzzi et al., 2008 | 3 | 15/0 | M=23.5; (19–31) | core disgust affective film clips | Disgust > Neutral | MNI | SPM2 |
| Burklund et al., 2007 | 3 | 13/6 | Mean age about 27 | facial expressions film clips (self-validated) | Disgust > Fixation | MNI | SPM9 |
| Calder et al., 2007 | 3 | 7/5 | M=22; SD=2.4 | core disgust affective picture (self-validated) | Disgusting > Non-foods<br>Disgusting > Bland<br>Disgusting > Appetizing | MNI | SPM99 |
| Chakrabarti et al., 2006 | 3 | 13/13 | M=23.4; SD=4.23 | facial expressions film clips (self-validated) | Disgust > Neutral | TAL | SPM2 |
| Chen et al., 2021 | 3 | 15/16 | M=21.34 | core disgust affective picture (IAPS+CAPS) | Watching-disgust > Watching-neutral | MNI | SPM8 |
| de Greck et al., 2012 | 1.5 | 12/8 | M=37; SD=10.6 | facial expressions (JACFEE & JACNeuF) | Disgust > Control | TAL | AFNI |
| Deeley et al., 2008 | 1.5 | 0/40 | M=24; SD=9.6; (8–50) | facial expressions (FEEST) | Disgust > Neutral | TAL | Other |
| Deeley et al., 2007 | 1.5 | 0/9 | M=27; SD=5 | facial expressions (FEEST) | Disgust > Fixation | TAL | Other |
| Fitzgerald et al., 2006 | 4 | 10/10 | M=26; (19–42) | facial expressions (the standardized Penn Emotion Recognition set) | Disgust > Baseline | MNI | SPM2 |
| Goldin et al. 2008 | 3 | 17/0 | M=22.7; SD=3.5 | core disgust affective film clips (self-validated) | Watch-disgust > Watch-neutral | TAL | AFNI |
| Harris et al., 2006 | 3 | 10mix | | core disgust affective picture | Disgust > Fixation | TAL | Brain Voyager |
| Harrison et al., 2010 | 3 | 7/5 | M=25.9; SD=5.7 | core disgust affective film clips (self-validated) | Disgust > Control | MNI | SPM5 |

| | | | | | | | |
|---|---|---|---|---|---|---|---|
| Hennenlotter et al., 2004 | 1.5 | 4/5 | M age about 37 | facial expressions (FEEST) | Disgust > Neutral | TAL | SPM99 |
| Hermann et al., 2007 | 1.5 | 10/0 | M=27.6; SD=10.7 | core disgust affective picture (IAPS + self-validated) | Disgust > Neutral | MNI | SPM2 |
| Jabbi et al., 2008 | 3 | 6/6 | | facial expressions film clips | Disgust > Neutral | MNI | SPM2 |
| Jehna et al., 2011a | 3 | 10/5 | M=30.3; SD=10.6 | facial expressions (KDEF) | Disgust > Neutral | MNI | FEAT (part of FSL) v5.63 |
| Jehna et al., 2011b | 3 | 21/9 | M=36.1; SD=14.1; (17–66) | facial expressions (the Karolinska Directed Emotional Faces set) | Disgust > Neutral | MNI | FEAT (part of FSL) v5.98 |
| Karama et al., 2011 | 1.5 | 0/18 | M=about 25.5; SD=about 3.4;(about 21-30) | core disgust affective film clips (self-validated) | Disgust > Neutral | MNI | SPM5 |
| Lassalle et al., 2019 | 3 | 3/17 | M=24.15; SD=7.57 | core disgust affective film clips | Disgust > Neutral | MNI | FEAT (part of FSL) v6 |
| Malhi et al., 2007 | 3 | 10/0 | M=32.4; SD=6.4; (24–45) | facial expressions (Ekman and Friesen) | Disgust > Neutral | TAL | Brain Voyager |
| Phillips et al., 2000 | 1.5 | 7/7 | M=31; (20–48) | core disgust affective picture (IAPS) | Disgust > Neutral | TAL | Other |
| Phillips et al., 2001 | 1.5 | 2/4 | M=33.8; (24-48) | core disgust affective picture (IAPS) | Aversive > Neutral | TAL | Other |
| Phillips et al., 2004 | 1.5 | 0/8 | M=31.9; (25–36) | facial expressions (Ekman and Friesen) | Disgust > Neutral | TAL | Other |
| Phillips et al., 1999 | 1.5 | 5mix | M=30; (22-43) | facial expressions (Ekman and Friesen) | Disgust > Neutral | TAL | Other |
| Phillips et al., 1998 | 1.5 | 0/6 | M=37; (25-43) | facial expressions (Ekman & Friesen) | Disgust > Neutral | TAL | ANMR |
| Phillips et al., 1997 | 1.5 | 5/2 | M=27 | facial expressions (Ekman & Friesen) | 150% Disgust > Neutral | TAL | Other |
| Pitskel et al., 2011 | 3 | 6/9 | M=13.03; SD=2.20; (7–17) | core disgust affective picture (IAPS) | Look-disgust > Look-neutral | TAL | Brain Voyager |
| Pujol et al., 2018 | 1.5 | 15/15 | M=27.9; SD=7.8; (19-45) | core disgust affective film clips (self-validated) | Disgusting > Appetizing | MNI | SPM8 |
| Radua et al., 2014 | 1.5 | 28/12 | M=38; SD=11; (19–59) | core disgust affective picture (IAPS) | Disgust > Neutral | MNI | FEAT (part of FSL) |

| | | | | | | | |
|---|---|---|---|---|---|---|---|
| Reidy et al., 2016 | 3 | 0/15 | M=8.67; SD=1.05; (7-11) | facial expressions (NimStim set) | Disgust > Neutral | MNI | SPM8 |
| Rymarczyk et al., 2019 | 3 | 21/25 | M=23.8; SD=2.5 | facial expressions film clips (The Amsterdam Dynamic Facial Expression Set) | Disgust dynamic > Neutral dynamic<br><br>Disgust static > Neutral static | MNI | SPM12 |
| Salloum et al., 2007 | 1.5 | 0/11 | M=36; SD=5.9; (25-45) | facial expressions (EFE, Matsumoto and Ekman) | 70% Disgust > Baseline | TAL | AFNI |
| Sambataro et al., 2006 | 3 | 13/11 | M=26.8; SD=5.6 | facial expressions (JACFEE) | Disgust > Neutral | TAL | SPM99 |
| Schäfer et al., 2009 | 3 | 18/0 | M=24.8; SD=2.4 | core disgust affective picture (IAPS + self-validated) | Disgust > Neutral | MNI | SPM2 |
| Schäfer et al., 2005 | 1.5 | 10/10 | M=23.93; (19–32) | core disgust affective picture (IAPS + self-validated) | Disgust > Neutral (block design)<br>Disgust > Neutral (event-related design) | MNI | SPM2 |
| Schienle et al., 2020 | 3 | 5/21 | M=32.65; SD=13.47 | core disgust affective picture | Disgust > Neutral | MNI | SPM12 |
| Schienle et al., 2015 | 3 | 11/11 | M=51.8; SD=9.8 | core disgust affective picture (self-validated) | Disgust > Neutral | MNI | SPM12 |
| Schienle et al., 2009 | 1.5 | 19/0 | M=22.3; SD=2.6 | core disgust affective picture | Disgust > Neutral healthy control-normal weight | MNI | SPM2 |
| | | 17/0 | M=25.0; SD=4.7 | core disgust affective picture | Disgust > Neutral healthy control-outweight | | |
| Schienle et al., 2006 | 1.5 | 12/0 | (19-41) | core disgust affective picture (IAPS + self-validated) | Contamination > Neutral<br><br>Mutilation > Neutral | MNI | SPM2 |
| Schienle et al., 2005a | 1.5 | 63/0 | M=27.3; SD=8.4 | core disgust affective picture (IAPS + self-validated) | Disgust > Neutral | MNI | SPM99 |
| Schienle et al., 2005b | 1.5 | 13/0 | M=23.9; SD=6.8 | core disgust affective picture (IAPS + self-validated) | Disgust > Neutral | MNI | SPM99 |
| Schienle et al., 2013 | 1.5 | 34/0 | M=23; SD=3.4 | core disgust affective picture (IAPS + self-validated) | Disgust > Neutral | MNI | SPM8 |
| Schienle et al., 2004 | 1.5 | 12/0 | M=26.3; SD=6.4 | core disgust affective picture (IAPS + self-validated) | Disgust > Neutral | MNI | SPM99 |

| | | | | | | | |
|---|---|---|---|---|---|---|---|
| Schienle et al., 2002 | 1.5 | 12/0 | M=26.3; (21–41) | core disgust affective picture (IAPS + self-validated) | Disgust > Neutral | MNI | SPM99 |
| Schienle et al., 2014 | 3 | 34/0 | M=23.9; SD=4 | core disgust affective picture (IAPS + self-validated) | Disgust > Neutral | MNI | SPM8 |
| Schroeder et al., 2005 | 1.5 | 10/10 | M=32.5; SD=8.3 | facial expressions (FEEST) | Disgust > Neutral | TAL | SPM99 |
| Shapira et al., 2003 | 3 | 5/3 | M=38; (34-44) | core disgust affective picture (IAPS) | Disgust > Neutral | TAL | MEDx |
| Shimamura et al., 2013 | 4 | 13/7 | M=21.1; (18–33) | core disgust affective film clips (self-validated) | Disgust express > Neutral express | MNI | SPM2 |
| Sprengelmeyer et al., 1998 | 2 | 4/2 | M=23.5; SD=1.3 | facial expressions (Ekman & Friesen) | Disgust > Neutral | TAL | SPM96 |
| Stark et al., 2005a | 1.5 | 6/6 | M about 28.2; (20–40) | core disgust affective picture (IAPS + self-validated) | Disgust > Neutral (nonSM) | MNI | SPM99 |
| | | 6/6 | M about 28.2; (20–40) | core disgust affective picture (IAPS + self-validated) | Disgust > Neutral (SM) | | |
| Stark et al., 2005b | 1.5 | 11/4 | M=29.1; (20–41) | core disgust affective film clips (self-validated); | Disgust > Neutral | MNI | SPM2 |
| Stark et al., 2004 | 1.5 | 0/24 | M=25.5; SD=2.65; (20–31) | core disgust affective picture (IAPS + self-validated) | Disgust > Neutral | MNI | SPM99 |
| Stark et al., 2007 | 1.5 | 32/34 | M=24.7; SD=5.2; (19-44) | core disgust affective picture (IAPS + self-validated) | Disgust > Neutral | MNI | SPM2 |
| Surguladze et al., 2003 | 1.5 | 4/5 | M=39.6; (23–63) | facial expressions (FEEST) | Disgust > Neutral | TAL | Other |
| Surguladze et al., 2010 | 1.5 | 4/5 | M=39.7; SD=14.6 | facial expressions (FEEST) | Disgust (100%) > Neutral | TAL | XBAM |
| Tettamanti et al., 2012 | 3 | 19/0 | M=24.1; SD=5.2; (19–39) | core disgust affective film clips | Disgust > Neutral | MNI | SPM5 |
| Trautmann et al., 2009 | 3 | 16/0 | M=21.6; SD=2.3; (19-27) | facial expressions (self-validated) | dynamic_disgust > dynamic_neutral<br><br>static_disgust > static_neutral | TAL | SPM2 |
| Viol et al., 2019 | 3 | 17mix | M age about 43.5 | core disgust affective picture (IAPS) | Disgust > Neutral | MNI | SPM12 |
| von dem Hagen et al., 2009 | 3 | 14/13 | M=27; SD=8 | facial expressions (NimStim face stimulus | Canonical disgust > Neutral | MNI | SPM2 |

| | | | | | | | |
|---|---|---|---|---|---|---|---|
| | | | | set + self-validated) | | | |
| Wabnegger et al., 2018 | 3 | 16/0 | M=31.13; SD=12.18 | core disgust affective picture (IAPS + self-validated) | Disgust > Neutral | MNI | SPM12 |
| Wicker et al., 2003 | 3 | 0/14 | (20–27) | facial expressions film clips | Disgust > Neutral | MNI | SPM99 |
| Williams et al., 2005 | 1.5 | 8/5 | M=24; SD=8 | facial expressions (Ekman & Friesen) | Disgust > Neutral | TAL | Other |
| Wittfoth et al., 2020 | | 8/9 | M=23.47; SD=2.45 | core disgust affective picture (IAPS) | Disgust > Neutral | MNI | SPM12 |
| Wright et al., 2004 | 3 | 4/4 | (20–26) | core disgust affective picture (IAPS) | Mutilation > Neutral | TAL | Brain Voyager v4.9.6 |
| | | | | | Contamination > Neutral | | |
| Ziegler et al., 2018 | 3 | 23/13 | M=24.5; SD=4.0; (19-35) | facial expressions film clips | Disgust > Neutral | MNI | SPM8 |

**Reference of studies included for meta-analysis**


Ahn, W.-Y., Kishida, K.T., Gu, X., Lohrenz, T., Harvey, A., Alford, J.R., Smith, K.B., Yaffe, G., Hibbing, J.R., Dayan, P., Montague, P.R., 2014. Nonpolitical images evoke neural predictors of political ideology. Curr. Biol. 24, 2693-2699. https://doi.org/10.1016/j.cub.2014.09.050.

Ashworth, F., Pringle, A., Norbury, R., Harmer, C.J., Cowen, P.J., Cooper, M.J., 2011. Neural response to angry and disgusted facial expressions in bulimia nervosa. Psychol. Med. 41, 2375-2384. https://doi.org/10.1017/S0033291711000626.

Azuma, R., Deeley, Q., Campbell, L.E., Daly, E.M., Giampietro, V., Brammer, M.J., Murphy, K.C., Murphy, D.G., 2015. An fMRI study of facial emotion processing in children and adolescents with 22q11.2 deletion syndrome. J. Neurodev. Disord. 7, 1. https://doi.org/10.1186/1866-1955-7-1.

Baumann, O., Mattingley, J.B., 2012. Functional topography of primary emotion processing in the human cerebellum. Neuroimage 61, 805-811. https://doi.org/10.1016/j.neuroimage.2012.03.044.

Benuzzi, F., Lui, F., Duzzi, D., Nichelli, P., Porro, C., 2008. Does it look painful or disgusting? Ask your parietal and cingulate cortex. J. Neurosci. 28, 923-931. https://doi.org/10.1523/JNEUROSCI.4012-07.2008.

Burklund, L., Eisenberger, N., Lieberman, M., 2007. The face of rejection: rejection sensitivity moderates dorsal anterior cingulate activity to disapproving facial expressions. Soc. Neurosci. 2, 238-253. https://doi.org/10.1080/17470910701391711.

Calder, A.J., Beaver, J.D., Davis, M.H., Van Ditzhuijzen, J., Keane, J., Lawrence, A.D., 2007. Disgust sensitivity predicts the insula and pallidal response to pictures of disgusting foods. Eur. J. Neurosci. 25, 3422-3428. https://doi.org/10.1111/j.1460-9568.2007.05604.x.

Chakrabarti, B., Bullmore, E., Baron-Cohen, S., 2006. Empathizing with basic emotions: common and discrete neural substrates. Soc. Neurosci. 1, 364-384. https://doi.org/10.1080/17470910601041317.

Chen, S., Ding, N., Wang, F., Li, Z., Qin, S., Biswal, B.B., Yuan, J., 2021. Functional decoupling of emotion coping network subsides automatic emotion regulation by implementation intention. Neural Plast. 2021, 6639739. https://doi.org/10.1155/2021/6639739.

de Greck, M., Scheidt, L., Bölter, A.F., Frommer, J., Ulrich, C., Stockum, E., Enzi, B., Tempelmann, C., Hoffmann, T., Han, S., Northoff, G., 2012. Altered brain activity during emotional empathy in somatoform disorder. Hum. Brain Mapp. 33, 2666-2685. https://doi.org/10.1002/hbm.21392.

Deeley, Q., Daly, E.M., Azuma, R., Surguladze, S., Giampietro, V., Brammer, M.J., Hallahan, B., Dunbar, R.I.M., Phillips, M.L., Murphy, D.G.M., 2008. Changes in male brain responses to emotional faces from adolescence to middle age. Neuroimage 40, 389-397. https://doi.org/10.1016/j.neuroimage.2007.11.023.

Deeley, Q., Daly, E.M., Surguladze, S., Page, L., Toal, F., Robertson, D., Curran, S., Giampietro, V., Seal, M., Brammer, M.J., Andrew, C., Murphy, K., Phillips, M.L., Murphy, D.G.M., 2007. An event related functional magnetic resonance imaging study of facial emotion processing in asperger syndrome. Biol. Psychiatry 62, 207-217. https://doi.org/10.1016/j.biopsych.2006.09.037.

Fitzgerald, D.A., Angstadt, M., Jelsone, L.M., Nathan, P.J., Phan, K.L., 2006. Beyond threat: amygdala reactivity across multiple expressions of facial affect. Neuroimage 30, 1441-1448. https://doi.org/10.1016/j.neuroimage.2005.11.003.

Goldin, P.R., McRae, K., Ramel, W., Gross, J.J., 2008. The neural bases of emotion regulation: reappraisal and suppression of negative emotion. Biol. Psychiatry 63, 577-586. https://doi.org/10.1016/j.biopsych.2007.05.031.

Harris, L.T., Fiske, S.T., 2006. Dehumanizing the lowest of the low: neuroimaging responses to extreme out-groups. Psychol. Sci. 17, 847-853. https://doi.org/10.1111/j.1467-9280.2006.01793.x.

Harrison, N.A., Gray, M.A., Gianaros, P.J., Critchley, H.D., 2010. The embodiment of emotional feelings in the brain. J. Neurosci. 30, 12878-12884. https://doi.org/10.1523/JNEUROSCI.1725-10.2010.

Hennenlotter, A., Schroeder, U., Erhard, P., Haslinger, B., Stahl, R., Weindl, A., Einsiedel, H.G., Lange, K., Ceballos-Baumann, A., 2004. Neural correlates associated with impaired disgust processing in pre-symptomatic Huntington's disease. Brain 127, 1446-1453. https://doi.org/10.1093/brain/awh165.

Hermann, A., Schäfer, A., Walter, B., Stark, R., Vaitl, D., Schienle, A., 2007. Diminished medial prefrontal cortex activity in blood-injection-injury phobia. Biol. Psychol. 75, 124-130. https://doi.org/10.1016/j.biopsycho.2007.01.002.

Jabbi, M., Bastiaansen, J., Keysers, C., 2008. A common anterior insula representation of disgust observation, experience and imagination shows divergent functional connectivity pathways. PLoS ONE 3, e2939. https://doi.org/10.1371/journal.pone.0002939.

Jehna, M., Langkammer, C., Wallner-Blazek, M., Neuper, C., Loitfelder, M., Ropele, S., Fuchs, S., Khalil, M., Pluta-Fuerst, A., Fazekas, F., Enzinger, C., 2011a. Cognitively preserved MS patients demonstrate functional differences in processing neutral and emotional faces. Brain Imaging Behav. 5, 241-251. https://doi.org/10.1007/s11682-011-9128-1.

Jehna, M., Neuper, C., Ischebeck, A., Loitfelder, M., Ropele, S., Langkammer, C., Ebner, F., Fuchs, S., Schmidt, R., Fazekas, F., Enzinger, C., 2011b. The functional correlates of face perception and recognition of emotional facial expressions as evidenced by fMRI. Brain Res. 1393, 73-83. https://doi.org/10.1016/j.brainres.2011.04.007.

Karama, S., Armony, J., Beauregard, M., 2011. Film excerpts shown to specifically elicit various affects lead to overlapping activation foci in a large set of symmetrical brain regions in males. PLoS ONE 6, e22343. https://doi.org/10.1371/journal.pone.0022343.

Lassalle, A., Zürcher, N.R., Porro, C.A., Benuzzi, F., Hippolyte, L., Lemonnier, E., Åsberg Johnels, J., Hadjikhani, N., 2019. Influence of anxiety and alexithymia on brain activations associated with the perception of others' pain in autism. Soc. Neurosci. 14, 359-377. https://doi.org/10.1080/17470919.2018.1468358.

Malhi, G., Lagopoulos, J., Sachdev, P., Ivanovski, B., Shnier, R., Ketter, T., 2007. Is a lack of disgust something to fear? An fMRI facial recognition study in euthymic bipolar disorder patients. Bipolar Disord. 9, 345-357. https://doi.org/10.1111/j.1399-5618.2007.00485.x.

Phillips, M., Marks, I., Senior, C., Lythgoe, D., O'Dwyer, A., Meehan, O., Williams, S., Brammer, M., Bullmore, E., McGuire, P., 2000. A differential neural response in obsessive-compulsive disorder patients with washing compared with checking symptoms to disgust. Psychol. Med. 30, 1037-1050. https://doi.org/10.1017/S0033291799002652.

Phillips, M., Medford, N., Senior, C., Bullmore, E.T., Suckling, J., Brammer, M.J., Andrew, C., Sierra, M., Williams, S.C.R., David, A.S., 2001. Depersonalization disorder: thinking without feeling. Psychiatry Res Neuroimaging 108, 145-160. https://doi.org/10.1016/S0925-4927(01)00119-6.

Phillips, M., Williams, L., Heining, M., Herba, C., Russell, T., Andrew, C., Bullmore, E., Brammer, M., Williams, S., Morgan, M., Young, A., Gray, J., 2004. Differential neural responses to overt and covert presentations of facial expressions of fear and disgust. Neuroimage 21, 1484-1496. https://doi.org/10.1016/j.neuroimage.2003.12.013.

Phillips, M., Williams, L., Senior, C., Bullmore, E.T., Brammer, M.J., Andrew, C., Williams, S.C.R., David, A.S., 1999. A differential neural response to threatening and non-threatening negative facial expressions in paranoid and non-paranoid schizophrenics. Psychiatry Res Neuroimaging 92, 11-31. https://doi.org/10.1016/S0925-4927(99)00031-1.

Phillips, M., Young, A.W., Scott, S.K., Calder, A.J., Andrew, C., Giampietro, V., Williams, S.C.R., Bullmore, E.T., Brammer, M., Gray, J.A., 1998. Neural responses to facial and vocal expressions of fear and disgust. Proc. R. Soc. Lond. B 265, 1809-1817. https://doi.org/doi:10.1098/rspb.1998.0506.

Phillips, M., Young, A.W., Senior, C., Brammer, M., Andrew, C., Calder, A.J., Bullmore, E.T., Perrett, D.I., Rowland, D., Williams, S.C.R., Gray, J.A., David, A.S., 1997. A specific neural substrate for perceiving facial expressions of disgust. Nature 389, 495-498. https://doi.org/10.1038/39051.

Pitskel, N.B., Bolling, D.Z., Kaiser, M.D., Crowley, M.J., Pelphrey, K.A., 2011. How grossed out are you? The neural bases of emotion regulation from childhood to adolescence. Dev. Cogn. Neurosci. 1, 324-337. https://doi.org/10.1016/j.dcn.2011.03.004.

Pujol, J., Blanco-Hinojo, L., Coronas, R., Esteba-Castillo, S., Rigla, M., Martínez-Vilavella, G., Deus, J., Novell, R., Caixàs, A., 2018. Mapping the sequence of brain events in response to disgusting food. Hum. Brain Mapp. 39, 369-380. https://doi.org/10.1002/hbm.23848.

Radua, J., Sarró, S., Vigo, T., Alonso-Lana, S., Bonnín, C.M., Ortiz-Gil, J., Canales-Rodríguez, E.J., Maristany, T., Vieta, E., McKenna, P.J., Salvador, R., Pomarol-Clotet, E., 2014. Common and specific brain responses to scenic emotional stimuli. Brain Struct. Funct. 219, 1463-1472. https://doi.org/10.1007/s00429-013-0580-0.

Reidy, B.L., Hamann, S., Inman, C., Johnson, K.C., Brennan, P.A., 2016. Decreased sleep duration is associated with increased fMRI responses to emotional faces in children. Neuropsychologia 84, 54-62. https://doi.org/10.1016/j.neuropsychologia.2016.01.028.

Rymarczyk, K., Żurawski, Ł., Jankowiak-Siuda, K., Szatkowska, I., 2019. Empathy in facial mimicry of fear and disgust: simultaneous EMG-fMRI recordings during observation of static and dynamic facial expressions. Front. Psychol. 10, 701. https://doi.org/10.3389/fpsyg.2019.00701.

Salloum, J.B., Ramchandani, V.A., Bodurka, J., Rawlings, R., Momenan, R., George, D., Hommer, D.W., 2007. Blunted rostral anterior cingulate response during a simplified decoding task of negative emotional facial expressions in alcoholic patients. Alcohol. Clin. Exp. Res. 31, 1490-1504. https://doi.org/10.1111/j.1530-0277.2007.00447.x.

Sambataro, F., Dimalta, S., Di Giorgio, A., Taurisano, P., Blasi, G., Scarabino, T., Giannatempo, G., Nardini, M., Bertolino, A., 2006. Preferential responses in amygdala and insula during presentation of facial contempt and disgust. Eur. J. Neurosci. 24, 2355-2362. https://doi.org/10.1111/j.1460-9568.2006.05120.x.

Schäfer, A., Leutgeb, V., Reishofer, G., Ebner, F., Schienle, A., 2009. Propensity and sensitivity measures of fear and disgust are differentially related to emotion-specific brain activation. Neurosci. Lett. 465, 262-266. https://doi.org/10.1016/j.neulet.2009.09.030.

Schäfer, A., Schienle, A., Vaitl, D., 2005. Stimulus type and design influence hemodynamic responses towards visual disgust and fear elicitors. Int. J. Psychophysiol. 57, 53-59. https://doi.org/10.1016/j.ijpsycho.2005.01.011.

Schienle, A., Höfler, C., Keck, T., Wabnegger, A., 2020. Neural underpinnings of perception and experience of disgust in individuals with a reduced sense of smell: an fMRI study. Neuropsychologia 141, 107411. https://doi.org/10.1016/j.neuropsychologia.2020.107411.

Schienle, A., Ille, R., Wabnegger, A., 2015. Experience of negative emotions in Parkinson's disease: an fMRI investigation. Neurosci. Lett. 609, 142-146. https://doi.org/10.1016/j.neulet.2015.10.046.

Schienle, A., Schäfer, A., Hermann, A., Vaitl, D., 2009. Binge-eating disorder: reward sensitivity and brain activation to images of food. Biol. Psychiatry 65, 654-661. https://doi.org/10.1016/j.biopsych.2008.09.028.

Schienle, A., Schäfer, A., Hermann, A., Walter, B., Stark, R., Vaitl, D., 2006. fMRI responses to pictures of mutilation and contamination. Neurosci. Lett. 393, 174-178. https://doi.org/10.1016/j.neulet.2005.09.072.

Schienle, A., Schäfer, A., Stark, R., Walter, B., Vaitl, D., 2005a. Relationship between disgust sensitivity, trait anxiety and brain activity during disgust induction. Neuropsychobiology 51, 86-92. https://doi.org/10.1159/000084165.

Schienle, A., Schäfer, A., Walter, B., Stark, R., Vaitl, D., 2005b. Brain activation of spider phobics towards disorder-relevant, generally disgust- and fear-inducing pictures. Neurosci. Lett. 388, 1-6. https://doi.org/10.1016/j.neulet.2005.06.025.

Schienle, A., Scharmüller, W., 2013. Cerebellar activity and connectivity during the experience of disgust and happiness. Neuroscience 246, 375-381. https://doi.org/10.1016/j.neuroscience.2013.04.048.

Schienle, A., Stark, R., Schäfer, A., Walter, B., Kirsch, P., Vaitl, D., 2004. Disgust and disgust sensitivity in bulimia nervosa: an fMRI study. Eur. Eat. Disorders Rev. 12, 42-50. https://doi.org/10.1002/erv.562.

Schienle, A., Stark, R., Walter, B., Blecker, C., Ott, U., Kirsch, P., Sammer, G., Vaitl, D., 2002. The insula is not specifically involved in disgust processing: an fMRI study. Neuroreport 13, 2023-2026. https://doi.org/10.1097/00001756-200211150-00006.

Schienle, A., Übel, S., Schöngaßner, F., Ille, R., Scharmüller, W., 2014. Disgust regulation via placebo: an fMRI study. Soc. Cogn. Affect. Neurosci. 9, 985-990. https://doi.org/10.1093/scan/nst072.

Schroeder, U., Hennenlotter, A., Erhard, P., Haslinger, B., Stahl, R., Lange, K., Ceballos-Baumann, A., 2005. Functional neuroanatomy of perceiving surprised faces. Hum. Brain Mapp. 23, 181-187. https://doi.org/10.1002/hbm.20057.

Shapira, N.A., Liu, Y., He, A.G., Bradley, M.M., Lessig, M.C., James, G.A., Stein, D.J., Lang, P.J., Goodman, W.K., 2003. Brain activation by disgust-inducing pictures in obsessive-compulsive disorder. Biol. Psychiatry 54, 751-756. https://doi.org/10.1016/S0006-3223(03)00003-9.

Shimamura, A.P., Marian, D.E., Haskins, A.L., 2013. Neural correlates of emotional regulation while viewing films. Brain Imaging Behav. 7, 77-84. https://doi.org/10.1007/s11682-012-9195-y.

Sprengelmeyer, R., Rausch, M., Eysel, U., Przuntek, H., 1998. Neural structures associated with recognition of facial expressions of basic emotions. Proc. R. Soc. Lond. B 265, 1927-1931. https://doi.org/10.1098/rspb.1998.0522.

Stark, R., Schienle, A., Girod, C., Walter, B., Kirsch, P., Blecker, C., Ott, U., Schäfer, A., Sammer, G., Zimmermann, M., Vaitl, D., 2005a. Erotic and disgust-inducing pictures - differences in the hemodynamic responses of the brain. Biol. Psychol. 70, 19-29. https://doi.org/10.1016/j.biopsycho.2004.11.014.

Stark, R., Schienle, A., Sarlo, M., Palomba, D., Walter, B., Vaitl, D., 2005b. Influences of disgust sensitivity on hemodynamic responses towards a disgust-inducing film clip. Int. J. Psychophysiol. 57, 61-67. https://doi.org/10.1016/j.ijpsycho.2005.01.010.

Stark, R., Schienle, A., Walter, B., Kirsch, P., Blecker, C., Ott, U., Schäfer, A., Sammer, G., Zimmermann, M., Vaitl, D., 2004. Hemodynamic effects of negative emotional pictures – a test-retest analysis. Neuropsychobiology 50, 108-118. https://doi.org/10.1159/000077948.

Stark, R., Zimmermann, M., Kagerer, S., Schienle, A., Walter, B., Weygandt, M., Vaitl, D., 2007. Hemodynamic brain correlates of disgust and fear ratings. Neuroimage 37, 663-673. https://doi.org/10.1016/j.neuroimage.2007.05.005.

Surguladze, S.A., Brammer, M.J., Young, A.W., Andrew, C., Travis, M.J., Williams, S.C.R., Phillips, M.L., 2003. A preferential increase in the extrastriate response to signals of danger. Neuroimage 19, 1317-1328. https://doi.org/10.1016/S1053-8119(03)00085-5.

Surguladze, S.A., El-Hage, W., Dalgleish, T., Radua, J., Gohier, B., Phillips, M.L., 2010. Depression is associated with increased sensitivity to signals of disgust: a functional magnetic resonance imaging study. J. Psychiatr. Res. 44, 894-902. https://doi.org/10.1016/j.jpsychires.2010.02.010.

Tettamanti, M., Rognoni, E., Cafiero, R., Costa, T., Galati, D., Perani, D., 2012. Distinct pathways of neural coupling for different basic emotions. Neuroimage 59, 1804-1817. https://doi.org/10.1016/j.neuroimage.2011.08.018.

Trautmann, S.A., Fehr, T., Herrmann, M., 2009. Emotions in motion: dynamic compared to static facial expressions of disgust and happiness reveal more widespread emotion-specific activations. Brain Res. 1284, 100-115. https://doi.org/10.1016/j.brainres.2009.05.075.

Viol, K., Aas, B., Kastinger, A., Kronbichler, M., Scholler, H., Reiter, E.-M., Said, S., Kronbichler, L., Kravanja-Spannberger, B., Stöger-Schmidinger, B., Aichhorn, W., Schiepek, G., 2019. Erroneously disgusted: fMRI study supports disgust-related neural reuse in obsessive-compulsive disorder (OCD). Front. Behav. Neurosci. 13, 81. https://doi.org/10.3389/fnbeh.2019.00081.

von dem Hagen, E.A.H., Beaver, J.D., Ewbank, M.P., Keane, J., Passamonti, L., Lawrence, A.D., Calder, A.J., 2009. Leaving a bad taste in your mouth but not in my insula. Soc. Cogn. Affect. Neurosci. 4, 379-386. https://doi.org/10.1093/scan/nsp018.

Wabnegger, A., Übel, S., Suchar, G., Schienle, A., 2018. Increased emotional reactivity to affective pictures in patients with skin-picking disorder: evidence from functional magnetic resonance imaging. Behav. Brain Res. 336, 151-155. https://doi.org/10.1016/j.bbr.2017.08.040.

Wicker, B., Keysers, C., Plailly, J., Royet, J.-P., Gallese, V., Rizzolatti, G., 2003. Both of us disgusted in my insula: the common neural basis of seeing and feeling disgust. Neuron 40, 655-664. https://doi.org/10.1016/S0896-6273(03)00679-2.

Williams, L.M., Das, P., Liddell, B., Olivieri, G., Peduto, A., Brammer, M.J., Gordon, E., 2005. BOLD, sweat and fears: fMRI and skin conductance distinguish facial fear signals. Neuroreport 16, 49-52. https://doi.org/10.1097/00001756-200501190-00012.

Wittfoth, D., Pfeiffer, A., Bohne, M., Lanfermann, H., Wittfoth, M., 2020. Emotion regulation through bifocal processing of fear inducing and disgust inducing stimuli. BMC Neurosci. 21, 47. https://doi.org/10.1186/s12868-020-00597-x.

Wright, P., He, G., Shapira, N., Goodman, W., Liu, Y., 2004. Disgust and the insula: fMRI responses to pictures of mutilation and contamination. Neuroreport 15, 2347-2351. https://doi.org/10.1097/00001756-200410250-00009.

Ziegler, J., Montant, M., Briesemeister, B., Brink, T., Wicker, B., Ponz, A., Bonnard, M., Jacobs, A., Braun, M., 2018. Do words stink? Neural reuse as a principle for understanding emotions in reading. J. Cogn. Neurosci. 30, 1023-1032. https://doi.org/10.1162/jocn_a_01268.
