## Supplementary Table 2 for "Common and distinct neurofunctional representations of core and social disgust in the brain: Coordinate-based and network meta-analyses"

Supplementary Table 2. Brain regions activated by conjunction analysis between core and social disgust processing in healthy subjects ($p < 0.05$ (uncorrected), minimum cluster size of 100 $mm^3$ and 5,000 permutations).

| Volume ($mm^3$) | Side | Brain region | Percentage of overlap | BA | MNI coordinates (x, y, z) | | | ALE value ($10^{-2}$) |
|---|---|---|---|---|---|---|---|---|
| 136 | right | Inferior Frontal Gyrus | 80% Inferior Frontal Gyrus, 20% Insula. | 47 | 44 | 26 | -8 | 1.49 |
| 16 | left | Fusiform Gyrus | 100% Fusiform Gyrus. | 37 | -40 | -52 | -16 | 1.21 |
| 16 | left | Fusiform Gyrus | 100% Fusiform Gyrus. | 37 | -42 | -50 | -14 | 1.37 |

Abbreviations: ALE = Action Likelihood Estimation, BA = Brodmann Area, MNI = Montreal Neurological Institute.
