## Supplementary Table 3 for "Common and distinct neurofunctional representations of core and social disgust in the brain: Coordinate-based and network meta-analyses"

Supplementary Table 3. Brain regions activated by contrast analysis between core and social disgust processing in healthy subjects ($p < 0.05$ (uncorrected), minimum cluster size of 100 $mm^3$ and 5,000 permutations).

| Volume ($mm^3$) | Side | Brain region | Percentage of overlap | BA | MNI coordinates (x, y, z) | | | Z |
|---|---|---|---|---|---|---|---|---|
| *Hyper-activation (core > social)* | | | | | | | | |
| 1272 | left | Postcentral Gyrus | 67.9% Inferior Parietal Lobule, 32.1% Postcentral Gyrus. | 2 | -66 | -20 | 34 | 2.67 |
| | left | Postcentral Gyrus | | 2 | -62 | -19 | 34 | 2.58 |
| | left | Postcentral Gyrus | | 2 | -60 | -22 | 38 | 2.56 |
| | left | Inferior Parietal Lobule | | 40 | -66.7 | -24 | 30.7 | 2.54 |
| | left | Inferior Parietal Lobule | | 40 | -64.7 | -27.3 | 35 | 2.42 |
| | left | Inferior Parietal Lobule | | 40 | -64 | -28 | 28 | 2.34 |
| 984 | left | Inferior Temporal Gyrus | 47.8% Middle Occipital Gyrus, 34.8% Inferior Temporal Gyrus, 8.7% Inferior Occipital Gyrus, 8.7% Middle Temporal Gyrus. | 37 | -51 | -74 | 0 | 3.16 |
| | left | Middle Occipital Gyrus | | 37 | -44 | -72 | 6 | 2.56 |
| 848 | left | Parahippocampal Gyrus | 50.1% Brodmann area 34, 44.8% Amygdala, 3.4% Globus Pallidus, 1.7% Brodmann area 28. | 34 | -16 | 0 | -22 | 2.20 |
| | left | Parahippocampal Gyrus | | 34 | -18 | 0 | -14 | 1.93 |
| 672 | right | Superior Frontal Gyrus | 73.7% Superior | 9 | 4 | 60 | 21 | 2.46 |

| | | | | | | | | |
|---|---|---|---|---|---|---|---|---|
| | right | Superior Frontal Gyrus | Frontal Gyrus, 26.3% Medial | 9 | 4 | 64 | 16 | 2.28 |
| | left | Medial Frontal Gyrus | Frontal Gyrus. | 10 | 0 | 64 | 14 | 2.24 |
| 304 | left | Inferior Frontal Gyrus | 100% Inferior Frontal Gyrus. | 47 | -28 | 34 | -10 | 2.20 |
| | left | Claustrum | | | -28 | 30 | -8 | 2.12 |
| 248 | left | Fusiform Gyrus | 69.2% Declive, | 37 | -40 | -66 | -12 | 2.14 |
| | left | Declive | 30.8% Fusiform Gyrus. | | -32 | -70 | -10 | 1.84 |
| 216 | left | Lingual Gyrus | 92.3% Lingual Gyrus, 7.7% Inferior Occipital Gyrus. | 17 | -8 | -100 | -6 | 1.91 |
| *Hyper-activation (social > core)* | | | | | | | | |
| 1336 | right | Insula | 71.4% Insula, 20.0% Inferior | 13 | 50 | 16 | 10 | 3.72 |
| | right | Inferior Frontal Gyrus | Frontal Gyrus, 8.6% | 44 | 50 | 20 | 8 | 3.24 |
| | right | Inferior Frontal Gyrus | Precentral Gyrus. | 44 | 46 | 20.7 | 7.3 | 3.16 |
| 1160 | left | Medial Frontal Gyrus | 57.5% Medial Frontal Gyrus, | 32 | -2 | 14 | 48 | 3.54 |
| | right | Cingulate Gyrus | 31.3% Superior Frontal Gyrus, 11.3% Cingulate Gyrus. | 24 | 4 | 10 | 44 | 3.09 |
| 1064 | right | Middle Occipital Gyrus | 100% Middle Occipital | | 30 | -90 | 2 | 3.35 |
| | right | Middle Occipital Gyrus | Gyrus. | 19 | 38 | -88 | 12 | 3.24 |
| 1048 | right | Declive | 81.3% | | 42 | -76 | -16 | 3.72 |
| | right | Declive | Declive, 18.7% Fusiform Gyrus. | | 41 | -70 | -18 | 3.24 |

| | | | | | | | | |
|---|---|---|---|---|---|---|---|---|
| 728 | right | Superior Temporal Gyrus | 55.6% Middle Temporal Gyrus, 44.4% Superior Temporal Gyrus. | 22 | 53.8 | -34.2 | 2 | 3.54 |
| 472 | left | Fusiform Gyrus | 78.9% Fusiform Gyrus, 21.1% Culmen. | 37 | -36 | -46 | -16 | 2.58 |

Abbreviations: ALE = Action Likelihood Estimation, BA = Brodmann Area, MNI = Montreal Neurological Institute.
