## Supplementary Table 4 for "Common and distinct neurofunctional representations of core and social disgust in the brain: Coordinate-based and network meta-analyses"

Supplementary Table 4. Network of right inferior frontal gyrus revealed by MACM-B analysis.

| Volume (mm$^3$) | Side | Brain region | BA | MNI coordinates (x, y, z) | | | ALE value | Z |
|---|---|---|---|---|---|---|---|---|
| 123176 | right | Extra-Nuclear | 47 | 40 | 24 | -8 | 0.64 | 37.09 |
| | left | Insula | | -34 | 22 | -2 | 0.35 | 20.53 |
| | right | Thalamus | | 10 | -14 | 4 | 0.19 | 11.84 |
| | left | Thalamus | | -10 | -16 | 4 | 0.18 | 11.11 |
| | left | Inferior Frontal Gyrus | 9 | -46 | 8 | 28 | 0.14 | 9.11 |
| | right | Inferior Frontal Gyrus | 9 | 48 | 10 | 26 | 0.13 | 8.11 |
| | left | Middle Frontal Gyrus | 6 | -42 | 2 | 48 | 0.13 | 8.08 |
| | right | Middle Frontal Gyrus | 6 | 50 | 6 | 42 | 0.13 | 7.94 |
| | left | Middle Frontal Gyrus | 46 | -48 | 26 | 16 | 0.12 | 7.67 |
| | right | Caudate | | 12 | 8 | 4 | 0.12 | 7.58 |
| | right | Middle Frontal Gyrus | 9 | 50 | 22 | 28 | 0.12 | 7.32 |
| | left | Putamen | | -14 | 6 | 0 | 0.10 | 6.52 |
| | right | Middle Frontal Gyrus | 46 | 52 | 32 | 20 | 0.10 | 6.50 |
| | right | Superior Frontal Gyrus | 9 | 40 | 44 | 22 | 0.10 | 6.30 |
| | right | Middle Frontal Gyrus | 10 | 44 | 42 | 20 | 0.10 | 6.26 |
| | left | Putamen | | -22 | 0 | 4 | 0.09 | 5.90 |
| | left | Precentral Gyrus | 6 | -30 | -2 | 52 | 0.09 | 5.36 |
| | right | Amygdala | | 20 | -4 | -16 | 0.09 | 5.34 |
| | right | Inferior Frontal Gyrus | 45 | 54 | 20 | 12 | 0.08 | 5.12 |
| | right | Inferior Frontal Gyrus | | 44 | 50 | -6 | 0.08 | 4.74 |
| | right | Middle Frontal Gyrus | 6 | 42 | 0 | 54 | 0.08 | 4.52 |
| | left | Amygdala | | -20 | -6 | -14 | 0.08 | 4.52 |
| | right | Thalamus | | 20 | -30 | -4 | 0.07 | 4.37 |

| | | | | | | | | |
|---|---|---|---|---|---|---|---|---|
| | left | Middle Frontal Gyrus | 9 | -40 | 38 | 22 | 0.06 | 3.39 |
| | left | Thalamus | | -18 | -30 | -4 | 0.06 | 3.39 |
| 28056 | left | Superior Frontal Gyrus | 6 | 2 | 12 | 52 | 0.23 | 14.04 |
| | left | Medial Frontal Gyrus | 8 | 0 | 26 | 42 | 0.22 | 13.57 |
| 10896 | left | Inferior Parietal Lobule | 40 | -34 | -54 | 48 | 0.15 | 9.60 |
| | left | Superior Parietal Lobule | 7 | -26 | -64 | 50 | 0.10 | 6.51 |
| | left | Inferior Parietal Lobule | 40 | -42 | -44 | 44 | 0.09 | 5.51 |
| | left | Precuneus | 19 | -26 | -72 | 38 | 0.07 | 4.11 |
| 6776 | left | Fusiform Gyrus | 37 | -42 | -58 | -16 | 0.12 | 7.90 |
| 6504 | right | Precuneus | 19 | 34 | -60 | 46 | 0.11 | 6.86 |
| | right | Inferior Parietal Lobule | 40 | 50 | -38 | 48 | 0.09 | 5.54 |
| | right | Inferior Parietal Lobule | 40 | 42 | -42 | 46 | 0.09 | 5.42 |
| 2864 | left | Middle Temporal Gyrus | 22 | -56 | -42 | 4 | 0.10 | 6.45 |
| | left | Superior Temporal Gyrus | 13 | -54 | -40 | 22 | 0.07 | 4.01 |
| 2184 | right | Superior Temporal Gyrus | 22 | 54 | -34 | 2 | 0.09 | 5.43 |
| 1896 | left | Superior Temporal Gyrus | | -60 | -20 | 2 | 0.08 | 4.66 |
| | left | Transverse Temporal Gyrus | 41 | -58 | -20 | 12 | 0.07 | 4.17 |
| | left | Postcentral Gyrus | 40 | -60 | -20 | 20 | 0.07 | 3.87 |

ALE = Action Likelihood Estimation, BA = Brodmann Area, MNI = Montreal Neurological Institute.
