## Supplementary Table 5 for "Common and distinct neurofunctional representations of core and social disgust in the brain: Coordinate-based and network meta-analyses"

Supplementary Table 5. Network of left fusiform gyrus revealed by MACM-B analysis.

| Volume (mm³) | Side | Brain region | BA | MNI coordinates (x, y, z) | | | ALE value | Z |
|---|---|---|---|---|---|---|---|---|
| 42864 | left | Fusiform Gyrus | 37 | -42 | -52 | -16 | 0.54 | 32.08 |
| | left | Middle Occipital Gyrus | 37 | -46 | -68 | -4 | 0.15 | 9.99 |
| | left | Fusiform Gyrus | 19 | -42 | -76 | -10 | 0.13 | 9.01 |
| | left | Inferior Occipital Gyrus | 18 | -28 | -94 | -2 | 0.11 | 7.83 |
| | left | Middle Occipital Gyrus | 18 | -32 | -88 | 6 | 0.11 | 7.59 |
| | left | Middle Temporal Gyrus | 22 | -60 | -44 | 4 | 0.10 | 6.71 |
| | left | Middle Temporal Gyrus | 22 | -62 | -34 | 4 | 0.08 | 5.39 |
| | left | Superior Temporal Gyrus | 22 | -58 | -24 | 2 | 0.06 | 3.89 |
| | left | Lingual Gyrus | 17 | -14 | -98 | -2 | 0.05 | 3.50 |
| 39472 | left | Inferior Frontal Gyrus | 9 | -44 | 8 | 28 | 0.21 | 13.93 |
| | left | Middle Frontal Gyrus | 46 | -48 | 26 | 16 | 0.16 | 10.68 |
| | left | Claustrum | | -32 | 22 | 0 | 0.13 | 8.82 |
| | left | Inferior Frontal Gyrus | 47 | -48 | 28 | -4 | 0.11 | 7.69 |
| | left | Precentral Gyrus | 44 | -52 | 12 | 6 | 0.08 | 5.46 |
| 30888 | right | Culmen | | 40 | -50 | -20 | 0.22 | 14.26 |
| | right | Inferior Occipital Gyrus | 19 | 42 | -78 | -6 | 0.13 | 8.85 |
| | right | Fusiform Gyrus | 19 | 44 | -70 | -10 | 0.12 | 8.01 |
| | right | Middle Occipital Gyrus | 18 | 26 | -90 | -4 | 0.11 | 7.43 |
| | right | Middle Occipital Gyrus | 18 | 34 | -86 | 4 | 0.09 | 6.45 |
| | right | Declive | | 28 | -68 | -12 | 0.07 | 4.83 |
| | right | Middle Temporal Gyrus | 39 | 50 | -58 | 10 | 0.06 | 3.92 |
| 20072 | right | Inferior Frontal Gyrus | 9 | 48 | 12 | 28 | 0.12 | 8.59 |
| | right | Insula | | 36 | 22 | -6 | 0.12 | 8.32 |

| | | | | | | | | |
|---|---|---|---|---|---|---|---|---|
| | right | Middle Frontal Gyrus | 46 | 50 | 28 | 20 | 0.09 | 6.35 |
| | right | Inferior Frontal Gyrus | 13 | 44 | 30 | 8 | 0.08 | 5.90 |
| | right | Inferior Frontal Gyrus | 13 | 48 | 28 | 8 | 0.08 | 5.79 |
| | right | Insula | | 48 | 20 | -4 | 0.06 | 4.26 |
| | right | Putamen | | 24 | 8 | 2 | 0.06 | 4.22 |
| | right | Middle Frontal Gyrus | 6 | 46 | 2 | 48 | 0.06 | 3.51 |
| 12000 | left | Superior Frontal Gyrus | 6 | -2 | 14 | 50 | 0.17 | 11.44 |
| | right | Cingulate Gyrus | 32 | 4 | 26 | 38 | 0.07 | 5.13 |
| 11592 | left | Inferior Parietal Lobule | 40 | -34 | -52 | 48 | 0.12 | 8.30 |
| | left | Superior Parietal Lobule | 7 | -28 | -60 | 48 | 0.11 | 7.76 |
| | left | Inferior Parietal Lobule | 40 | -44 | -36 | 46 | 0.09 | 6.31 |
| 4688 | left | Amygdala | | -20 | -8 | -16 | 0.13 | 8.89 |
| | left | Putamen | | -22 | 6 | 4 | 0.06 | 4.20 |
| 4608 | right | Superior Parietal Lobule | 7 | 32 | -54 | 48 | 0.13 | 8.71 |
| 1944 | right | Amygdala | | 22 | -6 | -18 | 0.11 | 7.57 |

ALE = Action Likelihood Estimation, BA = Brodmann Area, MNI = Montreal Neurological Institute.
